# On the variation of structural divergence among residues in enzyme evolution

**DOI:** 10.1101/2024.12.23.629899

**Authors:** Julian Echave, Mathilde Carpentier

**Affiliations:** Instituto de Ciencias Físicas (ICIFI-CONICET), Universidad Nacional de San Martín, Martín de Irigoyen 3100, 1650 San Martín, Buenos Aires, Argentina; Institut de Systématique, Evolution, Biodiversité (ISYEB, UMR 7205), CNRS-MNHN-SU-EPHE-UA, Sorbonne Université, 57 rue Cuvier, 75005 Paris, France

**Keywords:** Enzyme structural evolution, Residue-specific divergence, Evolutionary constraints, Evolution-dynamics relationship, Structure-function relationship, Active site conservation

## Abstract

Structural divergence varies among protein residues, yet this variation has been largely overlooked compared with the well-studied case of sequence rate variation. Here we show that, in families of functionally conserved homologous enzymes, structural divergence increases with both residue flexibility and distance from the active site. Although these properties are correlated, modelling reveals that the pattern arises from two independent types of evolutionary constraints: non-functional and functional. The balance between these constraints varies widely across enzyme families, from non-functional to functional dominance. As functional constraints strengthen, structural divergence patterns are reshaped, becoming increasingly distinct from flexibility patterns and breaking the commonly assumed correspondence between evolutionary and dynamical structural ensembles. Active sites are more structurally conserved than average, but this conservation stems not only from functional constraints. Because active sites typically lie in rigid regions where non-functional constraints are high, both constraint types contribute comparably on average, with dominance shifting from one to the other depending on active-site rigidity. Together, these findings revise two long-standing assumptions: that evolutionary structural variation universally mirrors protein dynamics, and that active-site conservation reflects functional requirements alone. Both depend on the balance between non-functional and functional constraints that shape enzyme structural evolution.

## 1 Introduction

Protein structures evolve more slowly than sequences, a long-recognized property that enables comparisons between distantly related proteins whose sequences have diverged beyond recognition [1, 2, 3, 4]. This property makes structural data particularly valuable for structural phylogenetics, providing an alternative for studying evolutionary relationships when sequences can no longer provide meaningful comparisons [5, 6, 7, 8, 9, 10, 11, 12, 13, 14, 15, 16, 17]. Despite their potential, structures have remained underused due to the limited availability of structural data compared to sequences. However, machine learning methods have substantially improved both the speed and accuracy of protein structure prediction, making structural data more accessible [18, 19, 20, 21, 22]. The use of structural data in structuralphylogenetics and other evolutionary studies is now limited mostly by our incomplete understanding of protein structural evolution.

Most studies of structural evolution focus on whole-protein structural divergence [1, 2, 3, 4]. These studies typically measure structural similarity using global metrics such as RMSD, conservation of secondary structures, contacts, or solvent accessibility. They have shown that structural divergence increases with sequence divergence (either linearly or exponentially depending on the measure used) and that structures generally evolve more slowly than sequences. However, they do not examine how structural divergence varies within proteins, which limits our current understanding.

The variation of amino acid substitution rates among residues is one of the classic problems of protein sequence evolution (reviewed in Echave et al. [23]; see also Nagar et al. [24] and references therein). Through iterative cycles of pattern discovery and mechanistic explanation, this problem has revealed the major determinants of substitution rate variation and their underlying causes, providing fundamental insights into the evolutionary process. In contrast, the analogous problem in structure evolution—the variation of structural divergence among residues—has received little attention. Here, we examine how evolutionary constraints determine the variation of structural divergence among residues within enzymes under neutral evolution with conserved function.

To anticipate key issues that we will develop in this work, we introduce class A beta lactamases as an illustrative case of how structural divergence varies among residues. Their residue-level structural divergence (Figure 1a) shows a conserved core surrounded by variable regions. The variation of structural divergence correlates with both conformational flexibility (Figure 1b) and distance from the active site (Figure 1c). As we will discuss next, previous work on protein evolution suggests that these correlations reflect two types of constraints in functionally conserved homologues: non-functional constraints and functional constraints. Non-functional constraints lead to an increase in divergence with flexibility, while functional constraints lead to an increase in divergence with distance from the active site. However, because flexibility and distance are correlated, disentangling the respective contributions of these constraints requires careful analysis. Below, we examine these issues in detail to understand their roles in shaping structural divergence.

**Figure 1:**
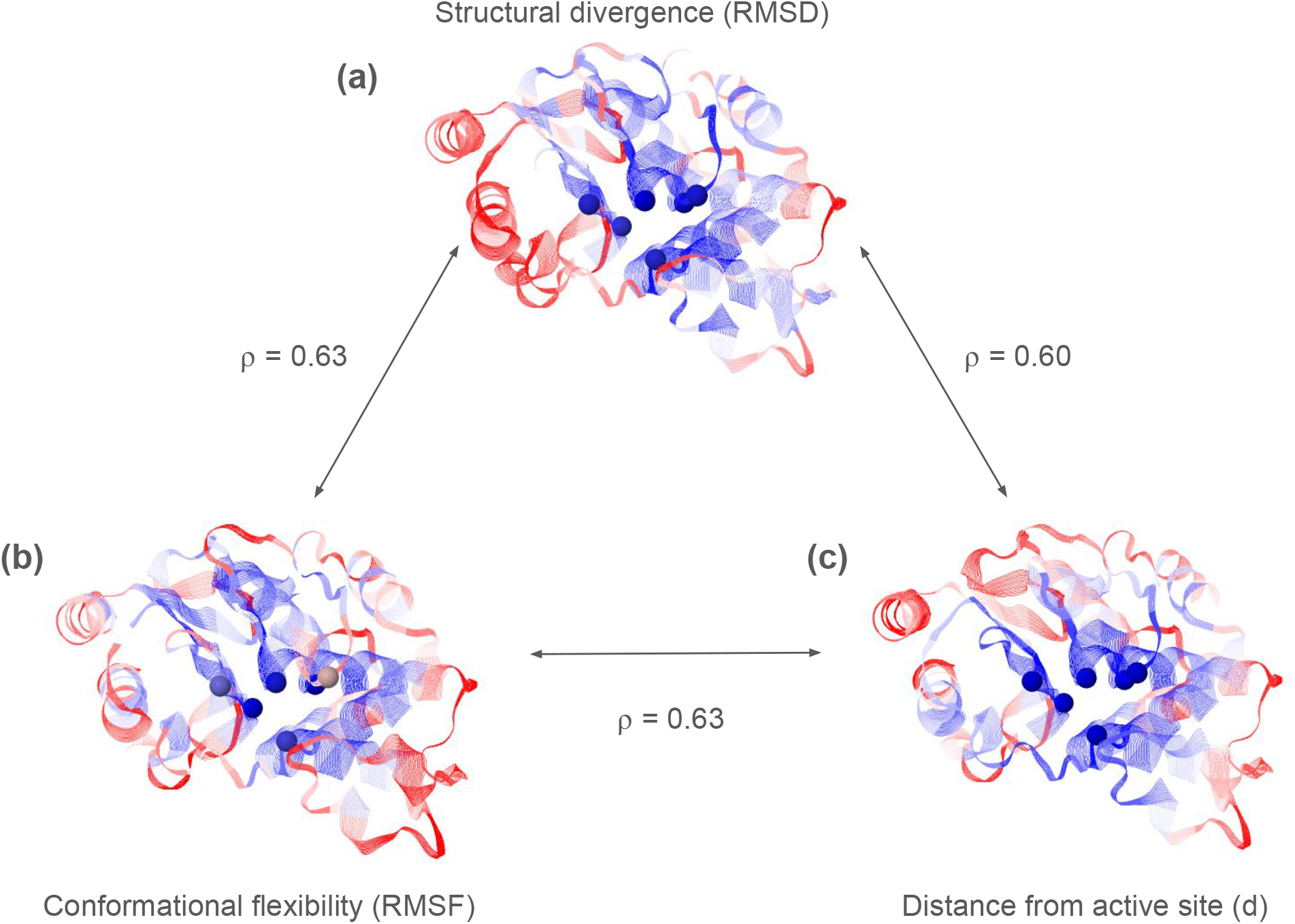
Patterns of residue-level structural divergence, flexibility, and distance to the active site in class A beta-lactamases (M-CSA ID = 2). **a**: Structural divergence at each residue position, quantified by the root mean square deviation (`RMSD`) of residue coordinates across homologous structures. **b**: Local flexibility of each residue, measured by root mean square fluctuation (`RMSF`) in the reference enzyme during protein dynamics. **c**: Each residue’s distance in the reference enzyme to the closest active-site residue (*d*). Spearman correlation coefficients between these patterns are displayed next to the double-headed arrows connecting the panels. All measurements use *C*_*α*_ coordinates. Values are shown as ranks mapped onto the 3D structure of the reference enzyme (PDB code: 1btl), coloured from blue (low) to red (high). Active-site *C*_*α*_ atoms are shown as spheres.

Under non-functional constraints - arising from mutational effects and stability requirements - structural divergence increases with flexibility. The set of structures that emerge through evolution from a common ancestor (the evolutionary ensemble) resembles the set of conformations sampled by an individual protein during dynamics (the dynamical ensemble) [25, 26, 27, 28, 29, 30, 31, 32, 33, 34, 35] This similarity is reflected in the correlation between measures that characterize each ensemble: residue structural divergence (`RMSD`) across homologues correlates with residue flexibility (`RMSF`) during dynamics (Figure 1) [34]. Biophysical models show this similarity arises from two non-functional constraints: mutational sensitivity, whereby flexible residues accumulate larger structural displacements because they respond more strongly to perturbations, and selection on stability, which concentrates substitutions in flexible regions where mutations are better tolerated, further amplifying their divergence [30, 34].

In enzymes, we expect functional constraints to lead to increasing structural divergence with distance from the active site. Active sites are structurally conserved among homologous enzymes that maintain their ancestral function, suggesting the presence of functional constraints [36, 37]. Studies of sequence evolution show that amino acid substitution rates increase gradually with distance from the active site [38], a pattern explained by the gradual decay of functional constraints with distance [39]. Because mutations tend to perturb local structure [40, 41], this gradual decay of functional constraints should also be reflected in structural divergence patterns.

Because of their interdependence, the observed divergence-flexibility and divergence-distance trends cannot be directly attributed to purely non-functional or functional constraints, as each may reflect contributions from both. The observed divergence-flexibility trend could arise from a direct effect of non-functional constraints, or an indirect effect of functional constraints, via the positive flexibility-distance correlation, or both. Similarly, the observed divergence-distance trend could stem from a direct effect of functional constraints, an indirect effect of non-functional constraints via the flexibility-distance correlation, or both. Further analysis is needed to disentangle the independent contributions of functional and non-functional constraints to structural divergence.

Here we analyse how structural divergence varies among residues within enzymes under neutral evolution with conserved function. To determine how functional and non-functional constraints influence structural divergence, we examine how it relates to residue flexibility and distance from the active site, accounting for their interdependence. Contrasting scenarios of adaptive evolution, where active sites undergo structural changes to accommodate novel functions, will be addressed in the discussion section.

## 2 Results

To investigate how structural divergence varies under neutral evolution with conserved function, we curated a dataset of families of functionally conserved enzymes. Starting from the Mechanism and Catalytic Site Atlas (M-CSA) database [42, 43], we selected entries for which the reference enzyme, the one for which the catalytic mechanism has been determined, is monomeric and single-domain, to avoid additional constraints from domain or chain interactions. We then filtered for homologues likely to share the reference enzyme’s function, keeping only those with identical active-site residues, >25% sequence identity, and the same CATH classification. Additional filtering steps ensured data quality and representative sampling by removing redundant structures and potential outliers. This approach yielded 34 families representing a diverse range of structures and functions. (Detailed dataset properties are provided in Section 4.1.)

Next, for each family, we calculated three measures at each residue position (illustrated in Figure1): (1) root mean square deviation(`RMSD`), using the *C*_*α*_ coordinates of aligned and superimposed homologous enzymes, quantifying residue-level structural divergence; (2) root mean square fluctuation(`RMSF`), using a simple *C*_*α*_-based coarse-grained elastic network model of the reference enzyme, quantifying the flexibility of individual residues; (3) distance from the active site (*d*), defined as the distance between a residue’s *C*_*α*_ atom and the *C*_*α*_ atom of the nearest active-site residue, quantifying its proximity to the catalytic region.

Finally, we analysed how structural divergence (`RMSD`) relates to flexibility (`RMSF`) and distance from the active site (*d*).

Given the wide variation in structural divergence between conserved and variable regions, we primarily use logarithmic transformations (`lRMSD`, `lRMSF`) for their better statistical properties for modelling. We also employ normalized versions (`nlRMSD`, `nlRMSF`) that express residue-level divergence and flexibility relative to the protein average (see Section 4 for detailed definitions). We specify whether we use the raw, logarithmic, or normalized-logarithmic versions of `RMSD` and `RMSF` only when necessary. In contexts where the precise metric does not affect the interpretation, such as general discussions of trends (e.g., ‘divergence increases with flexibility’), we may refer to these variables without explicitly specifying the measure used.

### 2.1 Variation of structural divergence among residues

Structural divergence varies widely among residues. Specifically, `RMSD` differs markedly between conserved and variable residues. In class A beta-lactamases, the Q90/Q10 `RMSD` ratio, calculated using the 90th and 10th percentiles, is 6.6 (Figure 2a); the evolutionary deformation of variable regions is almost 7 times faster than that of conserved regions. This wide variation is consistent across all enzyme families: Q90/Q10 `RMSD` ratios range from 3.6 to 14.0 (mean = 7.7 ± 2.5; ± denotes SD; Figure 2d; and Figures S6a to S39a).

**Figure 2:**
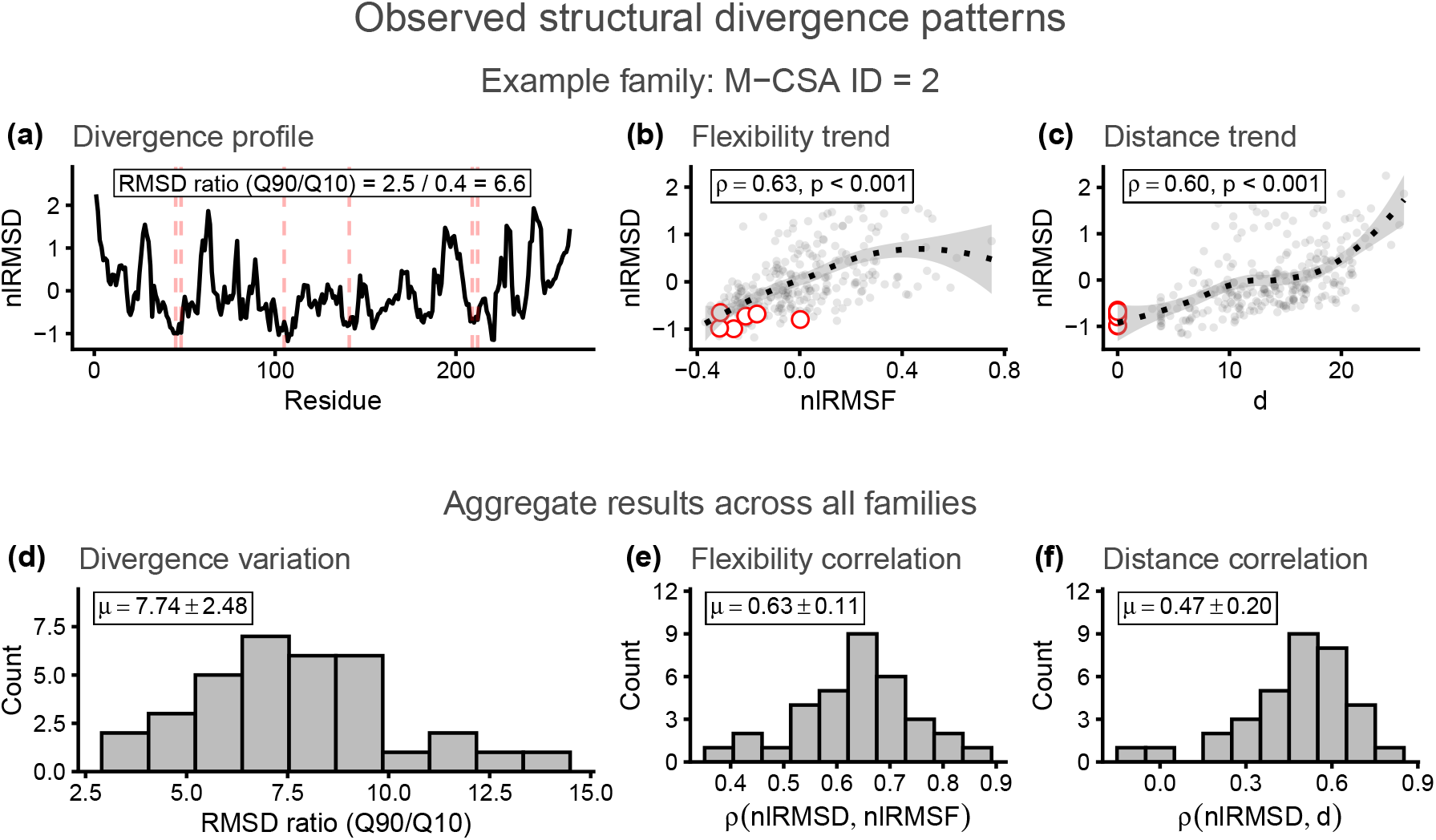
Observed structural divergence patterns. **a**: Residue-specific structural divergence (`nlRMSD`) vs. residue number for class A beta-lactamases (M-CSA ID = 2), with ratio of 90th to 10th percentile `RMSD` values shown. Active-site residues are indicated by vertical dashed red lines. **b**: `nlRMSD` vs. residue flexibility (`nlRMSF`) for the same family. **c**: `nlRMSD` vs. distance to the closest active-site residue (*d*) for the same family. In **b** and **c**, grey points represent individual residues, active-site residues are marked in red, and the black dotted line represents a LOESS fit to the data points; *ρ* is Spearman’s correlation coefficient, with corresponding p-values provided. **d**: Distribution of Q90/Q10 `RMSD` ratios across all enzyme families. **e**,**f**: Distributions of Spearman’s correlation coefficients across all families between `nlRMSD` and: **e**) `nlRMSF`; **f**) *d. µ* indicates the mean ± standard deviation for each distribution.

To investigate the determinants of this variation, we analysed the dependence of `RMSD` on two properties: residue flexibility (`RMSF`) and distance to the active site (*d*) .

Structural divergence increases with flexibility. As shown in Figure 2b, for class A beta-lactamases, residue RMSD increases with residue `RMSF`, with a Spearman’s correlation coefficient *ρ* = 0.63 (p < 0.001). This trend is observed in all enzyme families, with *ρ*(`RMSD`, `RMSF`) ranging from 0.36 to 0.85 (all p < 0.001,) and averaging 0.63 ± 0.11(Figure 2e; Figures S6b to S39b).

Structural divergence increases with distance to the active site in most enzyme families. In class A beta-lactamases, residue `RMSD` increases with *d* (*ρ* = 0.60, p < 0.001; Figure 2c). This trend is observed in most enzyme families, with *ρ*(`RMSD`, *d*) averaging 0.47 ± 0.20. Of the 34 families studied, 32 show positive correlations ranging from 0.16 to 0.78 (Figure 2f; Figures S6c to S39c). Only two families deviate from this pattern: M-CSA ID 749 shows a slight negative correlation (*ρ* = −0.12, p = 0.03; Figure S33c) and M-CSA ID 258 shows no correlation (*ρ* = 0; Figure S19c). (These cases will be discussed in Section 3.5.)

If flexibility (`RMSF`) and distance (*d*) were independent variables, the observed divergence-flexibility dependence could be attributed to non-functional constraints, and the observed divergence-distance dependence to functional constraints. We could then conclude that both constraints are influencing the observed patterns of variation of structural divergence within enzymes.

However, flexibility and distance are not independent. There is a positive correlation between `RMSF` and *d* across enzyme families, with an average *ρ* = 0.5 ± 0.2. This correlation arises because active sites in most enzymes are located in rigid regions, where flexibility is low, so that flexibility tends to increase with distance from the active site. The strength of this effect depends on active site rigidity, as evidenced by a strong negative correlation (*R* = − 0.84) between active site flexibility and *ρ*(`RMSF`, *d*) (Figure S2b). Most families exhibit positive correlations ranging from 0.16 to 0.85 (Figure S2a), but in cases like M-CSA ID 362, where the active site resides in a flexible region, flexibility and distance are nearly uncorrelated (*ρ* = −0.04, p = 0.7), confirming that the correlation depends on active site location.

Because of this interdependence, the observed divergence-flexibility and divergence-distance trends cannot be directly attributed to purely non-functional or functional constraints, as each may reflect contributions from both. The observed divergence-flexibility trend could arise from a direct effect of non-functional constraints, or an indirect effect of functional constraints, via the positive flexibility-distance correlation, or both. Similarly, the observed divergence-distance trend could arise from a direct effect of functional constraints, an indirect effect of non-functional constraints via the flexibility-distance correlation, or both. Further analysis is needed to disentangle the independent contributions of functional and non-functional constraints to structural divergence.

### 2.2 Independent contributions of non-functional and functional constraints

#### Three alternative models of structural divergence

In the previous section, we established that structural divergence varies widely among residues, increasing with both flexibility and distance from the active site. To disentangle the contributions of evolutionary constraints to these patterns, we formulate three mathematical models (Figure 3). The first model (*M*_1_) represents a scenario where only non-functional constraints operate, with a direct effect making structural divergence increase with flexibility (`lRMSD` = *s*_1_(`lRMSF`)). Due to the correlation between flexibility and distance, *M*_1_ would also predict an indirect increase of divergence with distance, even without functional constraints. The second model (*M*_2_) represents a scenario where only functional constraints operate, with a direct effect making structural divergence increase with distance (`lRMSD` = *s*_2_(*d*)). Due to the correlation between flexibility and distance, *M*_2_ would also predict an indirect increase of divergence with flexibility, even without non-functional constraints. The third model (*M*_12_) combines both direct effects (`lRMSD` = *s*_1_(`lRMSF`) + *s*_2_(*d*)), allowing both types of constraints to operate simultaneously. Since *a priori* any of these three models could explain the observed increases in structural divergence with both flexibility and distance, we need to fit and compare them to determine which better describes the data.

**Figure 3:**
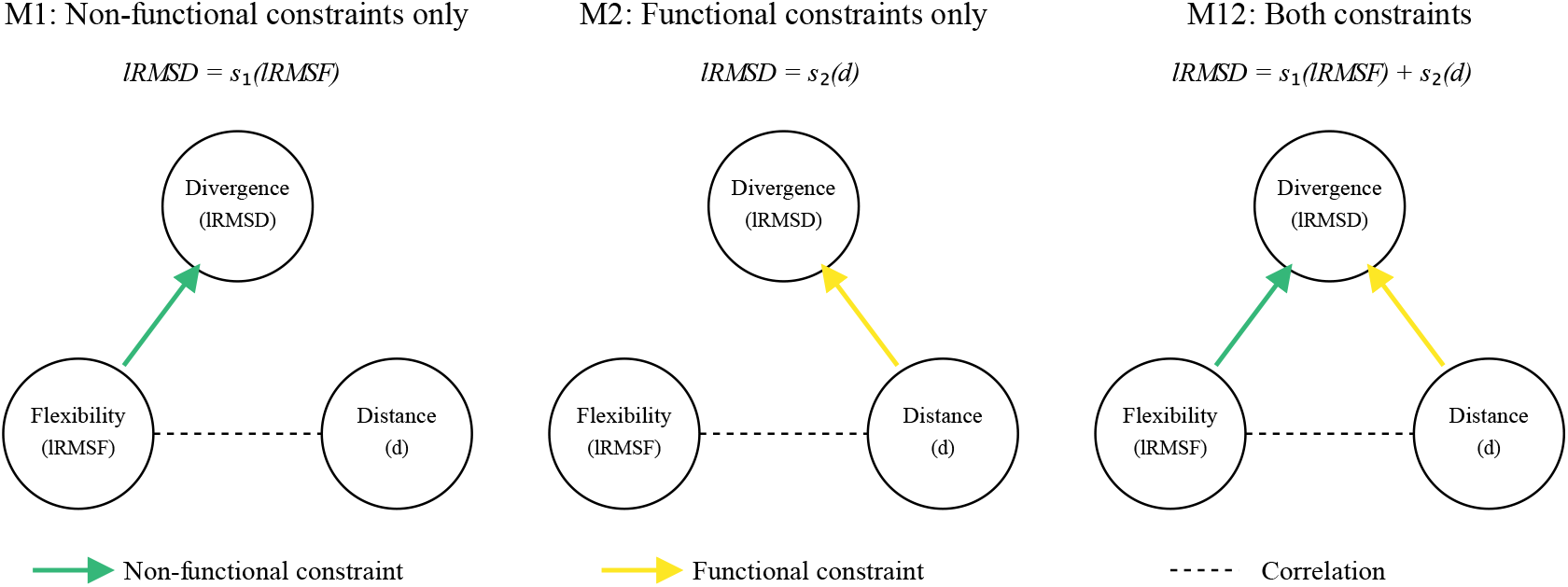
Alternative models for how evolutionary constraints influence structural divergence. Three models represent different scenarios for how structural divergence (`lRMSD`) depends on flexibility (`lRMSF`) and distance from active site (*d*). Green arrows represent direct effects of non-functional constraints (function *s*_1_), yellow arrows represent direct effects of functional constraints (function *s*_2_), and dashed lines indicate correlation between variables. In *M*_1_, only non-functional constraints act; in *M*_2_, only functional constraints act; in *M*_12_, both constraint types operate. The correlation between variables means each single-constraint model predicts relationships with both variables through direct and indirect effects.

We implemented these models using shape-constrained additive models (SCAMs), which are a variant of generalized additive models (GAMs) that incorporate specific shape constraints. In our case, these shape constraints enforce monotone-increasing functions to reflect expected direct effects of evolutionary constraints: structural divergence is expected to increase with flexibility due to weakening non-functional constraints and with distance due to weakening functional constraints.

#### Predicted vs. observed residue structural divergence

We fit the models *M*_1_, *M*_2_, and *M*_12_ to residue `lRMSD` values for each enzyme family separately. Model selection relied on the Akaike information criterion (AIC), which balances model complexity and goodness of fit. To evaluate model performance, we used explained deviance, a measure analogous to explained variance in linear models but well-suited for non-linear relationships.

The findings demonstrate that functional and non-functional constraints make independent contributions to the variation in structural divergence among residues. In class A beta-lactamases (Figure 4a), *M*_12_ is the best model according to AIC, with values of 447.7, 454.9, and 410.7 for *M*_1_, *M*_2_, and *M*_12_, respectively. (The best model is the one with lowest AIC, and AIC differences greater than two are considered significant [44, 45].) Explained deviance values—0.36 for *M*_1_, 0.35 for *M*_2_, and 0.46 for *M*_12_—further confirm that *M*_12_ outperforms the single-constraint-type models, indicating that both *s*_1_(`lRMSF`) and *s*_2_(*d*) contribute independently to `lRMSD`.

**Figure 4:**
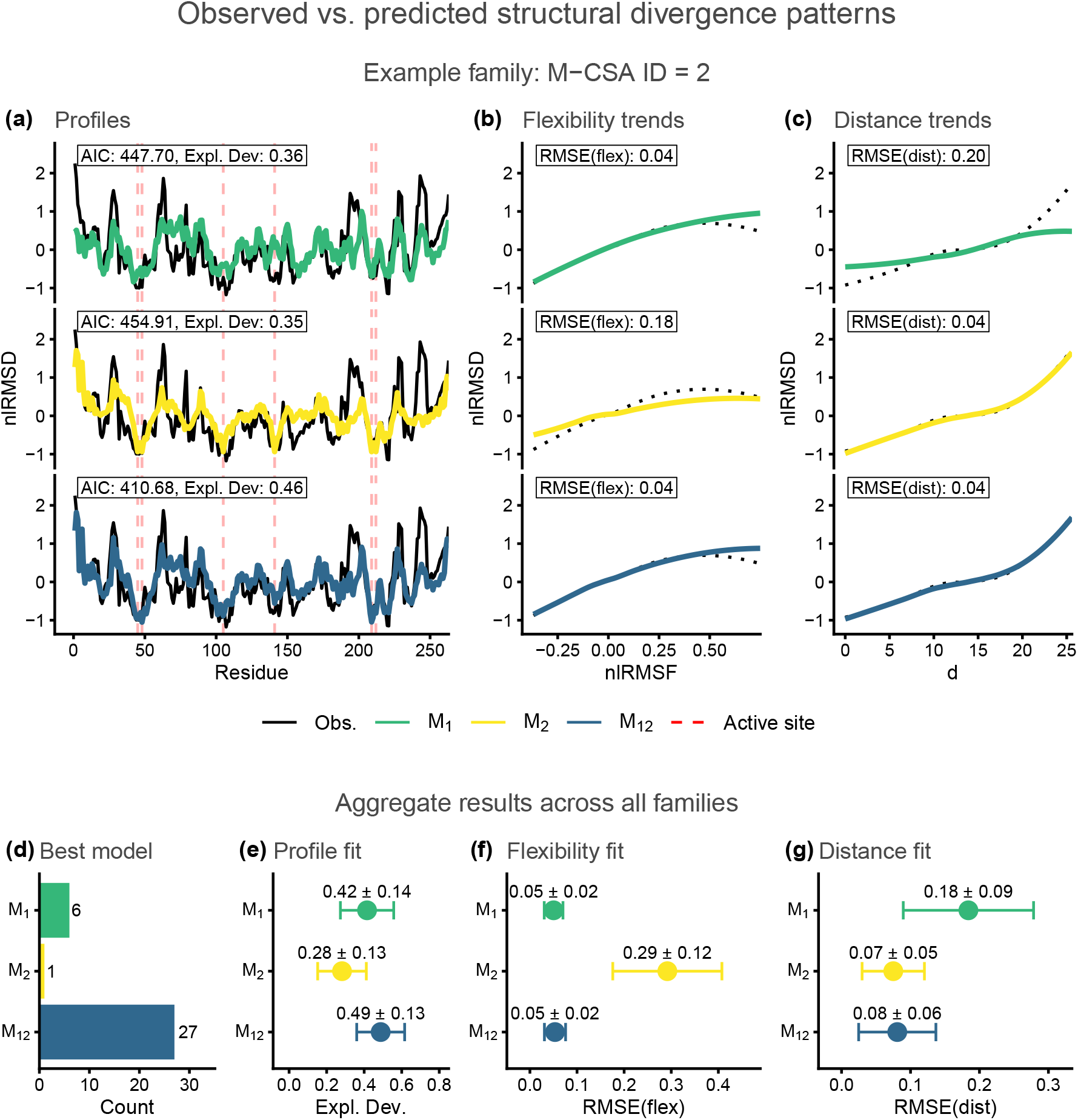
Observed vs. predicted structural divergence patterns. **a**: Residue-specific `nlRMSD` profiles for class A beta-lactamases (M-CSA ID = 2), showing observed values (black) and predictions from three models: *M*_1_ (green, reflecting non-functional constraints only, *s*_1_), *M*_2_ (yellow, reflecting functional constraints only, *s*_2_), and *M*_12_ (blue, reflecting both constraints combined). Active-site residues are indicated by vertical dashed red lines. AIC and explained deviance values for each model are shown. **b**: Divergence-flexibility trends for class A beta-lactamases. **c**: Divergence- distance trends for class A beta-lactamases. In **b** and **c**, black dotted lines show LOESS fits to observed data, coloured lines show LOESS fits to model predictions, and RMSE values quantify the differences between these trends. **d**: Number of enzyme families (bar chart) for which each model is optimal (lowest AIC). **e**: Explained deviance for each model across enzyme families. **f**: RMSE(flex) across enzyme families, measuring fit to divergence-flexibility trends. **g**: RMSE(dist) across enzyme families, measuring fit to divergence-distance trends. In **e**–**g**, points and error bars show mean ± standard deviation (SD). Paired t-tests comparing means: **e**: *M*_12_ *> M*_1_ *> M*_2_ (all *p <* 0.001); **f**: *M*_12_ ≈ *M*_1_ *< M*_2_ (*M*_12_ vs *M*_1_: ns; *M*_12_ vs *M*_2_ and *M*_1_ vs *M*_2_: *p <* 0.001); **g**: *M*_12_ ≈ *M*_2_ *< M*_1_ (*M*_12_ vs *M*_2_: ns; *M*_12_ vs *M*_1_ and *M*_2_ vs *M*_1_: *p <* 0.001).

The superiority of the model *M*_12_ observed in beta-lactamases holds across most enzyme families, with AIC identifying *M*_12_ as the best model in 27 of 34 cases (Figure 4d; Figures S6d to S39d). Across all enzyme families, *M*_12_ explains more deviance on average (0.49) than *M*_1_ (0.42) or *M*_2_ (0.28) (Figure 4e). Paired t-tests comparing these average explained deviance values confirmed that these differences were statistically significant, with *M*_12_ performing better than *M*_1_ (*p* ≪ 0.001), which in turn performed better than *M*_2_ (*p* ≪ 0.001), demonstrating that *M*_12_ consistently fits better across enzyme families. These results confirm that both non-functional and functional constraints independently contribute to structural divergence.

#### Predicted vs. observed divergence-flexibility and divergence-distance trends

After evaluating the models’ fit to the observed `lRMSD` values, we assessed their ability to reproduce the observed `lRMSD` ∼ `lRMSF` and `lRMSD` ∼ *d* trends. To quantify accuracy, we used two root mean square error (RMSE) metrics: RMSE(flex), which evaluates how well a model captures the divergence-flexibility trend, and RMSE(dist), which assesses its fit to the divergence-distance trend.

In class A beta-lactamases, the single-constraint-type models each capture only one trend while model *M*_12_ captures both. The non-functional-constraints model *M*_1_ accurately reproduces the divergence-flexibility trend (Figure 4b, RMSE(flex) = 0.04) but predicts an overly flat divergence-distance trend (Figure 4c, RMSE(dist) = 0.20). This misfit leads to overestimation of structural divergence near the active site and underestimation at greater distances. Conversely, the functional-constraints model *M*_2_ accurately reproduces the divergence-distance trend (Figure 4c, RMSE(dist) = 0.04) but predicts a too flat divergence-flexibility trend (Figure 4b, RMSE(flex) = 0.18), overestimating divergence in rigid regions and underestimating it in flexible regions. In contrast, *M*_12_, which incorporates both non-functional and functional constraints, accurately reproduces both trends, with RMSE(flex) = 0.04 and RMSE(dist) = 0.04 (Figures 4b and 4c).

The superior performance of model *M*_12_ extends across our entire dataset of enzyme families. As shown in Figures 4f and 4g, and in panels e and f of Figures S6 to S39, *M*_12_ reliably captures both divergence-flexibility and divergence-distance trends across families, unlike the single-constraint-type models *M*_1_ and *M*_2_, which each perform well for one trend but fail for the other (Figures 4f-g). *M*_1_ accurately reproduces the divergence-flexibility trend, with a mean RMSE(flex) of 0.05, but fails to capture the divergence-distance trend, yielding a higher mean RMSE(dist) of 0.18; *M*_2_ accurately reproduces the divergence-distance trend, with a mean RMSE(dist) of 0.07, but fails to capture the divergence-flexibility trend, resulting in a mean RMSE(flex) of 0.29. Model *M*_12_, that considers both non-functional and functional constraints, accurately describes the dependence of structural divergence on both flexibility and distance, offering a more comprehensive explanation than either single-constraint-type model. This is supported by its mean RMSE(flex) of 0.05 and mean RMSE(dist) of 0.08, demonstrating that *M*_12_ captures both trends effectively.

In summary, these results show that model *M*_12_ effectively captures residue-specific structural divergence. Its higher explained deviance and superior fit to both the divergence-flexibility and divergence-distance trends, compared to single-constraint-type models, demonstrates that both non-functional (*s*_1_) and functional (*s*_2_) constraints contribute independently to structural divergence. The combination of these constraints determines a large part of the variation of structural divergence among residues.

### 2.3 Relative contributions of non-functional and functional constraints

Having established that both non-functional and functional constraints contribute independently to structural divergence patterns, we analysed the relative importance of each component. Model *M*_12_ naturally decomposes structural divergence into the sum of non-functional (*s*_1_) and functional (*s*_2_) constraint components. Figure 5a illustrates this decomposition for class A beta-lactamases (M-CSA ID = 2): the residue-dependent profile (blue line) splits into *s*_1_ (green) and *s*_2_ (yellow) components. This decomposition extends to the observed trends (Figures 5b-c): while *s*_1_ primarily determines the divergence-flexibility relationship and *s*_2_ primarily determines the divergence-distance relationship, each component also influences both trends indirectly because flexibility and distance are themselves correlated. While beta-lactamases represent a typical case, similar decompositions across all 34 enzyme families (shown in Figures S6-S39, panels g-i) reveal substantial variation in the relative contributions of non-functional and functional constraints.

**Figure 5:**
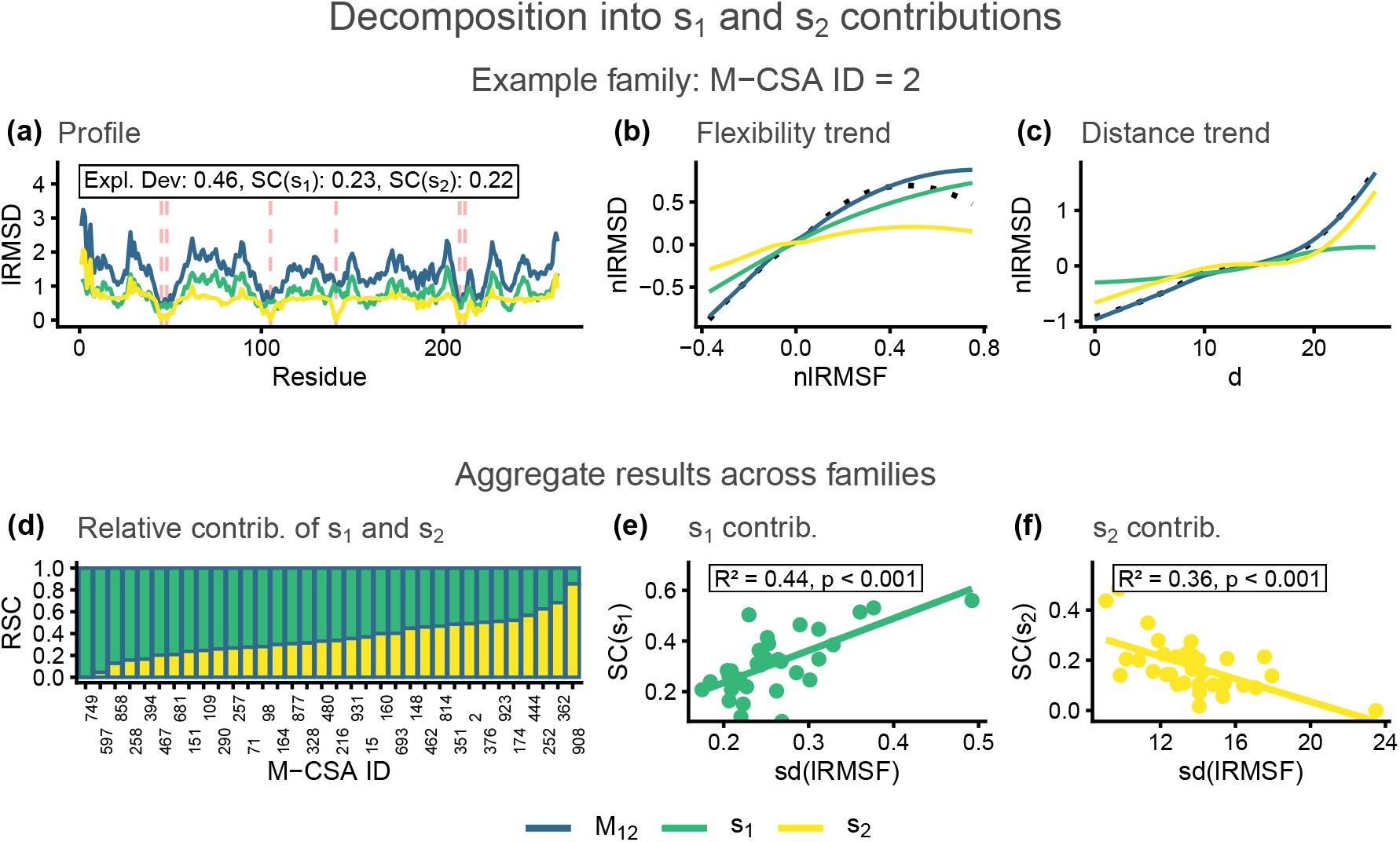
Decomposition of structural divergence into *s*_1_ and *s*_2_ contributions. **a**: Residue-dependent `lRMSD` profile for class A beta-lactamases (M-CSA ID = 2), showing *M*_12_ predictions (blue, constant term removed) decomposed into functions *s*_1_ (green) and *s*_2_ (yellow). Active-site residues are indicated by vertical dashed red lines. The model’s explained deviance and Shapley contributions are shown. **b**: Flexibility trend (`nlRMSD` vs. `nlRMSF`) for the same family. Black dotted line shows LOESS fit to observations, coloured lines show LOESS fits to *M*_12_ predictions (blue) and its components *s*_1_ (green) and *s*_2_ (yellow). **c**: Distance trend (`nlRMSD` vs. *d*) for the same family. Lines as in **b. d**: Stacked bars showing the relative Shapley contributions of *s*_1_ and *s*_2_ across enzyme families, sorted by increasing *s*_2_ contribution. **e**: Relationship between *s*_1_ contribution (*SC*(*s*_1_)) and the standard deviation of residue flexibilities within each enzyme. **f**: Relationship between *s*_2_ contribution (*SC*(*s*_2_)) and the mean residue distance to active site within each enzyme. Linear fits with corresponding *R*^2^ and p-values are shown in **e** and **f**.

To quantify these contributions, we applied the Shapley method [46]. Inspired by game theory, this approach provides a fair allocation of each variable’s contribution while accounting for their interactions. It distributes the explained deviance among *s*_1_ and *s*_2_ by averaging their marginal contributions across all models, ensuring fair attribution even when the components are correlated (see Methods).

The relative importance of non-functional and functional constraints varies widely among enzyme families. For class A beta-lactamases, the balance is nearly equal, with a total explained deviance of 0.46 and Shapley values of 0.23 and 0.22 for *s*_1_ and *s*_2_, respectively, representing nearly equal relative contributions of 51% and 49% of the explained deviance. However, this equal distribution is not uniform across the dataset. The explained deviance of model *M*_12_ averages 0.49 across families, with *s*_1_ contributing 0.31 and *s*_2_ 0.18. Thus, on average, *s*_1_ accounts for 64% of the explained deviance and *s*_2_ for 36%. However, these averages mask the wide variation across enzyme families. Excluding M-CSA ID 749, which shows no functional contribution and is discussed as a special case in Section 2.5, the contribution of functional constraints increases from 4% to 85% across families (with the contribution of non-functional constraints accordingly decreasing from 96% to 15%).

To understand why the contributions of *s*_1_ and *s*_2_ to structural divergence vary among enzymes, we identified properties that could predict these contributions. Through stepwise regression (see Methods), we found that only two properties account for most of the variation: `sd`(`lRMSF`), which measures how much flexibility varies among residues within a protein, and ⟨*d*⟩, the mean distance to the active site. The contribution of *s*_1_ increases with `sd`(`lRMSF`) (*R*^2^ = 0.44, p < 0.001, Figure 5e), while the contribution of *s*_2_ decreases with ⟨*d*⟩ (*R*^2^ = 0.36, p < 0.001, Figure 5f). A multiple regression model combining both predictors (lm(rsc(*s*_2_) sd(lrmsf) + ⟨*d*⟩)) explains 38% of the variation in relative contributions between families.

These predictors reflect aspects of protein architecture. `sd`(`lRMSF`) represents the heterogeneity in residue flexibility that results from the topology of residue-residue contact networks, as captured by elastic network models. ⟨*d*⟩ reflects the spatial distribution of residues relative to the active site, which is determined by protein size (larger proteins having more distant residues), shape (elongated versus spherical), and active site positioning (peripheral versus central). While these architectural features explain 38% of the variation in relative contributions of *s*_1_ and *s*_2_ across enzyme families, the remaining unexplained variation implies that other factors, possibly including varying degrees of selection pressure, also significantly influence which constraints dominate structural divergence in different enzyme families.

### 2.4 Robustness to reference protein and atom choice

To assess the robustness of our results, we tested whether the model performance metrics (Section 2.2) and constraint contributions (Section 2.3) are sensitive to how we calculate the predictors `lRMSF` and *d*, and the observed `lRMSD` profiles. Specifically, we examined robustness with respect to reference protein choice and atom choice.

We compared results using the designated reference protein versus the structurally closest homologue for calculating the predictors (`lRMSF` and *d*). (The the closest homologue is that with lowest overall RMSD to the reference, selected to reduce artifacts from gaps and non-evolutionary structural differences.) Protein choice has minimal impact on our findings: goodness-of-fit metrics exhibit strong correlations between the two approaches (*r* = 0.77–0.94; Figure S3), with nearly identical average values across all metrics (Table 1).

**Table 1:**
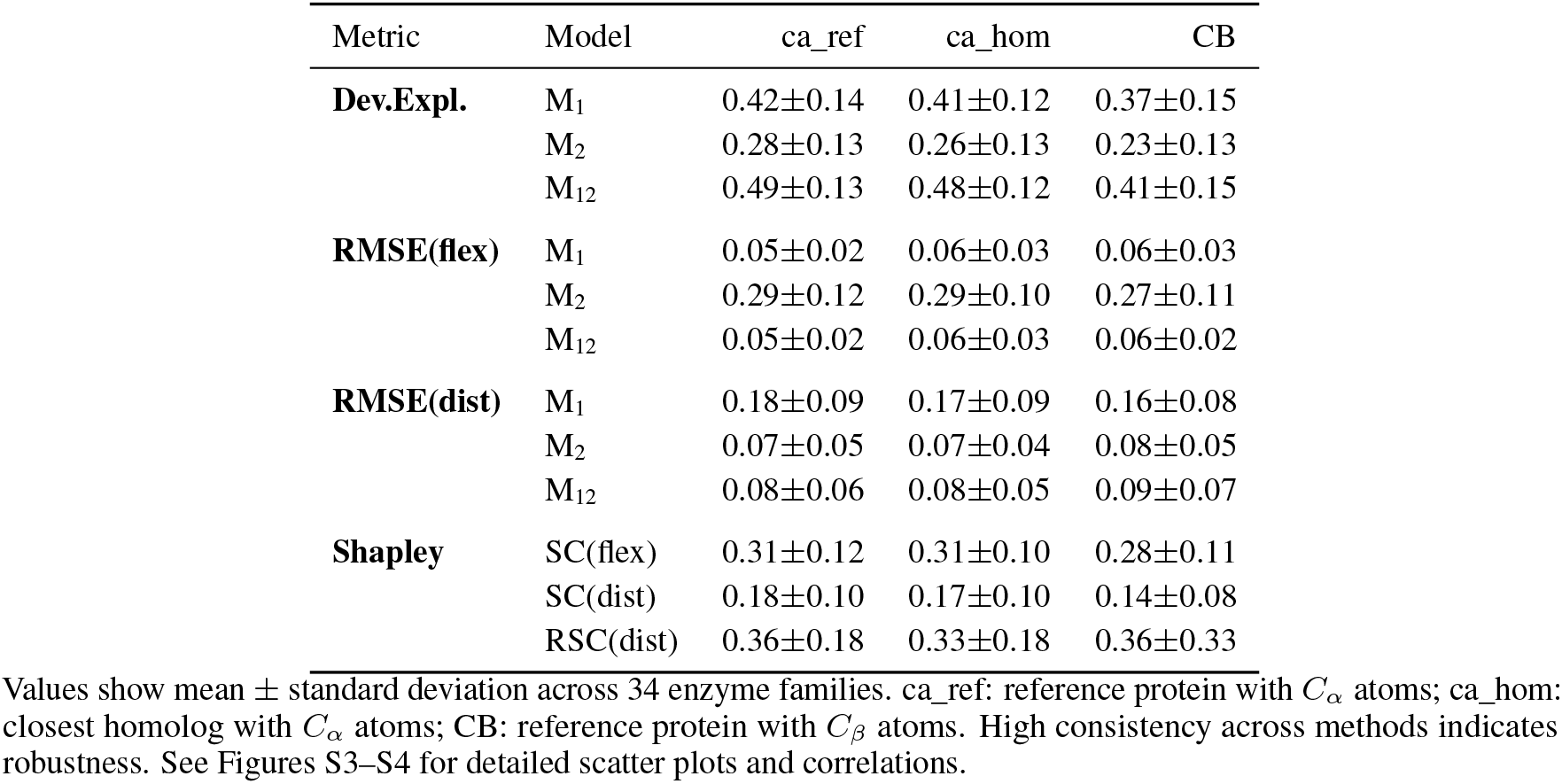
Robustness of goodness-of-fit metrics to methodological choices.

| Metric | Model | ca_ref | ca_hom | CB |
| --- | --- | --- | --- | --- |
| **Dev.Expl.** | M<sub>1</sub> | 0.42±0.14 | 0.41±0.12 | 0.37±0.15 |
|  | M<sub>2</sub> | 0.28±0.13 | 0.26±0.13 | 0.23±0.13 |
|  | M<sub>12</sub> | 0.49±0.13 | 0.48±0.12 | 0.41±0.15 |
| **RMSE(flex)** | M<sub>1</sub> | 0.05±0.02 | 0.06±0.03 | 0.06±0.03 |
|  | M<sub>2</sub> | 0.29±0.12 | 0.29±0.10 | 0.27±0.11 |
|  | M<sub>12</sub> | 0.05±0.02 | 0.06±0.03 | 0.06±0.02 |
| **RMSE(dist)** | M<sub>1</sub> | 0.18±0.09 | 0.17±0.09 | 0.16±0.08 |
|  | M<sub>2</sub> | 0.07±0.05 | 0.07±0.04 | 0.08±0.05 |
|  | M<sub>12</sub> | 0.08±0.06 | 0.08±0.05 | 0.09±0.07 |
| **Shapley** | SC(flex) | 0.31±0.12 | 0.31±0.10 | 0.28±0.11 |
|  | SC(dist) | 0.18±0.10 | 0.17±0.10 | 0.14±0.08 |
|  | RSC(dist) | 0.36±0.18 | 0.33±0.18 | 0.36±0.33 |

We also compared results using *C*_*α*_ atoms versus *C*_*β*_ atoms for calculating all structural measures (`lRMSD`, `lRMSF`, and *d*). Atom choice shows more pronounced effects: while *C*_*α*_- and *C*_*β*_-based analyses show significant correlations (*r* = 0.60–0.84; Figure S4), these correlations are weaker than for reference protein choice, and *C*_*β*_-based analyses consistently yield lower goodness-of-fit values across all metrics (Table 1). The weaker performance of *C*_*β*_-based analyses likely reflects the greater conformational variability of side chains compared to the backbone, which introduces additional noise into structural divergence measurements that is not purely evolutionary in origin.

These results confirm that our findings are robust to reference protein selection and validate our use of *C*_*α*_ atoms and the designated M-CSA reference proteins for all structural calculations.

### 2.5 Model performance across some example cases

To complement our general analysis of structural divergence patterns, we examine five specific cases that span the range of observed behaviours. Using the distribution of enzyme families based on their contributions of non-functional and functional constraints (Figure 6), we selected cases representing the full range of relative contributions of functional constraints to structural divergence: the aldo/keto reductase family (M-CSA ID 858; Figure S35) shows minimal contribution (RSC(*s*_2_) = 0.13); a metallo-beta-lactamase (M-CSA ID 15, RSC(*s*_2_) = 0.37; Figure S7), a class-A beta-lactamase (M-CSA ID 2, RSC(*s*_2_) = 0.49; Figure S6), and a TrpC family member (M-CSA ID 252, RSC(*s*_2_) = 0.63; Figure S17) show intermediate values; and the ribonuclease U2 family (M-CSA ID 908; Figure S37) shows the strongest functional constraints contribution (RSC(*s*_2_) = 0.85).

**Figure 6:**
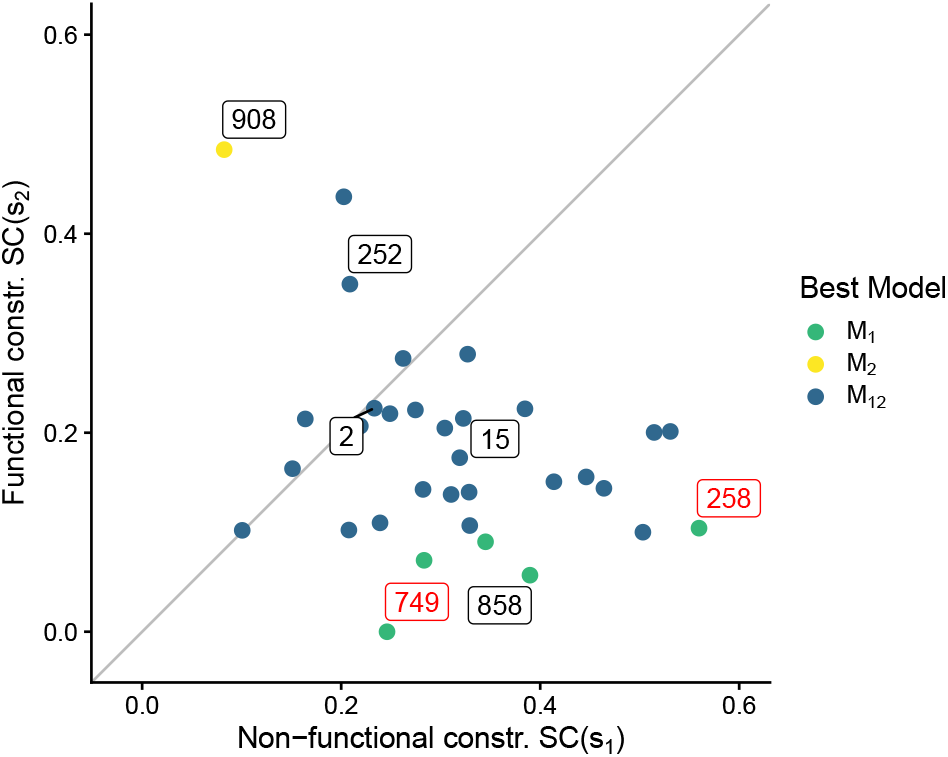
Map of enzyme families based on their Shapley contributions. Each point represents an enzyme family positioned according to its Shapley contributions from non-functional constraints (*SC*(*s*_1_), x-axis) and functional constraints (*SC*(*s*_2_), y-axis). The diagonal line indicates equal contributions from both. Colours indicate the best model (*M*_1_, *M*_2_, or *M*_12_ according to AIC, with *M*_1_ dominating at high non-functional contributions, *M*_2_ at high functional contributions, and *M*_12_ at intermediate values). Labels identify five representative cases (M-CSA IDs 858, 15, 2, 252, and 908, in black boxes, ordered by increasing relative contribution of functional constraints) and two outliers (IDs 258 and 749, in red boxes) that are discussed in Section 2.5.

Table 2 shows several trends across the five cases. As the relative contribution of functional constraints increases, protein size decreases (from 322 to 107 residues) and mean distance to active site shortens (from 15.3Å to 9.8Å). These changes align with the shift from *s*_1_ to *s*_2_ contributions. The spread of local flexibility, `sd`(`lRMSF`), which we showed drives *s*_1_ contributions, shows no consistent trend in these cases. This pattern illustrates our earlier finding that the spread of flexibility and mean distance from the active site explain only 38% of variation in relative contributions between constraints, with the remaining variation potentially arising from other factors such as the variation among families of selection pressure.

**Table 2:**
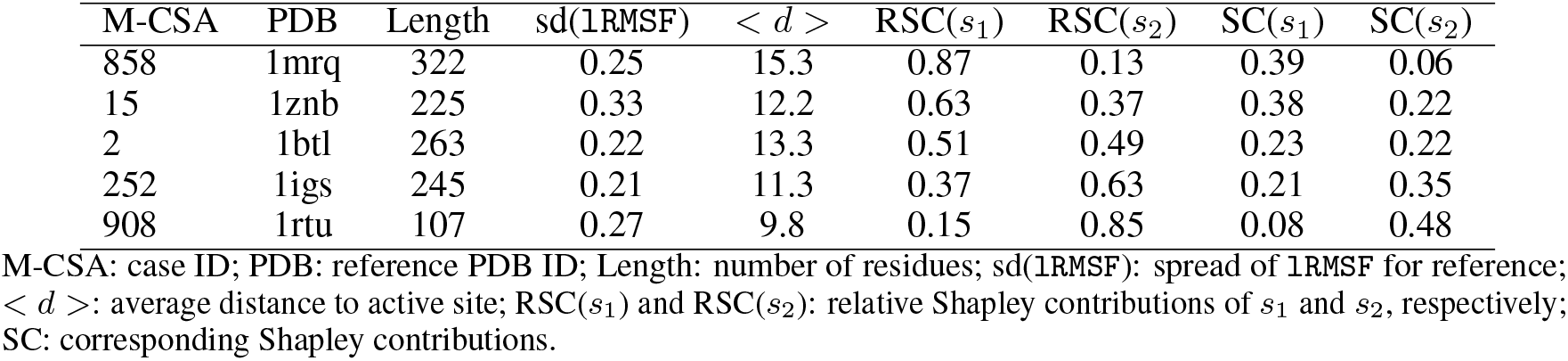
Contributions of non-functional and functional constraints in example families.

The explained deviance of each model reflects which type of constraint dominates (Table 3, columns Expl. Dev.). When the *s*_2_ contribution is low (M-CSA ID 858), *M*_1_ performs well and *M*_2_ poorly, with *M*_12_ matching *M*_1_’s performance. Through the intermediate cases (M-CSA 15, 2, and 252), neither single-constraint-type model alone suffices, and *M*_12_ outperforms both. When the *s*_2_ contribution is high (M-CSA ID 908), *M*_2_ performs well and *M*_1_ poorly, with *M*_12_ matching *M*_2_’s performance. (Although *M*_12_ explains slightly more deviance than *M*_2_ in this case, *M*_2_ is selected as the best model by AIC due to its lower complexity.)

**Table 3:**
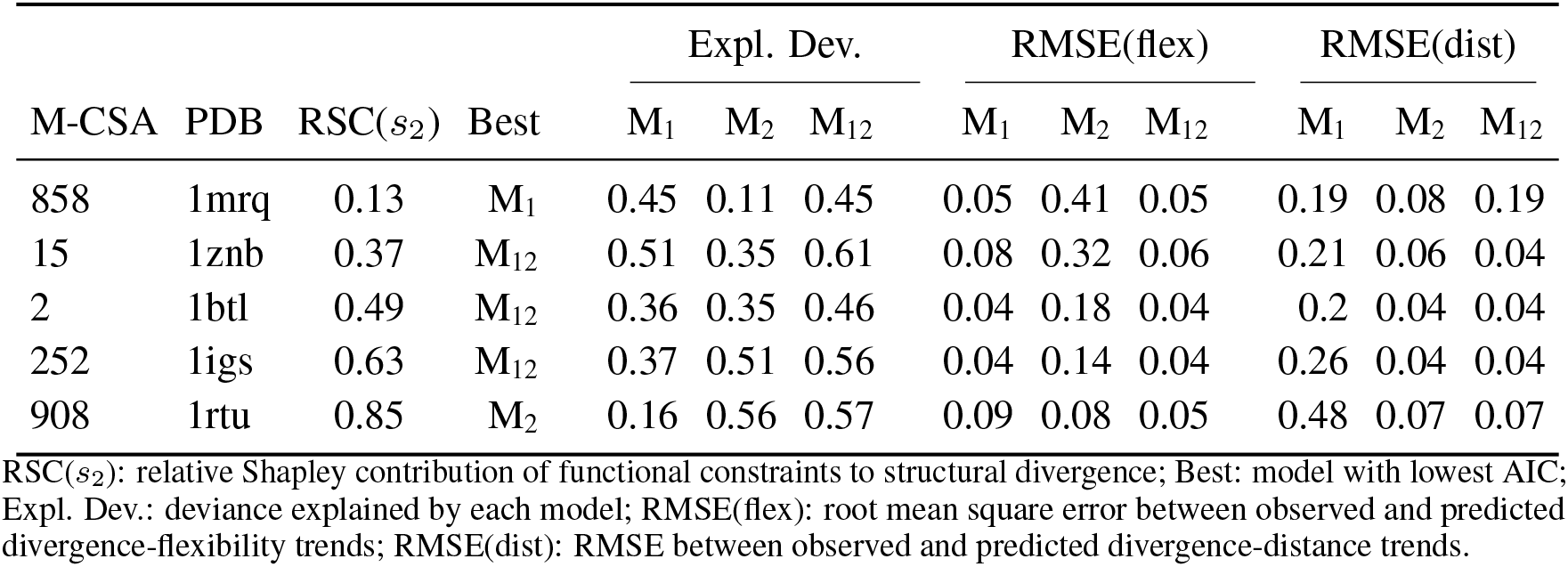
Variation of model performance across example families.

Each model’s ability to reproduce the divergence-flexibility and divergence-distance trends reveals their limitations (Table 2, columns RMSE(flex) and RMSE(dist)). *M*_1_ accurately captures the flexibility trend but, except when the functional constraint contribution is minimal, fails to account for the distance trend. Conversely, *M*_2_ captures the distance trend but, except when the functional constraint contribution is maximal, fails to account for the flexibility trend. Only *M*_12_ reproduces both trends across all cases, demonstrating that both constraints are needed to explain the observed patterns of variation.

The cases considered here illustrate how the balance of constraints determines model performance. Single-constraint-type models succeed only when their constraint type dominates, while *M*_12_ excels across all cases by incorporating both constraints.

### 2.6 Active site structural divergence

Having established that both non-functional and functional constraints influence structural divergence across residues, we examined whether this dual influence also determines the structural conservation of enzyme active sites. Active sites are traditionally assumed to be structurally conserved because of functional constraints. However, their typical location in rigid regions, where non-functional constraints are high, suggests that non-functional constraints may also contribute to their conservation.

We first confirmed that active-site residues are more structurally conserved than average across enzyme families (Figure 7a; Table S4). Model *M*_12_ accurately reproduces this conservation (observed mean = − 0.62 ±0.28, predicted mean = −0.69 ± 0.22, *R* = 0.86; Table S4), indicating that it captures the determinants of active-site structural conservation.

**Figure 7:**
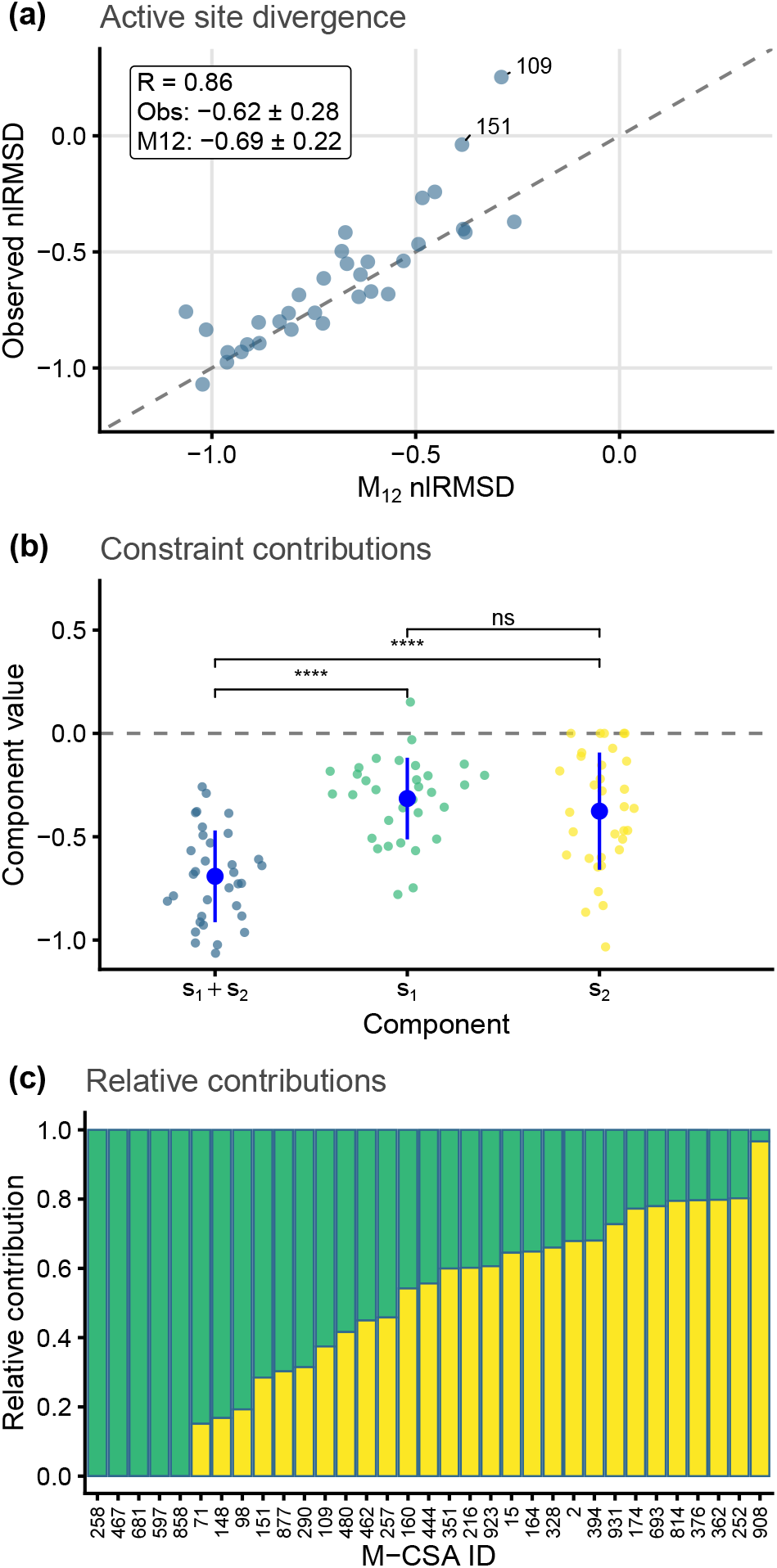
Structural divergence of enzyme active sites. **(a)** Observed versus predicted structural divergence (`nlRMSD`) of active sites across enzyme families; each point corresponds to the mean of residue values within an active site. **(b)** Components of model *M*_12_ at active sites, averaged over active-site residues: total predicted divergence (*s*_1_ + *s*_2_, blue), non-functional component (*s*_1_, green), and functional component (*s*_2_, yellow). Large points show mean ± SD. Paired t-test significance: **** *p <* 0.0001; ns *p >* 0.01. **(c)** Relative contributions of non-functional (*r*_*s*1_ = |*s*_1_| */*(|*s*_1_| + |*s*_2_|), green) and functional (*r*_*s*2_ = |*s*_2_| */*(|*s*_1_| + |*s*_2_|), yellow) constraints to active-site conservation across enzyme families (M-CSA IDs). Ratios were computed per residue and averaged within each active site.

To assess the contributions of each constraint type at active sites, we used the *s*_1_ and *s*_2_ components of *M*_12_ obtained in Section 2.3. For each enzyme family, we averaged *s*_1_ and *s*_2_ over active-site residues to quantify the strength of non-functional and functional constraints at those positions (Figure 7b). Both components contribute significantly to active-site conservation (mean *s*_1_ = −0.32 ±0.20, mean *s*_2_ = −0.38± 0.28, both *p <* 0.0001; Table S4), showing that non-functional constraints contribute nearly as much as functional ones on average.

To compare their relative magnitudes, we defined the normalized contributions

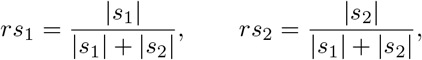

where higher absolute *s* values indicate stronger constraints (departure of *s* from the mean *s* = 0). These metrics quantify the relative magnitude of each constraint type at the active site, irrespective of their sign. Across enzyme families, *rs*_1_ ranges from 0.03 to 1.00, with *rs*_2_ varying inversely from 0.97 to 0.00 (Figure 7c; Table S5), indicating that some active sites are dominated by non-functional constraints while others by functional ones.

The balance between constraint types correlates with active-site rigidity: more rigid active sites show higher *rs*_1_ values (Figure S5a,c), consistent with the expectation that rigidity strengthens non-functional constraints.

Active-site structural conservation therefore results from both constraint types. Non-functional constraints, arising from the location of active sites in rigid regions, contribute substantially, with their relative importance increasing with active-site rigidity. On average, functional and non-functional constraints contribute approximately equally to maintaining active-site structure.

## 3 Discussion

### 3.1 Patterns of structural divergence and their constraints

Our analysis revealed two trends in how structural divergence varies among residues: divergence increases with both flexibility and distance from active sites (Section 2.1). While these trends suggest non-functional and functional constraints, their flexibility-distance correlation complicates isolating each constraint’s contribution.

Previous models attribute the divergence-flexibility trend to mutational effects and stability requirements [34], and the divergence-distance trend to functional constraints weakening with distance [38, 39]. However, because flexibility and distance are correlated, the trends alone cannot distinguish each constraint’s independent contribution.

Model *M*_12_ (Section 2.2) resolves this by separating divergence into independent non-functional (*s*_1_(`lRMSF`)) and functional (*s*_2_(*d*)) constraint components, revealing their combined direct and indirect effects (Section 2.3). Its success in reproducing both trends, outperforming single-constraint-type models, shows both constraints independently shape structural evolution, with their relative strength determining flexibility and distance dependence.

### 3.2 Variation in constraint balance across enzymes

The relative contributions of non-functional and functional constraints vary substantially across enzyme families. The contribution of functional constraints ranges from 4% to 85% of explained structural divergence, with non-functional constraints correspondingly decreasing from 96% to 15% (Figure 5d). These shifts in constraint balance reshape how structural divergence depends on flexibility and distance.

What determines which type of constraint dominates in a particular enzyme family? Our analysis shows that protein architecture influences constraint balance through two properties: the spread of flexibility among residues and the mean distance to active sites (Section 2.3). When flexibility varies widely within a protein structure, non-functional constraints become stronger determinants of structural divergence patterns (Figure 5e). In larger or more elongated proteins where residues tend to lie far from active sites, functional constraints become less influential (Figure 5f). Together, these architectural properties explain 38% of the differences in constraint balance among enzyme families.

The modest explanatory power of architecture suggests other factors influence the balance between non-functional and functional constraints. This is further supported by our observation that enzymes with similar architectural properties can show different patterns of structural divergence (Section 2.5). Previous sequence-based studies suggest that differences in functional selection pressure among enzymes could explain part of this remaining variation [39]. Testing this hypothesis will require mechanistic biophysical models that can explicitly account for both protein architecture and varying selection regimes.

### 3.3 Relationship between evolutionary and dynamical ensembles

Previous work has stressed the similarity between sets of structures related by homology with sets of conformations reached by the same protein during its dynamics [25, 26, 27, 28, 29, 30, 31, 32, 33, 34, 35] (see Introduction). If only non-functional constraints operated, then we would expect evolutionary and dynamical ensembles to be similar, with model *M*_12_’s non-functional term (*s*_1_) aligning divergence to dynamics [27, 30, 34]. However, the present results show that structural divergence patterns depend also on functional constraints, captured by *M*_12_’s functional term (*s*_2_) (Section 2.2). As functional constraints (*s*_2_) increase, their contribution to structural divergence becomes stronger, making evolutionary patterns increasingly different from those expected from non-functional constraints (*s*_1_) alone. The similarity between evolutionary and dynamical ensembles thus exists only to the extent that non-functional constraints dominate over functional ones. This finding revises our understanding of protein evolution by showing that the correspondence between evolutionary variation and protein dynamics is not universal but depends on the balance between these constraints.

### 3.4 Structural conservation of active sites

Active sites are more structurally conserved between homologous enzymes than between random proteins [36], and more conserved than the protein average [37]. This conservation is usually taken as evidence of functional constraints, but our results in Section 2.6 show that this interpretation is incomplete. Because active sites typically lie in rigid regions, where non-functional constraints are strong, part of their conservation reflects background structural effects rather than function alone. Decomposing model *M*_12_ shows that both constraint types contribute comparably, on average, to active-site conservation, with their relative weight varying widely among enzyme families. Non-functional contributions increase with active-site rigidity, whereas functional contributions dominate in more flexible active sites. Thus, the structural conservation of active sites stems from both constraint types, and cannot be interpreted as evidence of functional constraints alone.

### 3.5 Exceptions and special cases

Two enzyme families deviate from the general relationship between structural divergence and distance to active sites described in Section 2.1. M-CSA ID 258, an L1 metallo-beta-lactamase from Stenotrophomonas maltophilia, shows no correlation between divergence and distance to the active site. As a hospital-derived pathogen conferring broad-spectrum antibiotic resistance, it may have undergone recent adaptive evolution under selective pressures that differ from those leading to active site conservation in other enzymes.

M-CSA ID 749, a cytochrome c peroxidase, shows a negative correlation between divergence and distance, violating our assumption that functional constraints weaken with distance from active sites. Examination of the structures in the PDB reveals this apparent divergence actually reflects conformational changes between calcium-bound and calcium-free states rather than evolutionary differences. This explains why this family shows no contribution from function *s*_2_ (Section 2.3): what we measure is not evolutionary divergence but conformational variation.

While we could have detected and filtered out these cases by examining active site structural differences, doing so would have introduced circularity in our analysis of active site conservation. Instead, their detection as statistical outliers serves to validate our analytical approach by showing it can identify cases that violate its assumptions. Notably, the case of the L1 metallo-beta-lactamase highlights how adaptive evolution can produce patterns of structural divergence that differ from those we observe under purifying selection.

### 3.6 Model performance and future directions

Our model explains on average 49% of the variation in structural divergence using two interpretable predictors: flexibility and distance from the active site. This level of agreement between observed and predicted patterns exceeds that typically achieved in studies of sequence evolution rate variation among residues, where explained variance ranges from 32-44% [38, 24]. For sequence evolution, decades of research building on similarly constrained initial models have progressively refined our understanding through iterative improvement of predictors and expansion of scope.

The remaining unexplained variance defines clear targets for future work. Some may arise from factors we have not yet considered: cofactor binding sites may create additional functional constraint centres; allosteric sites could function as secondary regions of strong selection; protein-protein interaction interfaces in multimeric enzymes may introduce constraints absent in monomers. Others may come from refining the factors we do consider: alternative flexibility estimates (from all-atom molecular dynamics or NMR ensembles rather than elastic network models), improved distance metrics that account for the specific functional roles of different active-site residues (catalytic versus substrate-binding versus scaffolding), or alternative measures of functional importance beyond simple geometric distance. Each potential improvement is testable precisely because the current model provides quantitative predictions to compare against.

Our dataset was deliberately restricted to monomeric single-domain enzymes to avoid confounding effects from domain-domain and chain-chain interactions. This raises natural questions about generalizability. Can the two-constraint framework extend to more complex architectures? For multi-domain proteins with inter-domain active sites, which domain’s flexibility profile should dominate? Do allosteric coupling mechanisms in multimeric enzymes require additional terms in the model? While we cannot assume the current model generalizes without testing, the framework is applicable: we can calculate flexibility and distance measures for any protein structure, fit the model, and determine whether the two-constraint decomposition captures the patterns or whether additional factors are needed. Systematic comparison between monomeric and multimeric enzymes, or between single-domain and multi-domain proteins, would reveal whether our findings represent general principles or require architecture-specific extensions.

The model’s interpretable simplicity—two smooth functions explaining approximately half the variance (49% on average)—makes it a useful baseline for future research. The questions we can now pose about cofactor sites, allosteric coupling, interaction interfaces, or refined distance metrics arise precisely because we have established a clear quantitative target. The following sections explore some particularly promising directions this framework enables.

### 3.7 Contrasts with Adaptive Evolution

The L1 metallo-beta-lactamase example illustrates a broader point: under purifying selection, both functional and non-functional constraints contribute to the structural conservation of active sites, with non-functional constraints playing a substantial role. In contrast, during adaptive evolution, where new enzymatic functions are selected for, active sites undergo structural divergence to accommodate novel functional demands [47]. Under these adaptive scenarios, positive selection drives mutations that remodel the active site, with functional constraints primarily shaping these changes to favour new activities. The potential influence of non-functional constraints—such as the inherent mutational robustness of active site regions and the rejection of destabilizing mutations—remains insufficiently studied in adaptive scenarios. While a complete treatment of structural evolution under positive selection is beyond the scope of this work, we speculate that non-functional constraints may limit the spectrum of structurally viable adaptations, even under positive selection, and suggest that future studies evaluate their impact on shaping the evolutionary pathways of adaptive enzymes.

### 3.8 Applications to structural phylogenetics

Our findings have immediate applications for structural phylogenetics, a field of growing importance given the increased availability of protein structures through methods like AlphaFold. Understanding how structural divergence varies among residues suggests several possible improvements to existing approaches. For example, distance-based methods [13, 14, 15] could incorporate residue-specific weights based on flexibility and distance to active sites, while probabilistic models [8, 10, 11, 12, 16, 17] could account for varying structural divergence rates that depend on these variables.

### 3.9 Physical basis of protein structural evolution

Our empirical analysis revealed two types of constraints that independently shape structural divergence. These patterns suggest fundamental mechanisms in protein evolution. Non-functional constraints point to effects arising from basic protein physics: how different residues respond to mutations and how stability requirements influence where mutations are tolerated. These mechanisms would operate similarly across all proteins. Functional constraints, in contrast, point to effects arising from specific molecular interactions required for protein function, which would vary among proteins according to their particular roles.

Some biophysical models address aspects of these potential mechanisms: how mutational biases and selection on stability shape structural divergence [27, 30, 34], how functional constraints operate through ligand binding [48], while extensions of stability-based models that incorporate selection on catalytic activity have been applied to sequence evolution patterns [49, 39]. However, a complete mechanistic understanding of our findings will require models that integrate all these elements: how mutations affect protein structure and how selection on both stability and function governs their fixation. Such models would provide a mechanistic basis for the constraints we formalized through functions *s*_1_ and *s*_2_.

## 4 Materials and Methods

### 4.1 Dataset

To study the constraints driving enzyme structural divergence, we curated a dataset from the M-CSA database, focusing on entries where the reference protein is a monomeric single-domain enzyme. These represent the simplest case for analysing evolutionary constraints, free from complexities introduced by inter-domain or inter-chain interactions. Since we are focusing on a negative selection scenario in which function is conserved, homologues were selected to ensure functional conservation with their reference enzyme, retaining only proteins with conserved catalytic and substrate-binding residues and sufficient sequence similarity, as detailed below.

#### Initial filtering

The dataset was derived from the M-CSA (Mechanism and Catalytic Site Atlas) database, accessed on August 21, 2023. M-CSA is a manually curated database of catalytic residues and enzyme mechanisms, accessible at www.ebi.ac.uk/thornton-srv/m-csa/[42, 43]. It contains 1002 entries and extends to over 131,807 structures through homologue searches in the Protein Data Bank. Homologues are identified using PHMMER with an E-value threshold of 10^−6^ [37]. Each M-CSA entry includes a reference enzyme structure with manually annotated catalytic residues.

We focused on the simplest case of monomeric single-domain enzymes, as multi-domain and multi-chain proteins introduce additional complexity through domain and chain interactions that require separate analysis.

Starting from 1002 entries, we applied sequential filtering steps (remaining entries after each step shown in parentheses):

- Removal of inconsistencies and duplicates (921)
- Filtering for entries with intra-chain active sites (795)
- Filtering for X-ray determined structures (786)
- Filtering for entries with intra-domain active sites (568) (Note that this step became redundant after filtering for single-domain proteins, as they cannot have inter-domain active sites.)
- Filtering for single domain reference proteins (313)
- Filtering for monomeric biological assemblies (120)
- Filtering for entries with homologues with the same CATH ID as the reference (106)
- Within each remaining entry: for homologues only, when multiple PDB structures exist for the same UniProt ID, selecting the optimal structure based on sequence coverage and X-ray resolution

This filtering process resulted in a dataset of 106 enzyme families. Each entry comprises a manually annotated reference PDB chain and its homologues, totalling 1171 homologue structures. All reference enzymes are monomeric and single-domain.

#### Alignment, Superposition, and Further Filtering

For each entry, the reference and homologous chains were aligned using Muscle V (via `pdb.aln` in the `bio3d` R package) [50, 51]. Sequence alignment was chosen over structural alignment because (1) within families, sequences are similar enough for reliable alignments, and (2) it provides an independent basis for studying structural divergence.

Following alignment, structures were superimposed to minimize the root mean square deviation (`RMSD`). The dataset then underwent additional filtering:

- Removal of homologues with gaps or mutations at active-site residues (including catalytic and substrate-binding amino acids, ensuring the same reaction and substrate specificity)
- Clustering-based removal of structural outliers and redundant structures. We removed clusters with RMSD > 6Å from the reference structure (to exclude likely alternative conformational states) and merged clusters with RMSD < 1Å (to avoid redundancy from near-identical structures)
- Clustering-based filtering of sequence identity, removing clusters with identity > 95% (to reduce redundancy) or < 25% (below the SCOP threshold for functional conservation within families) relative to the reference structure
- Removal of families with fewer than 4 homologues

This process yields a dataset of non-redundant enzymes for each M-CSA entry, likely sharing the same function as their reference enzyme, as evidenced by sequence and structural similarity and identical active-site amino acids.

#### Dataset Properties

The final dataset contains 34 enzyme families: 27 alpha/beta, 5 all-beta, and 2 all-alpha structures. CATH architectures include Alpha-Beta Barrel, 3-Layer(aba) Sandwich, Alpha-Beta Complex, 4-Layer Sandwich, Distorted Sandwich, 2-Layer Sandwich, and Orthogonal Bundle. The EC distribution shows hydrolases (20 families) as most frequent, followed by transferases, lyases, isomerases, and oxidoreductases. Each family contains 4-23 proteins (mean 6.82, SD 3.98). Reference proteins range from 98 to 447 residues (mean 248.50, SD 91.31), and Id% ranges from 0.31 to 0.56 (mean 0.41, SD 0.06). `RMSD` ranges from 0.72Å to 2.88Å (mean 1.38, SD 0.45). Active sites comprise 1-8 residues (mean 4.74, SD 1.76), with active site `RMSD` from 1.22Å to 6.68Å (mean 2.26, SD 0.96). The highest active site `RMSD` occurs in M-CSA ID 749, where the reference structure lacks a calcium ion present in homologues. See Figure S1 for distributions.

### 4.2 Residue-specific structural divergence: RMSD

Let **R**_1_, **R**_2_, …, **R**_*N*_ represent the structures of *N* homologous enzymes. Each **R**_*p*_ is a vector of Cartesian coordinates of the alpha carbons (*C*_*α*_) in the native structure of enzyme *p*. These structures form a sample from the set of structures that evolved from the proteins’ common ancestor. We quantify the structural divergence of a residue *i* using its root mean square deviation, `RMSD`:

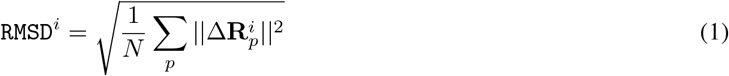

Here, 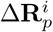 is the deviation of residue *i*’s position vector in protein *p* from its average position across all proteins.

The direct use of `RMSD` presents two challenges. First, its wide variation range can obscure smaller variations in conserved regions. Second, the error in `RMSD` estimates is proportional to `RMSD` itself. To address these issues, we use the natural logarithm of `RMSD`, `lRMSD` = ln(`RMSD`). This transformation converts multiplicative error into additive error (independent of measured value) and gives appropriate weight to small `RMSD` values in conserved regions.

We then define a normalized measure:

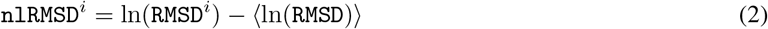

where ⟨ln(`RMSD`)⟩ is the average logarithmic RMSD across all residues in the protein. This normalized logarithmic root mean square deviation, `nlRMSD`, provides a relative measure of structural divergence. Negative values indicate residues that are structurally less diverged than average, while positive values indicate more diverged than average residues.

Implementation uses MUSCLE v5 [50] for sequence alignment, followed by structural superposition using `bio3d` (version 2.4.4) [51]. Each structure is superimposed to a reference by minimizing overall `RMSD`. We then calculate residue-specific values: `RMSD` (Eq.1, excluding gaps), its natural logarithm `lRMSD`, and the normalized version `nlRMSD` (Eq.2). We use `lRMSD` when analysing absolute divergence and `nlRMSD` when comparing divergence patterns across proteins of different sizes or overall divergence levels.

### 4.3 Residue-specific Flexibility: RMSF

We correlate evolutionary divergence with dynamical fluctuations, building on prior research [34]. We designate one homologous enzyme as the reference. This enzyme fluctuates, producing an equilibrium ensemble of conformations described by the distribution function *ρ*_dyn_(**r**), where **r** represents any given conformation. The native structure is the average conformation: **R** = ⟨**r**⟩.

We quantify the flexibility of residue *i* using the root mean square fluctuation (`RMSF`):

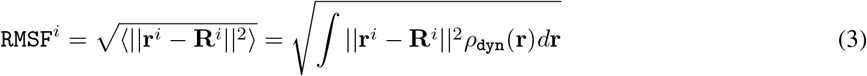

Here, **r**^*i*^ is the position vector of residue *i* in conformation **r**, and **R**^*i*^ is its position in the native conformation. The equilibrium distribution function, *ρ*_dyn_(**r**), follows from basic statistical mechanics:

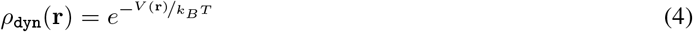

where *V* (**r**) is the potential energy, *k*_*B*_ the Boltzmann constant, and *T* the absolute temperature.

We obtain the potential energy using the elastic network model (ENM) developed by Ming and Wall [52]. This coarse-grained model represents the protein as a network of nodes (residues) connected by springs (interactions). The potential energy landscape of the elastic network is:

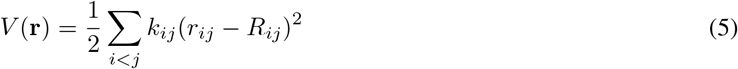

where *r*_*ij*_ is the distance between residues *i* and *j* in conformation **r**, and *R*_*ij*_ is the distance in the native conformation. The spring constant *k*_*ij*_ is defined as:

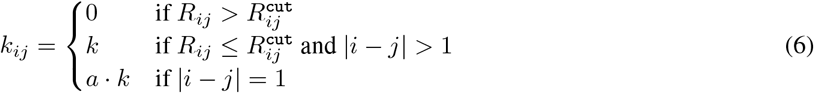

where 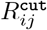 is a cut-off distance, and *a* is a scaling factor for consecutive residues. The parameter values of this model are 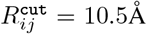, *k* = 4.5, and *a* = 42. This model was shown to yield accurate `RMSF` predictions [52], and it has been used to study non-functional constraints [34]. In the later study, this model was compared with 3 alternative ENM models (Hinsen’s ENM [53], the Anisotropic Network Model [54], and the parameter-free Anisotropic Network Model [55]), and found to lead to very similar results. ENMs work because of this robustness, and, therefore, results are largely independent of ENM model choice [34].

We compute `RMSF` by replacing Eq.5 into Eq.4, and applying the result in Eq.3. As with structural divergence, we use both the natural logarithm of `RMSF` (`lRMSF`) and a normalized version:

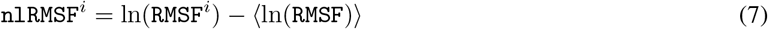

Like `nlRMSD`, `nlRMSF` provides a relative measure, with negative and positive values indicating residues less or more flexible than average, respectively. We use `lRMSF` when analysing absolute flexibility and `nlRMSF` when comparing flexibility patterns across proteins.

### 4.4 Distance from the active site: *d*

We define *d* for each residue as the minimum distance between its *C*_*α*_ and those of active-site residues.

### 4.5 Divergence-flexibility and divergence-distance trends

For each enzyme family, we analyse how `lRMSD` varies with `lRMSF` and *d*. We quantify these relationships through smooth fits (which we refer to as “trends”) using local polynomial regression (the `loess` function from R’s `stats` package):

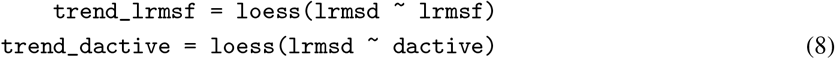

Here, `lrmsd`, `lrmsf`, and `dactive` correspond to their mathematical counterparts `lRMSD`, `lRMSF`, and *d*, respectively.

### 4.6 SCAM models of structural divergence

For each enzyme family, we fit three shape-constrained additive models using the `scam` function from R package `scam`:

*M*_1_: represents only non-functional constraints:

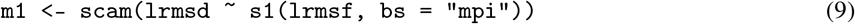

*M*_2_: represents only functional constraints:

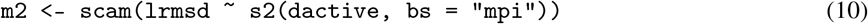

*M*_12_: combines non-functional and functional constraints through functions *s*_1_ and *s*_2_:

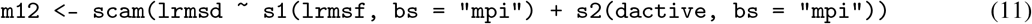

The models use monotone-increasing P-splines (`bs = “mpi”`), consistent with our expectations that structural divergence increases as non-functional constraints weaken with increasing flexibility, and as functional constraints weaken with increasing distance from the active site.

### 4.7 Goodness of Fit Assessment

We assess model performance using the Akaike information criterion AIC (a penalized likelihood criterion), explained deviance (a generalization of *R*^2^ for non-linear models), and RMSE between predicted and observed trends. AIC and explained deviance are calculated from residuals between observed and predicted `lRMSD` values, as provided by the `scam` function used to fit the models. For a given model, RMSE(flex) is the root mean square error between observed and predicted divergence-flexibility trends, defined via LOESS fits (Eq.8). Similarly, RMSE(dist) is the RMSE between observed and predicted divergence-distance trends. To compute these, we use the R function `loess` to fit curves to observed and predicted `lRMSD` values and calculate the RMSE between them.

### 4.8 Shapley decomposition

To quantify the relative contributions of non-functional and functional constraints to structural divergence, we applied Shapley’s method to the three empirical models (*M*_1_, *M*_2_, and *M*_12_). This method allocates the explained deviance between functions *s*_1_ and *s*_2_, accounting for both their individual and combined effects [46].

For each enzyme family, we calculated the Shapley contributions (*SC*) of *s*_1_ and *s*_2_ using the following formulas:

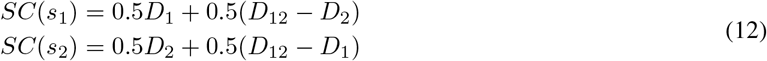

where *D*_1_, *D*_2_, and *D*_12_ represent the deviance explained by *M*_1_ (non-functional constraints through *s*_1_), *M*_2_ (functional constraints through *s*_2_), and *M*_12_ (both constraint functions).

Finally, we calculated the relative Shapley contributions (*RSC*):

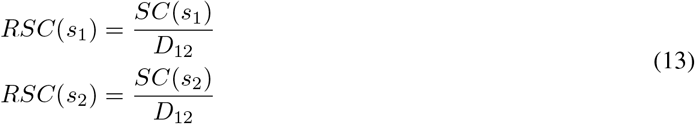

These contributions sum to 1 and represent the relative importance of non-functional and functional constraints in explaining structural divergence.

### 4.9 Determinants of Shapley contributions

To identify the determinants of Shapley contributions *SC* and *RSC*, we used bidirectional stepwise linear regression, using the `step` function from base R. For each family’s reference enzyme, we considered several potential predictors: protein size, the distribution of residue flexibilities (mean, standard deviation, range), the distribution of distances to the active site (mean, standard deviation, range), and the correlation between flexibility and distance (*ρ*(`lRMSF`, *d*)). We also examined two family-level characteristics: the number of homologous proteins and the overall structural divergence. The stepwise regression identified the most relevant predictors among these properties, allowing us to determine which factors best account for the variation in Shapley contributions across enzyme families.

## 5 Data availability

All processed data and code needed to reproduce the figures and tables in this paper are available at https://github.com/jechave/enzyme_evo_str_analysis. Additional figures and analyses are provided in Supplementary Material.

## Acknowledgments

We thank Ioannis Riziotis for sharing the data of Riziotis et al. [37]. This work was supported by CONICET (grant PIP-11220210100462) and regular ISYEB funding.

