## Supplementary Material for "On the variation of structural divergence among residues in enzyme evolution"

### 1. Dataset

Table S1: Dataset entries and reference protein

| M-CSA | PDB | Name | Source | Family |
|---|---|---|---|---|
| 2 | 1btl_A | Beta-lactamase TEM | Escherichia coli | Class-A beta-lactamase family |
| 15 | 1znb_A | Metallo-beta-lactamase type 2 | Bacteroides fragilis | Metallo-beta-lactamase superfamily |
| 71 | 1eug_A | Uracil-DNA glycosylase | Escherichia coli B | Uracil-DNA glycosylase (UDG) superfamily |
| 98 | 1bsz_C | Peptide deformylase | Escherichia coli K-12 | Polypeptide deformylase family |
| 109 | 1d3g_A | Dihydroorotate dehydrogenase (quinone), mitochondrial | Homo sapiens | Dihydroorotate dehydrogenase family |
| 148 | 1onr_A | Name not found | Escherichia coli | Transaldolase family |
| 151 | 1q0n_A | 2-amino-4-hydroxy-6-hydroxymethyldihydropteridine pyrophosphokinase | Escherichia coli | HPPK family |
| 160 | 1ako_A | Exodeoxyribonuclease III | Escherichia coli K-12 | DNA repair enzymes AP/ExoA family |
| 164 | 1ruv_A | Ribonuclease pancreatic | Bos taurus | Pancreatic ribonuclease family |
| 174 | 9pap_A | Papain | Carica papaya | Peptidase C1 family |
| 216 | 1ca2_A | Carbonic anhydrase 2 | Homo sapiens | Alpha-carbonic anhydrase family |
| 252 | 1igs_A | Indole-3-glycerol phosphate synthase | Saccharolobus solfataricus | TrpC family |
| 257 | 1xx2_A | Beta-lactamase | Enterobacter cloacae | Class-C beta-lactamase family |
| 258 | 1sml_A | Metallo-beta-lactamase L1 type 3 | Stenotrophomonas maltophilia | Metallo-beta-lactamase superfamily |
| 290 | 1zio_A | Adenylate kinase | Geobacillus stearothermophilus | Adenylate kinase family |
| 328 | 1lbm_A | N-(5&apos | Thermotoga maritima | TrpF family |
| 351 | 1snz_A | Galactose mutarotase | Homo sapiens | Aldose epimerase family |
| 362 | 1d6o_A | Peptidyl-prolyl cis-trans isomerase FKBP1A | Homo sapiens | FKBP-type PPIase family |
| 376 | 1a4l_A | Adenosine deaminase | Mus musculus | Metallo-dependent hydrolases superfamily |
| 394 | 1aj0_A | Dihydropteroate synthase | Escherichia coli | DHPS family |
| 444 | 1cel_A | Exoglucanase 1 | Trichoderma reesei | Glycosyl hydrolase 7 (cellulase C) family |
| 462 | 1pnt_A | Low molecular weight phosphotyrosine protein phosphatase | Bos taurus | Low molecular weight phosphotyrosine protein phosphatase family |
| 467 | 1cv2_A | Haloalkane dehalogenase | Sphingomonas paucimobilis | Haloalkane dehalogenase family |
| 480 | 1czf_A | Endopolygalacturonase II | Aspergillus niger | Glycosyl hydrolase 28 family |
| 597 | 1uch_A | Ubiquitin carboxyl-terminal hydrolase isozyme L3 | Homo sapiens | Peptidase C12 family |
| 681 | 1gq8_A | Pectinesterase | Daucus carota | Pectinesterase family |
| 693 | 2rnf_A | Ribonuclease 4 | Homo sapiens | Pancreatic ribonuclease family |
| 749 | 1nml_A | No information found | Marinobacter nauticus | No information found |
| 814 | 1glo_A | Cathepsin S | Homo sapiens | Peptidase C1 family |
| 858 | 1mrq_A | Aldo-keto reductase family 1 member C1 | Homo sapiens | Aldo/keto reductase family |
| 877 | 1pbg_A | 6-phospho-beta-galactosidase | Lactococcus lactis | Glycosyl hydrolase 1 family |
| 908 | 1rtu_A | Ribonuclease U2 | Ustilago sphaerogena | Ribonuclease U2 family |
| 923 | 2acy_A | Acylphosphatase-1 | Bos taurus | Acylphosphatase family |
| 931 | 2pth_A | Peptidyl-tRNA hydrolase | Escherichia coli K-12 | PTH family |

Table S2: Dataset properties

| M-CSA | N | <Id%> | <RMSD> | Length | AS size | AS RMSD | CATH | EC |
|---|---|---|---|---|---|---|---|---|
| 2 | 6 | 43 | 1.04 | 263 | 6 | 1.78 | alpha/beta | Hydrolases |
| 15 | 6 | 35 | 1.16 | 225 | 8 | 1.84 | alpha/beta | Hydrolases |
| 71 | 8 | 43 | 1.08 | 223 | 4 | 1.49 | alpha/beta | Hydrolases |
| 98 | 10 | 37 | 2.15 | 168 | 7 | 1.44 | alpha/beta | Hydrolases |
| 109 | 4 | 44 | 1.31 | 359 | 7 | 2.65 | alpha/beta | Oxidoreductases |
| 148 | 5 | 53 | 1.06 | 316 | 5 | 2.09 | alpha/beta | Transferases |
| 151 | 4 | 47 | 1.14 | 158 | 4 | 2.07 | alpha/beta | Transferases |
| 160 | 10 | 34 | 1.49 | 256 | 7 | 2.63 | alpha/beta | Hydrolases |
| 164 | 5 | 48 | 1.09 | 123 | 5 | 1.92 | alpha/beta | Lyases |
| 174 | 12 | 43 | 1.01 | 212 | 4 | 1.57 | alpha/beta | Hydrolases |
| 216 | 5 | 39 | 1.09 | 256 | 6 | 2.27 | alpha/beta | Lyases |
| 252 | 6 | 35 | 1.75 | 245 | 7 | 2.93 | alpha/beta | Lyases |
| 257 | 10 | 45 | 0.94 | 359 | 6 | 1.48 | alpha/beta | Hydrolases |
| 258 | 4 | 38 | 1.97 | 263 | 7 | 1.80 | alpha/beta | Hydrolases |
| 290 | 7 | 47 | 1.10 | 217 | 5 | 1.82 | alpha/beta | Transferases |
| 328 | 4 | 33 | 1.47 | 193 | 2 | 2.55 | alpha/beta | Isomerases |
| 351 | 4 | 33 | 1.36 | 341 | 3 | 2.65 | all-beta | Isomerases |
| 362 | 7 | 46 | 0.92 | 107 | 6 | 2.30 | alpha/beta | Isomerases |
| 376 | 4 | 44 | 1.30 | 349 | 6 | 2.42 | alpha/beta | Hydrolases |
| 394 | 10 | 40 | 2.02 | 280 | 2 | 2.35 | alpha/beta | Transferases |
| 444 | 6 | 46 | 1.19 | 431 | 4 | 2.12 | all-beta | Hydrolases |
| 462 | 6 | 37 | 1.15 | 155 | 6 | 1.81 | alpha/beta | Hydrolases |
| 467 | 5 | 45 | 1.03 | 293 | 5 | 1.90 | alpha/beta | Hydrolases |
| 480 | 4 | 44 | 1.05 | 335 | 6 | 1.86 | all-beta | Hydrolases |
| 597 | 4 | 38 | 1.76 | 204 | 4 | 3.15 | alpha/beta | Hydrolases |
| 681 | 4 | 31 | 2.88 | 300 | 5 | 6.68 | all-beta | Hydrolases |
| 693 | 7 | 38 | 1.62 | 120 | 3 | 2.20 | alpha/beta | Hydrolases |
| 749 | 6 | 56 | 1.96 | 316 | 1 | 3.89 | all-alpha | NA |
| 814 | 4 | 46 | 0.72 | 215 | 4 | 1.22 | alpha/beta | Hydrolases |
| 858 | 16 | 38 | 1.88 | 322 | 4 | 1.70 | alpha/beta | Oxidoreductases |
| 877 | 23 | 37 | 1.52 | 447 | 2 | 2.42 | alpha/beta | Hydrolases |
| 908 | 5 | 45 | 1.45 | 107 | 4 | 2.55 | alpha/beta | Lyases |
| 923 | 4 | 34 | 0.99 | 98 | 2 | 1.29 | alpha/beta | Hydrolases |
| 931 | 7 | 39 | 1.40 | 193 | 4 | 1.85 | alpha/beta | Hydrolases |

N: number of family members; <Id%>: average family seq. identity; <RMSD>: average family RMSD; Length: number of residues of reference protein; AS size: number of residues of active site; CATH: CATH class; EC: EC class

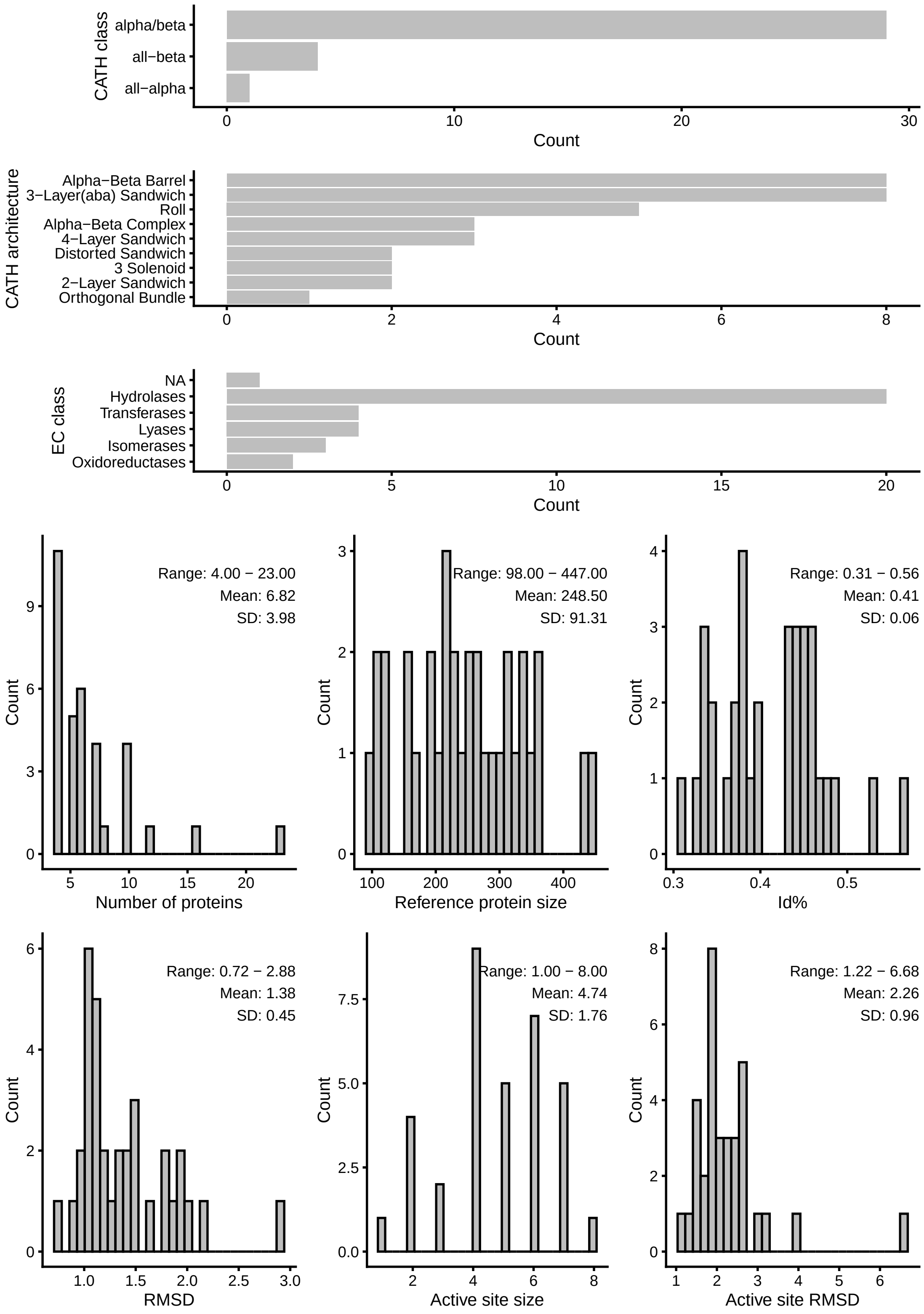

Figure S1: Distribution of properties of dataset families.

#### 2. Correlation between flexibility and distance

The following figure shows the distribution of spearman correlations between flexibility (lRMSF) and distance to the active site (d) (panel a). The correlation between these variables depends on the location of the active site within the enzyme structure. In most cases active sites are located in relatively rigid regions. Therefore, as one moves away from the active site, not only distance increases but flexibility too, which originates the correlation (panel b).

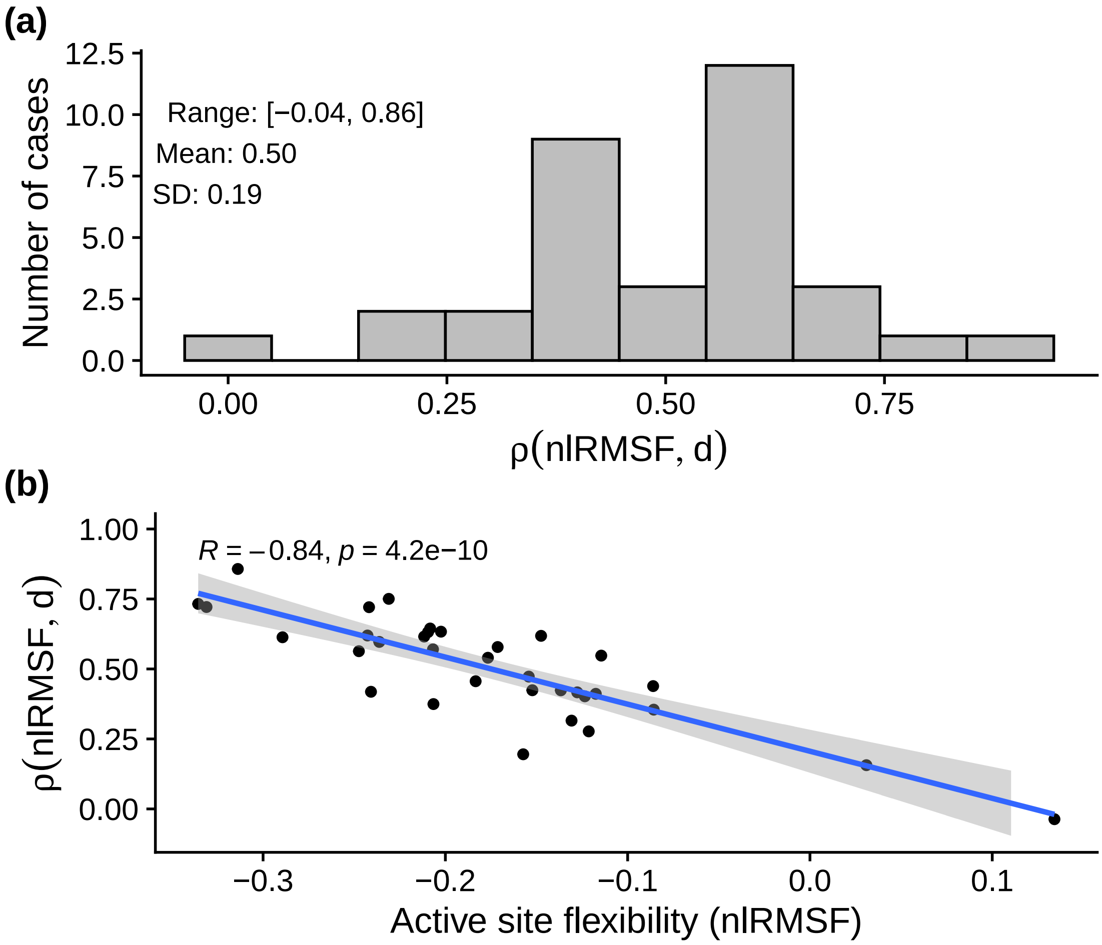

Figure S2: Correlation betwen flexibility and distance.

### 3. Model performance

The following table presents the goodness-of-fit metrics for the three models tested: $M_1$ (flexibility only), $M_2$ (distance only), and $M_{12}$ (both predictors). For each enzyme family, we report the Akaike Information Criterion (AIC), the deviance explained, the root mean square error between observed and predicted divergence-flexibility trends (RMSE(flex)), the root mean square error between observed and predicted divergence-distance trends (RMSE(dist)), and the relative Shapley contribution of functional constraints to structural divergence ($RSC(s_2)$). Families are sorted by increasing $RSC(s_2)$.

Table S3: Variation of model performance across all families

| | | | | AIC | | | Expl. Dev. | | | RMSE(flex) | | | RMSE(dist) | | |
|---|---|---|---|---|---|---|---|---|---|---|---|---|---|---|---|
| M-CSA | PDB | RSC($s_2$) | Best | $M_1$ | $M_2$ | $M_{12}$ | $M_1$ | $M_2$ | $M_{12}$ | $M_1$ | $M_2$ | $M_{12}$ | $M_1$ | $M_2$ | $M_{12}$ |
| 749 | 1nml_A | -0.00 | $M_1$ | 714 | 799 | 714 | 0.25 | -0.00 | 0.25 | 0.06 | 0.44 | 0.06 | 0.28 | 0.25 | 0.28 |
| 597 | 1uch_A | 0.04 | $M_1$ | 346 | 436 | 348 | 0.38 | 0.03 | 0.38 | 0.05 | 0.39 | 0.05 | 0.11 | 0.10 | 0.11 |
| 858 | 1mrq_A | 0.13 | $M_1$ | 645 | 797 | 647 | 0.45 | 0.11 | 0.45 | 0.05 | 0.41 | 0.05 | 0.19 | 0.08 | 0.19 |
| 258 | 1sml_A | 0.16 | $M_1$ | 409 | 636 | 412 | 0.66 | 0.21 | 0.66 | 0.04 | 0.48 | 0.05 | 0.08 | 0.11 | 0.09 |
| 394 | 1aj0_A | 0.17 | $M_{12}$ | 480 | 660 | 463 | 0.57 | 0.17 | 0.60 | 0.03 | 0.51 | 0.02 | 0.15 | 0.07 | 0.05 |
| 467 | 1cv2_A | 0.20 | $M_1$ | 550 | 634 | 550 | 0.36 | 0.14 | 0.36 | 0.04 | 0.29 | 0.04 | 0.14 | 0.07 | 0.14 |
| 681 | 1gq8_A | 0.21 | $M_1$ | 668 | 782 | 673 | 0.45 | 0.19 | 0.44 | 0.05 | 0.45 | 0.08 | 0.21 | 0.17 | 0.21 |
| 151 | 1q0n_A | 0.24 | $M_{12}$ | 224 | 308 | 205 | 0.57 | 0.25 | 0.61 | 0.02 | 0.42 | 0.03 | 0.22 | 0.10 | 0.17 |
| 109 | 1d3g_A | 0.24 | $M_{12}$ | 701 | 819 | 688 | 0.41 | 0.19 | 0.44 | 0.04 | 0.37 | 0.04 | 0.14 | 0.09 | 0.09 |
| 290 | 1zio_A | 0.26 | $M_{12}$ | 293 | 403 | 275 | 0.56 | 0.27 | 0.60 | 0.04 | 0.34 | 0.04 | 0.14 | 0.03 | 0.04 |
| 257 | 1xx2_A | 0.27 | $M_{12}$ | 367 | 527 | 358 | 0.54 | 0.28 | 0.56 | 0.01 | 0.24 | 0.01 | 0.06 | 0.03 | 0.03 |
| 71 | 1eug_A | 0.27 | $M_{12}$ | 153 | 327 | 144 | 0.72 | 0.39 | 0.73 | 0.03 | 0.27 | 0.03 | 0.07 | 0.03 | 0.04 |
| 98 | 1bsz_C | 0.28 | $M_{12}$ | 288 | 410 | 287 | 0.71 | 0.39 | 0.71 | 0.08 | 0.40 | 0.07 | 0.13 | 0.07 | 0.09 |
| 164 | 1ruv_A | 0.30 | $M_{12}$ | 264 | 297 | 252 | 0.40 | 0.21 | 0.47 | 0.08 | 0.45 | 0.08 | 0.23 | 0.09 | 0.07 |
| 877 | 1pbg_A | 0.31 | $M_{12}$ | 784 | 904 | 783 | 0.45 | 0.27 | 0.45 | 0.03 | 0.24 | 0.03 | 0.06 | 0.03 | 0.05 |
| 328 | 1lbm_A | 0.31 | $M_{12}$ | 306 | 338 | 298 | 0.30 | 0.17 | 0.35 | 0.05 | 0.22 | 0.06 | 0.11 | 0.05 | 0.05 |
| 480 | 1czf_A | 0.33 | $M_{12}$ | 566 | 618 | 556 | 0.29 | 0.18 | 0.31 | 0.04 | 0.24 | 0.05 | 0.11 | 0.06 | 0.07 |
| 216 | 1ca2_A | 0.34 | $M_{12}$ | 518 | 571 | 505 | 0.39 | 0.25 | 0.43 | 0.04 | 0.30 | 0.04 | 0.16 | 0.06 | 0.07 |
| 931 | 2pth_A | 0.35 | $M_{12}$ | 366 | 403 | 336 | 0.42 | 0.27 | 0.49 | 0.04 | 0.37 | 0.07 | 0.24 | 0.08 | 0.07 |
| 15 | 1znb_A | 0.37 | $M_{12}$ | 336 | 396 | 286 | 0.51 | 0.35 | 0.61 | 0.08 | 0.32 | 0.06 | 0.21 | 0.06 | 0.04 |
| 160 | 1ako_A | 0.40 | $M_{12}$ | 414 | 459 | 389 | 0.49 | 0.38 | 0.54 | 0.04 | 0.24 | 0.04 | 0.15 | 0.08 | 0.08 |
| 693 | 2rnf_A | 0.40 | $M_{12}$ | 243 | 261 | 218 | 0.39 | 0.29 | 0.51 | 0.08 | 0.37 | 0.09 | 0.31 | 0.15 | 0.11 |
| 148 | 1onr_A | 0.45 | $M_{12}$ | 587 | 619 | 575 | 0.47 | 0.41 | 0.50 | 0.06 | 0.13 | 0.04 | 0.11 | 0.05 | 0.06 |
| 462 | 1pnt_A | 0.46 | $M_{12}$ | 233 | 252 | 221 | 0.56 | 0.51 | 0.61 | 0.06 | 0.16 | 0.05 | 0.13 | 0.04 | 0.06 |
| 814 | 1glo_A | 0.47 | $M_{12}$ | 252 | 263 | 212 | 0.35 | 0.32 | 0.47 | 0.04 | 0.18 | 0.03 | 0.16 | 0.03 | 0.02 |
| 351 | 1snz_A | 0.49 | $M_{12}$ | 697 | 701 | 654 | 0.34 | 0.33 | 0.43 | 0.04 | 0.25 | 0.08 | 0.19 | 0.04 | 0.04 |
| 2 | 1btl_A | 0.49 | $M_{12}$ | 448 | 455 | 411 | 0.36 | 0.35 | 0.46 | 0.04 | 0.18 | 0.04 | 0.20 | 0.04 | 0.04 |
| 376 | 1a4l_A | 0.50 | $M_{12}$ | 781 | 778 | 768 | 0.17 | 0.17 | 0.20 | 0.06 | 0.13 | 0.08 | 0.13 | 0.06 | 0.04 |
| 923 | 2acy_A | 0.51 | $M_{12}$ | 199 | 197 | 177 | 0.41 | 0.42 | 0.54 | 0.09 | 0.28 | 0.10 | 0.29 | 0.12 | 0.09 |
| 174 | 9pap_A | 0.52 | $M_{12}$ | 334 | 333 | 315 | 0.24 | 0.25 | 0.31 | 0.04 | 0.16 | 0.06 | 0.15 | 0.05 | 0.05 |
| 444 | 1cel_A | 0.57 | $M_{12}$ | 820 | 791 | 756 | 0.27 | 0.32 | 0.38 | 0.05 | 0.15 | 0.04 | 0.21 | 0.04 | 0.04 |
| 252 | 1igs_A | 0.63 | $M_{12}$ | 436 | 377 | 353 | 0.37 | 0.51 | 0.56 | 0.04 | 0.14 | 0.04 | 0.26 | 0.04 | 0.04 |
| 362 | 1d6o_A | 0.68 | $M_{12}$ | 189 | 153 | 105 | 0.20 | 0.43 | 0.64 | 0.08 | 0.32 | 0.11 | 0.44 | 0.11 | 0.08 |
| 908 | 1rtu_A | 0.85 | $M_2$ | 249 | 180 | 181 | 0.16 | 0.56 | 0.57 | 0.09 | 0.08 | 0.05 | 0.48 | 0.07 | 0.07 |

RSC($s_2$): relative Shapley contribution of functional constraints to structural divergence; Best: model with lowest AIC; Expl. Dev.: deviance explained by each model; RMSE(flex): root mean square error between observed and predicted divergence-flexibility trends; RMSE(dist): RMSE between observed and predicted divergence-distance trends.

### 4. Robustness analysis

To assess the robustness of our results, we tested whether the model performance metrics are sensitive to methodological choices. Specifically, we examined robustness with respect to: (1) reference protein choice, comparing results when using the reference protein (ca_ref) versus the structurally closest homolog (ca_hom, selected based on lowest overall RMSD to reduce artifacts from gaps and non-evolutionary structural differences); and (2) atom choice, comparing results when using C$\alpha$ atoms (ca_ref) versus C$\beta$ atoms (cb_ref) for structural calculations.

Results show that the choice of reference protein has minimal impact: goodness-of-fit metrics exhibit strong correlations between the two approaches, and average values are nearly identical, confirming robustness to this methodological choice. In contrast, while C$\alpha$- and C$\beta$-based analyses show significant correlations, the correlations are weaker and C$\beta$-based analyses consistently yield lower goodness-of-fit values across all metrics, supporting our choice to use C$\alpha$ atoms for the main analyses.

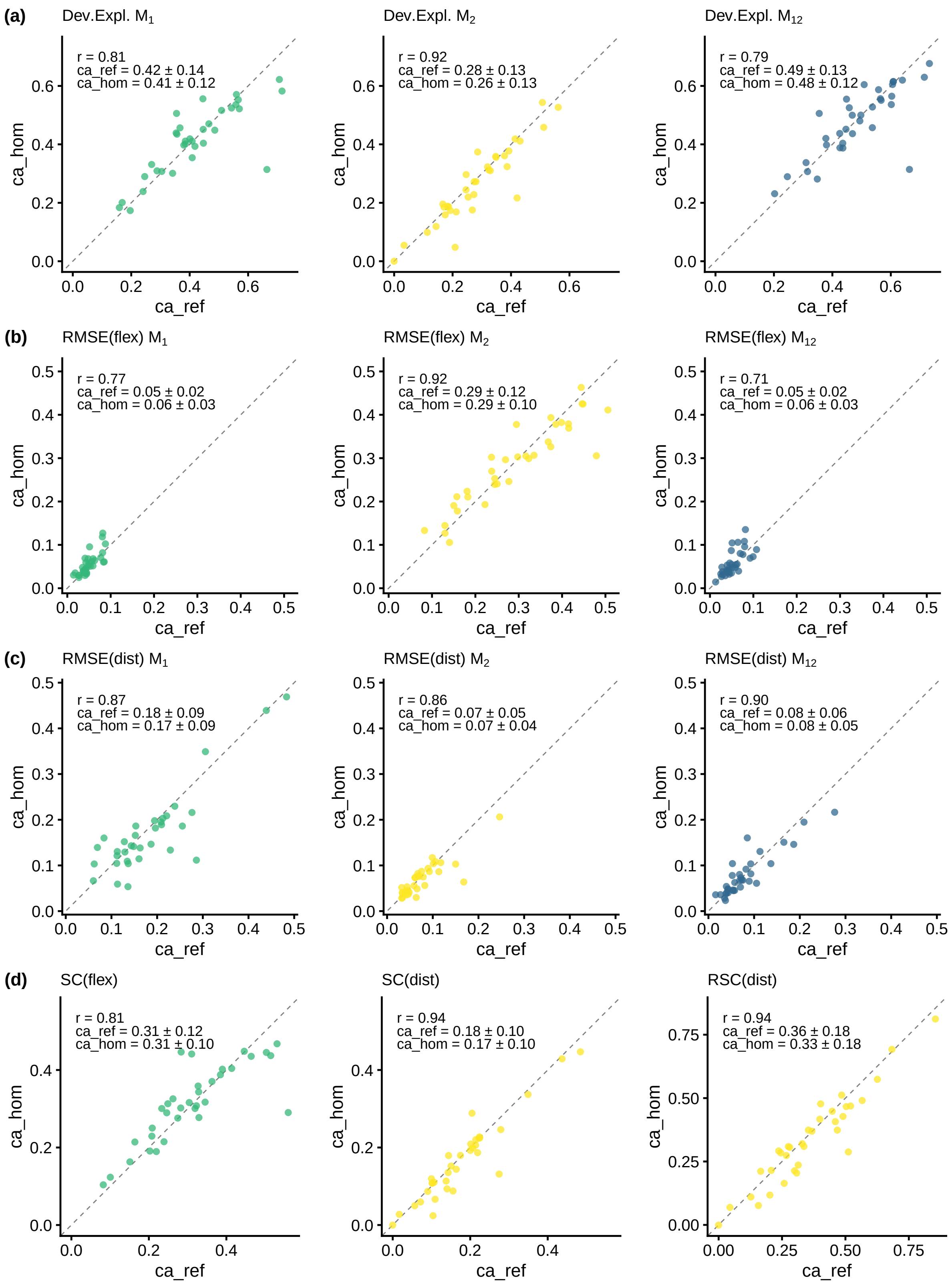

Figure S3: Robustness to reference protein choice. Each panel compares goodness-of-fit metrics between analyses using the reference protein (ca_ref) and a homolog (ca_hom). (a) Deviance explained for models M1, M2, and M12. (b) RMSE between observed and predicted divergence-flexibility trends. (c) RMSE between observed and predicted divergence-distance trends. (d) Shapley contributions: SC(flex), SC(dist), and RSC(dist).

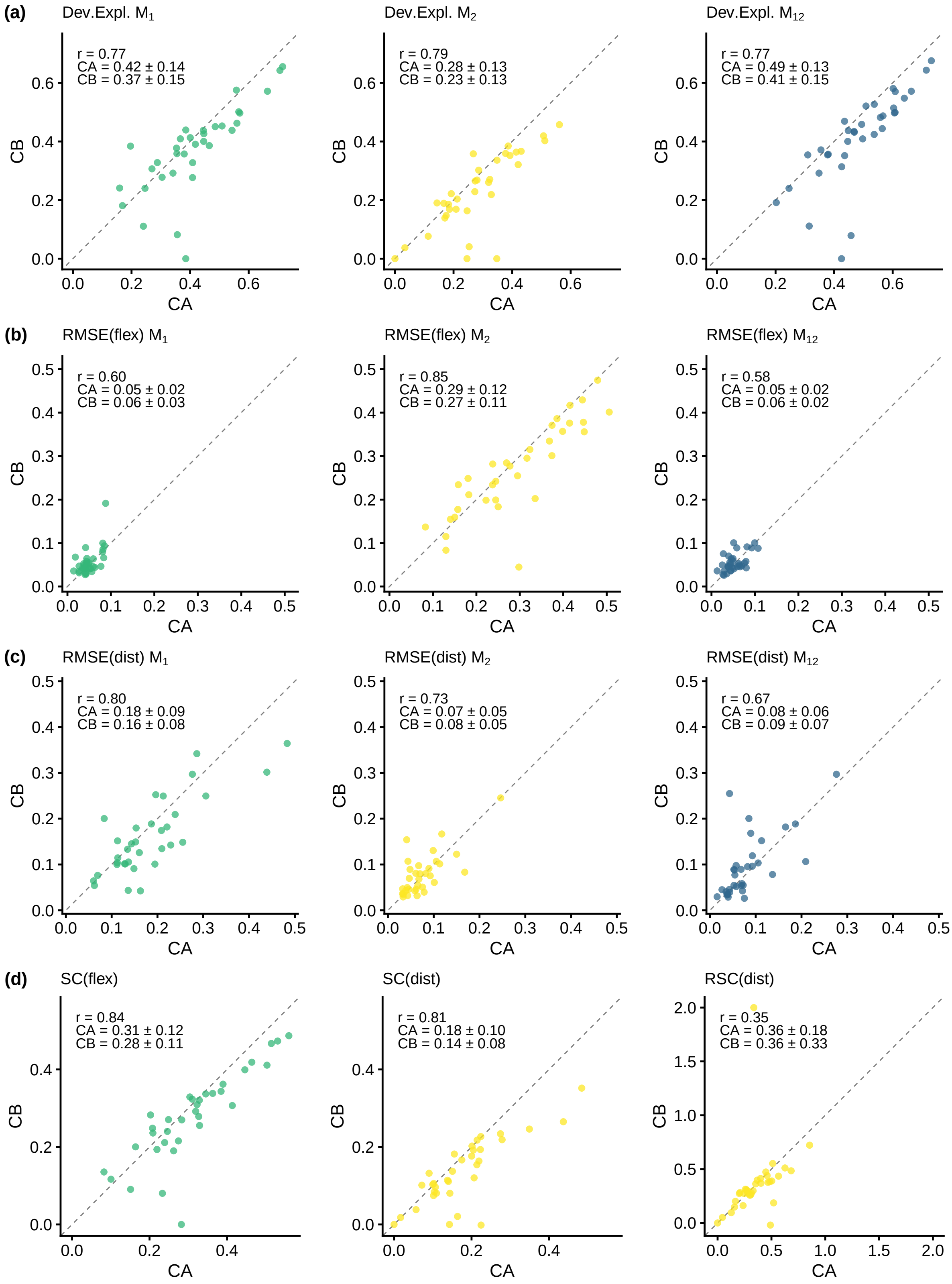

Figure S4: Robustness to atom choice. Each panel compares goodness-of-fit metrics between analyses using C$\alpha$ atoms (CA) and C$\beta$ atoms (CB). (a) Deviance explained for models M1, M2, and M12. (b) RMSE between observed and predicted divergence-flexibility trends. (c) RMSE between observed and predicted divergence-distance trends. (d) Shapley contributions: SC(flex), SC(dist), and RSC(dist).

#### 5. Active site conservation

We analyze structural divergence at active site residues ($d = 0$). For each family, divergence metrics are family-normalized, and $M_{12}$ predictions are decomposed into per-residue $s_1$ (non-functional constraint) and $s_2$ (functional constraint) contributions. For each active site residue, we calculate the relative magnitude $rs_1 = |s_1|/(|s_1| + |s_2|)$ and $rs_2 = |s_2|/(|s_1| + |s_2|)$. The values reported in the tables below ($s_1$, $s_2$, $rs_1$, $rs_2$) are the means of these per-residue metrics over all active site residues within each family.

Table S4: Summary statistics for active site divergence

| Statistic | Obs | $M_{12}$ | $s_1$ | $s_2$ | $rs_1$ | $rs_2$ |
|---|---|---|---|---|---|---|
| Mean | -0.62 | -0.69 | -0.32 | -0.38 | 0.52 | 0.48 |
| SD | 0.28 | 0.22 | 0.20 | 0.28 | 0.29 | 0.29 |
| Min | -1.07 | -1.06 | -0.78 | -1.03 | 0.03 | 0.00 |
| Max | 0.25 | -0.26 | 0.15 | -0.00 | 1.00 | 0.97 |

Obs: observed nlRMSD; $M_{12}$: predicted nlRMSD; $s_1$: non-functional constraint contribution; $s_2$: functional constraint contribution; $rs_1$, $rs_2$: relative magnitude contributions. Statistics calculated across 33 enzyme families (case 749 excluded). All values are family-averaged active site metrics ($d = 0$).

Table S5: Active site structural divergence and constraint contributions

| | Whole-protein | | | Active site | | | | | | |
|---|---|---|---|---|---|---|---|---|---|---|
| M-CSA | SC($s_1$) | SC($s_2$) | RSC($s_2$) | nlRMSF | Obs | $M_{12}$ | $s_1$ | $s_2$ | $rs_1$ | $rs_2$ |
| 597 | 0.36 | 0.02 | 0.04 | -0.21 | -0.40 | -0.38 | -0.38 | -0.00 | 1.00 | 0.00 |
| 858 | 0.39 | 0.06 | 0.13 | -0.21 | -0.68 | -0.57 | -0.57 | -0.00 | 1.00 | 0.00 |
| 258 | 0.56 | 0.10 | 0.16 | -0.16 | -0.37 | -0.26 | -0.26 | -0.00 | 1.00 | 0.00 |
| 394 | 0.50 | 0.10 | 0.17 | -0.12 | -0.89 | -0.88 | -0.32 | -0.56 | 0.32 | 0.68 |
| 467 | 0.28 | 0.07 | 0.20 | -0.20 | -0.54 | -0.53 | -0.53 | -0.00 | 1.00 | 0.00 |
| 681 | 0.35 | 0.09 | 0.21 | -0.24 | -0.76 | -0.75 | -0.75 | -0.00 | 1.00 | 0.00 |
| 151 | 0.46 | 0.14 | 0.24 | -0.18 | -0.04 | -0.39 | -0.29 | -0.09 | 0.72 | 0.28 |
| 109 | 0.33 | 0.11 | 0.24 | -0.09 | 0.25 | -0.29 | -0.16 | -0.13 | 0.63 | 0.37 |
| 290 | 0.45 | 0.16 | 0.26 | -0.18 | -0.42 | -0.38 | -0.22 | -0.15 | 0.69 | 0.31 |
| 257 | 0.41 | 0.15 | 0.27 | -0.21 | -0.60 | -0.64 | -0.36 | -0.28 | 0.54 | 0.46 |
| 71 | 0.53 | 0.20 | 0.27 | -0.25 | -0.47 | -0.49 | -0.42 | -0.07 | 0.85 | 0.15 |
| 98 | 0.51 | 0.20 | 0.28 | -0.34 | -0.93 | -0.96 | -0.78 | -0.18 | 0.81 | 0.19 |
| 164 | 0.33 | 0.14 | 0.30 | -0.13 | -0.42 | -0.67 | -0.20 | -0.47 | 0.35 | 0.65 |
| 877 | 0.31 | 0.14 | 0.31 | -0.23 | -0.81 | -0.73 | -0.51 | -0.22 | 0.70 | 0.30 |
| 328 | 0.24 | 0.11 | 0.31 | -0.15 | -0.61 | -0.73 | -0.25 | -0.48 | 0.34 | 0.66 |
| 480 | 0.21 | 0.10 | 0.33 | -0.21 | -0.67 | -0.61 | -0.36 | -0.25 | 0.58 | 0.42 |
| 216 | 0.28 | 0.14 | 0.34 | -0.11 | -0.90 | -0.91 | -0.27 | -0.64 | 0.40 | 0.60 |
| 931 | 0.32 | 0.17 | 0.35 | -0.15 | -0.80 | -0.83 | -0.23 | -0.60 | 0.27 | 0.73 |
| 15 | 0.38 | 0.22 | 0.37 | -0.15 | -0.83 | -0.81 | -0.20 | -0.60 | 0.35 | 0.65 |
| 160 | 0.32 | 0.21 | 0.40 | -0.29 | -1.07 | -1.02 | -0.51 | -0.51 | 0.46 | 0.54 |
| 693 | 0.30 | 0.20 | 0.40 | -0.09 | -0.84 | -1.01 | -0.15 | -0.87 | 0.22 | 0.78 |
| 148 | 0.27 | 0.22 | 0.45 | -0.31 | -0.55 | -0.67 | -0.56 | -0.11 | 0.83 | 0.17 |
| 462 | 0.33 | 0.28 | 0.46 | -0.33 | -0.93 | -0.93 | -0.55 | -0.38 | 0.55 | 0.45 |
| 814 | 0.25 | 0.22 | 0.47 | -0.12 | -0.54 | -0.62 | -0.13 | -0.49 | 0.21 | 0.79 |
| 351 | 0.22 | 0.21 | 0.49 | -0.24 | -0.69 | -0.79 | -0.32 | -0.47 | 0.40 | 0.60 |
| 2 | 0.23 | 0.22 | 0.49 | -0.21 | -0.80 | -0.89 | -0.30 | -0.59 | 0.32 | 0.68 |
| 376 | 0.10 | 0.10 | 0.50 | -0.24 | -0.98 | -0.96 | -0.20 | -0.77 | 0.20 | 0.80 |
| 923 | 0.26 | 0.27 | 0.51 | -0.12 | -0.24 | -0.45 | -0.18 | -0.27 | 0.39 | 0.61 |
| 174 | 0.15 | 0.16 | 0.52 | -0.14 | -0.27 | -0.48 | -0.12 | -0.36 | 0.23 | 0.77 |
| 444 | 0.16 | 0.21 | 0.57 | -0.24 | -0.69 | -0.64 | -0.29 | -0.35 | 0.44 | 0.56 |
| 252 | 0.21 | 0.35 | 0.63 | -0.17 | -0.76 | -0.81 | -0.17 | -0.65 | 0.20 | 0.80 |
| 362 | 0.20 | 0.44 | 0.68 | 0.13 | -0.50 | -0.68 | 0.15 | -0.83 | 0.20 | 0.80 |
| 908 | 0.08 | 0.48 | 0.85 | -0.13 | -0.76 | -1.06 | -0.03 | -1.03 | 0.03 | 0.97 |

Values represent family-averaged metrics for active site residues ($d = 0$). Obs: observed nlRMSD; $M_{12}$: predicted nlRMSD; $s_1$: non-functional constraint contribution; $s_2$: functional constraint contribution; $rs_1$, $rs_2$: relative magnitude contributions; SC($s_1$), SC($s_2$): Shapley constraint contributions (whole-family level); RSC($s_2$): relative Shapley contribution of $s_2$; nlRMSF: mean normalized flexibility at active site. Case 749 excluded as outlier. Families sorted by RSC($s_2$).

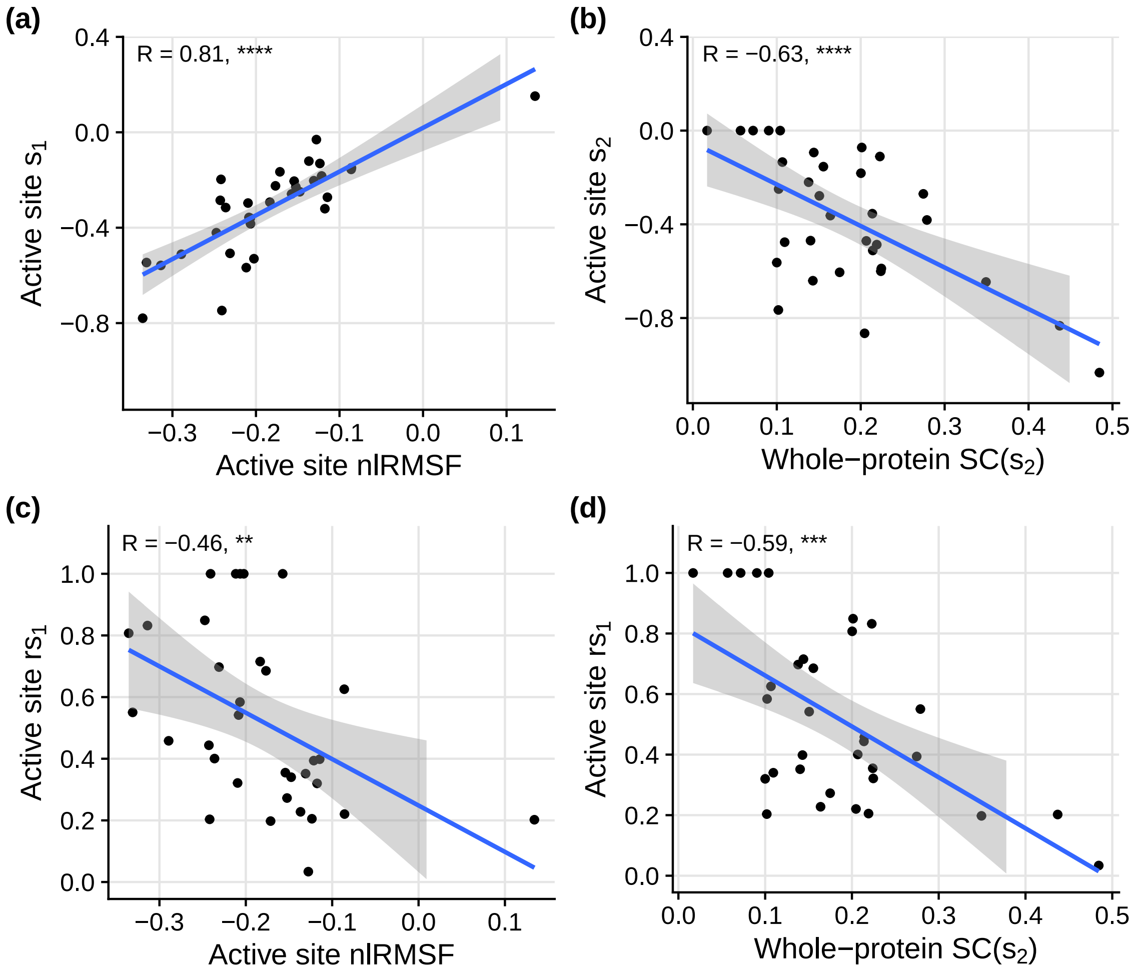

Figure S5: Active site constraint correlations. (a) Relationship between active site non-functional constraint contribution ($s_1$) and active site flexibility (nlRMSF). (b) Relationship between active site functional constraint contribution ($s_2$) and whole-protein Shapley contribution SC($s_2$). (c) Relationship between active site relative non-functional contribution ($rs_1$) and active site flexibility. (d) Relationship between active site relative non-functional contribution ($rs_1$) and whole-protein Shapley contribution SC($s_2$). All metrics are family-averaged over active site residues ($d = 0$). Lines show linear regression fits with 95% confidence intervals. Correlation statistics displayed on each panel.

#### 6. Family by family analysis

Each figure in this section represents a structural divergence analysis for a specific enzyme family, identified by its MCSA ID.

Each figure is organized as follows:

Observed patterns: (a) Residue-specific structural divergence profile, showing normalized log-RMSD (nlRMSD) across residues. (b) Relationship between structural divergence (nlRMSD) and residue flexibility (nlRMSF). (c) Relationship between structural divergence (nlRMSD) and distance from the active site (d).

Model predictions vs. observations: (d) Comparison of observed nlRMSD profile with predictions from models M1, M2, and M12. (e) Flexibility trends: observed data vs. model predictions for nlRMSD vs. nlRMSF. (f) Distance trends: observed data vs. model predictions for nlRMSD vs. d.

Decomposition into flexibility and distance contributions: (g) Profile of structural divergence decomposed into flexibility (s1) and distance (s2) components. (h) Flexibility (s1) component of structural divergence. (i) Distance (s2) component of structural divergence.

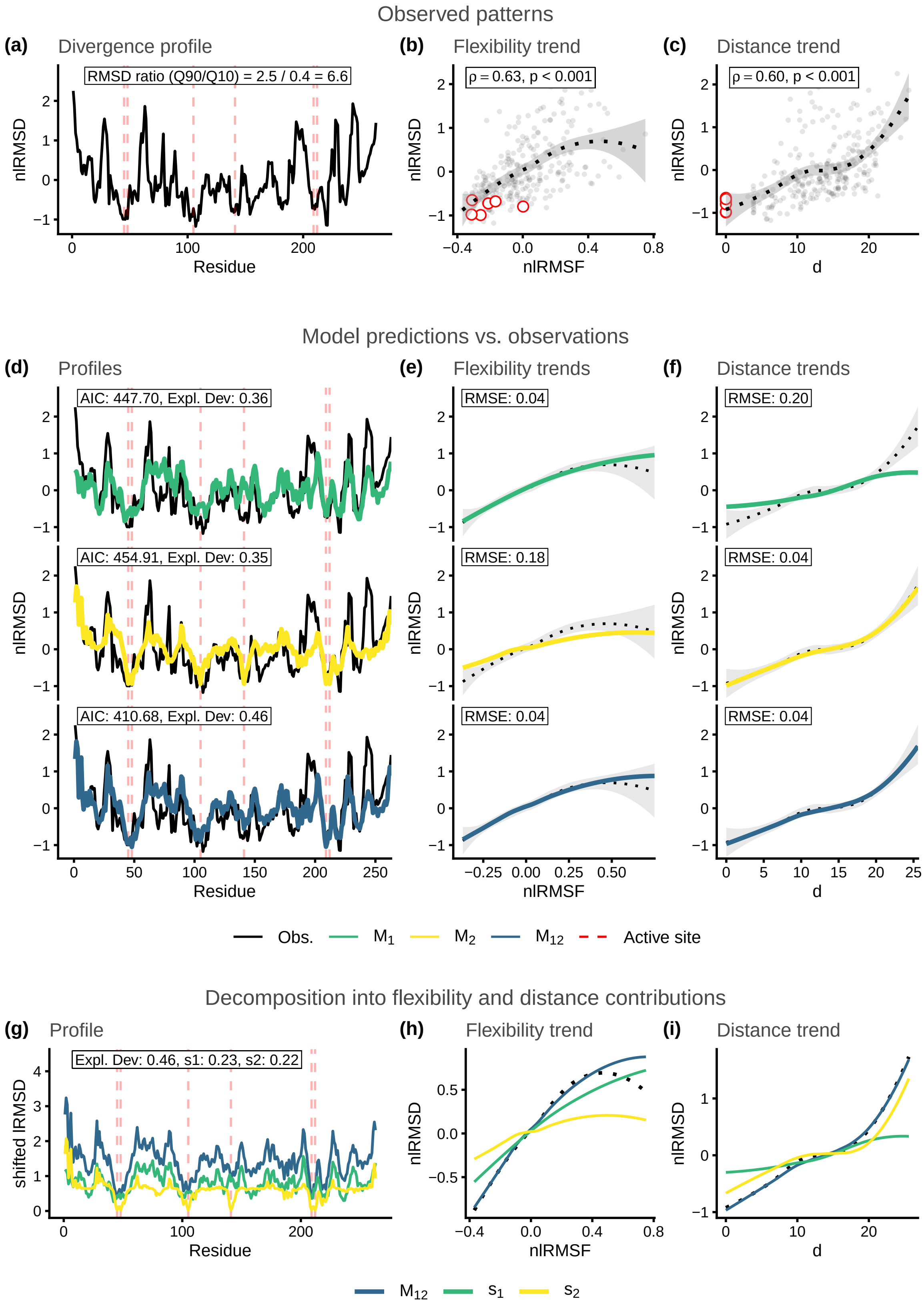

Figure S6: Structural divergence analysis for enzyme family MCSA ID: 2. Reference protein PDB ID: 1btl_A.

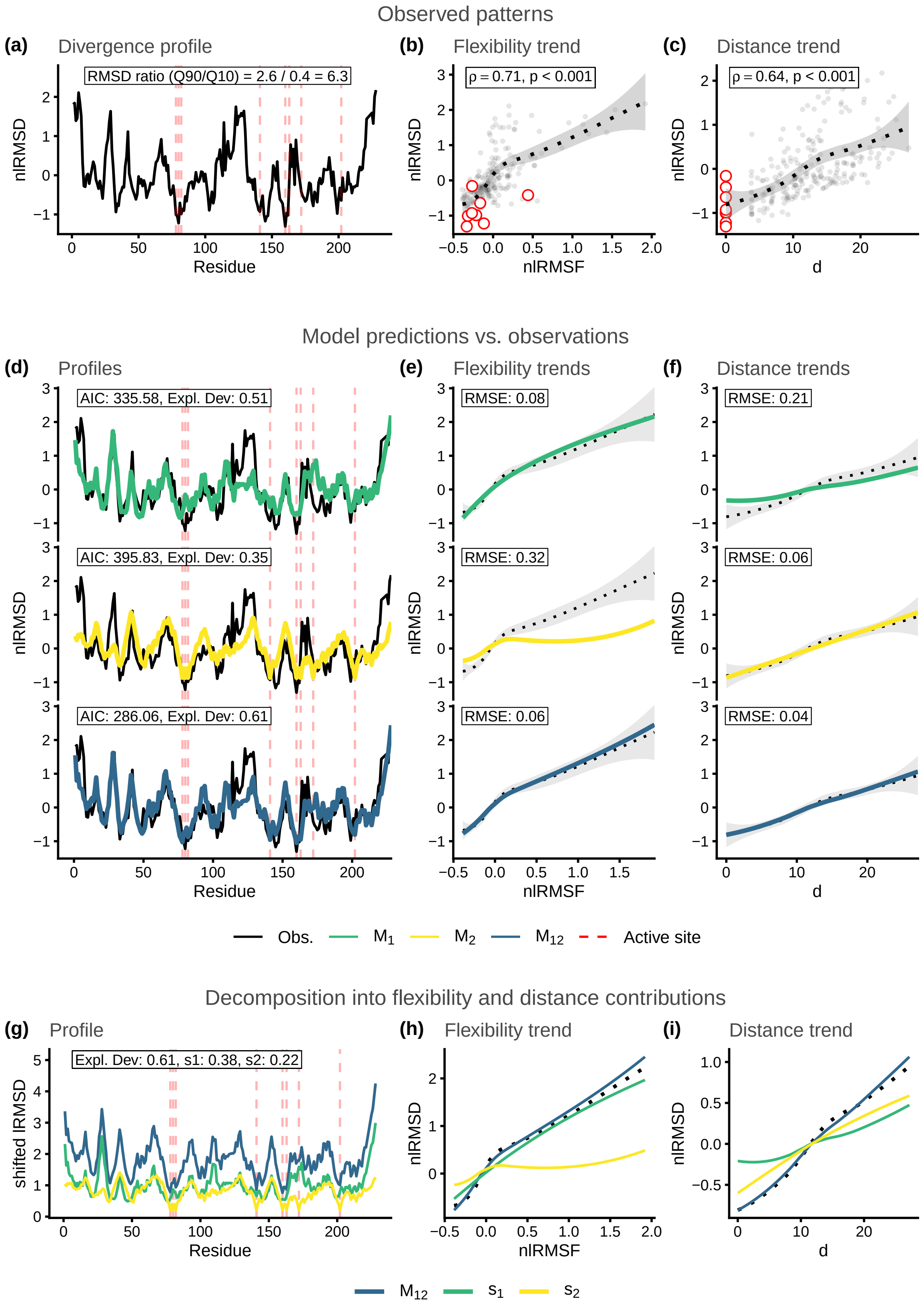

Figure S7: Structural divergence analysis for enzyme family MCSA ID: 15. Reference protein PDB ID: 1znb_A.

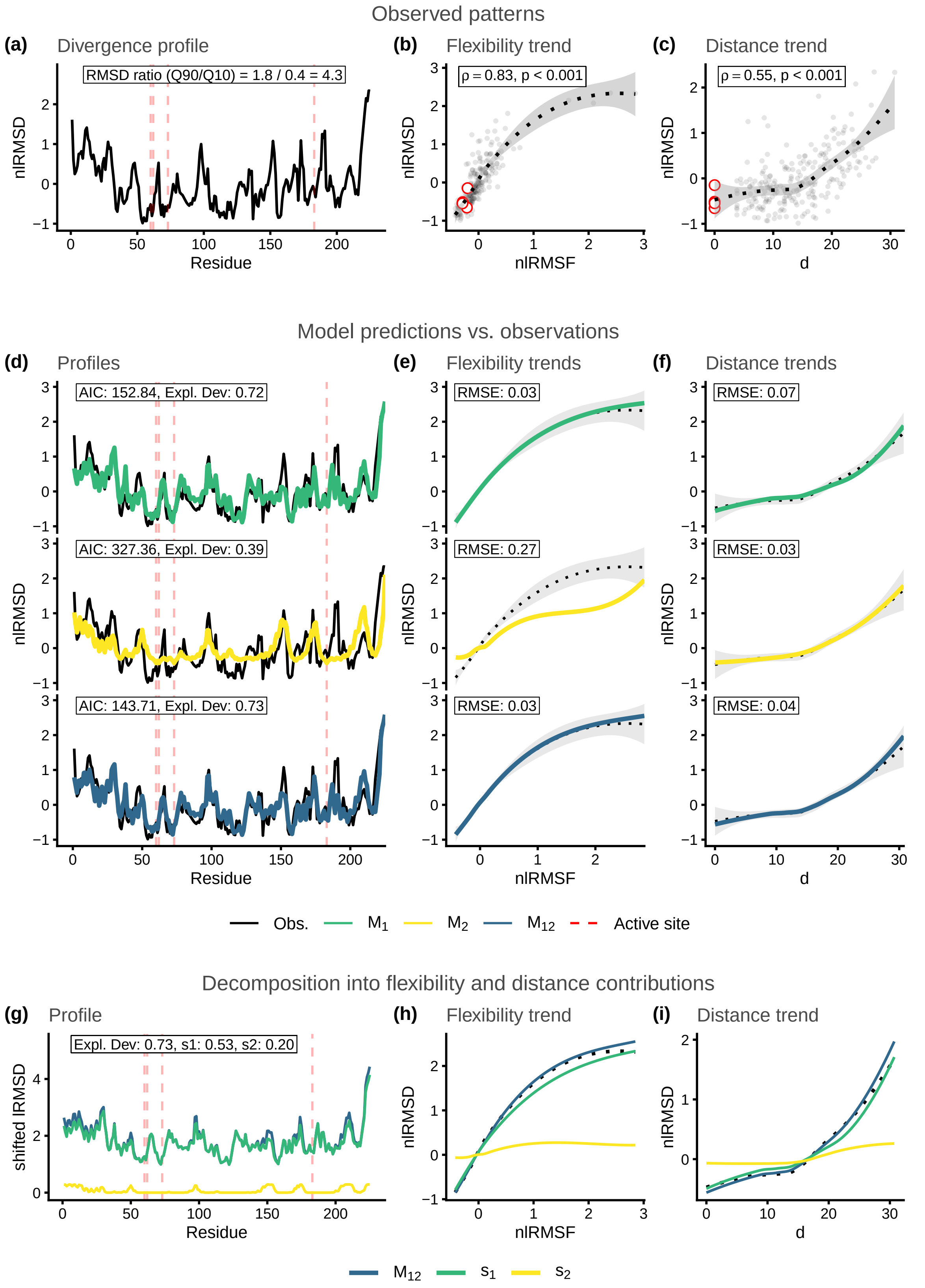

Figure S8: Structural divergence analysis for enzyme family MCSA ID: 71. Reference protein PDB ID: 1eug_A.

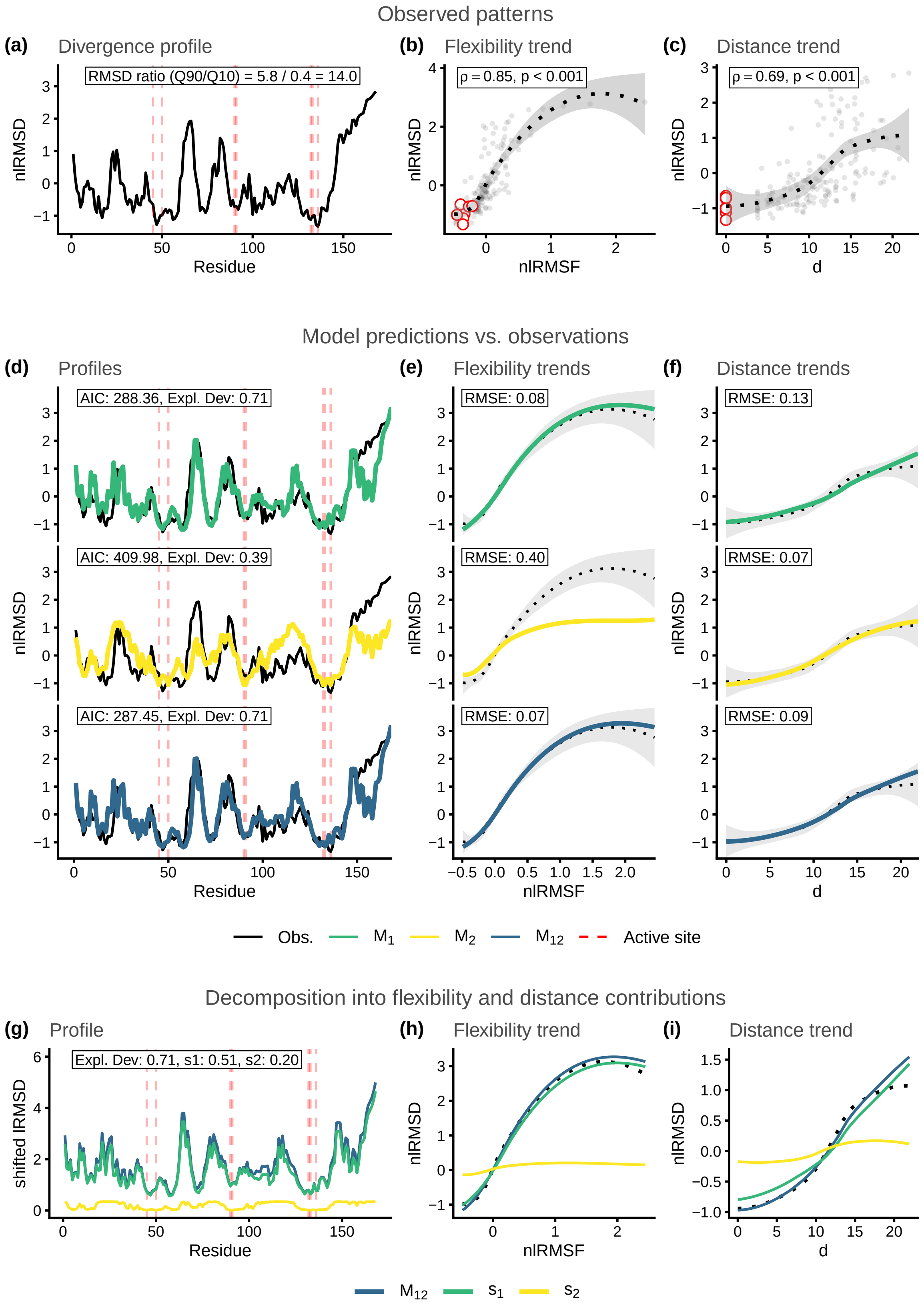

Figure S9: Structural divergence analysis for enzyme family MCSA ID: 98. Reference protein PDB ID: 1bsz_C.

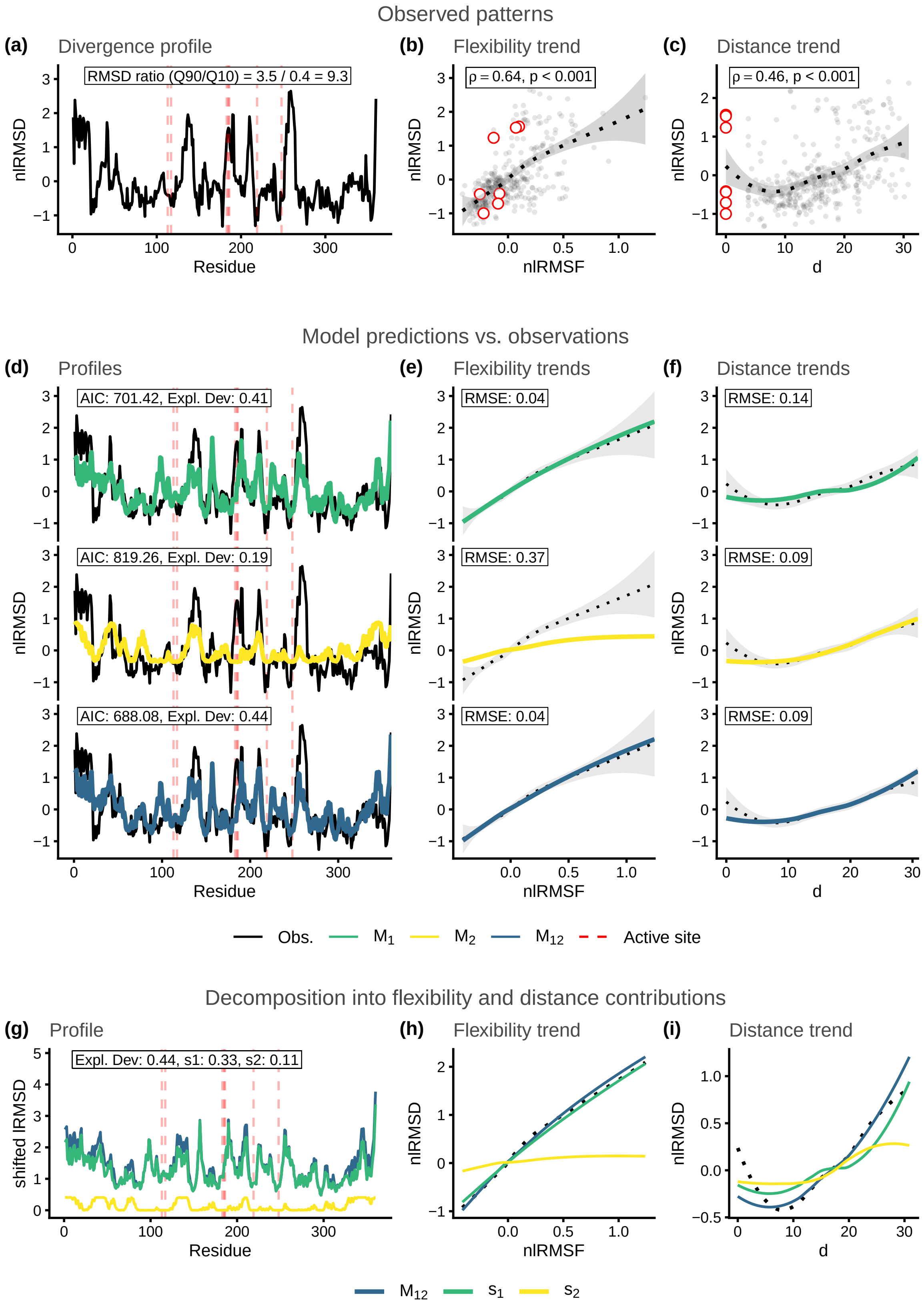

Figure S10: Structural divergence analysis for enzyme family MCSA ID: 109. Reference protein PDB ID: 1d3g_A.

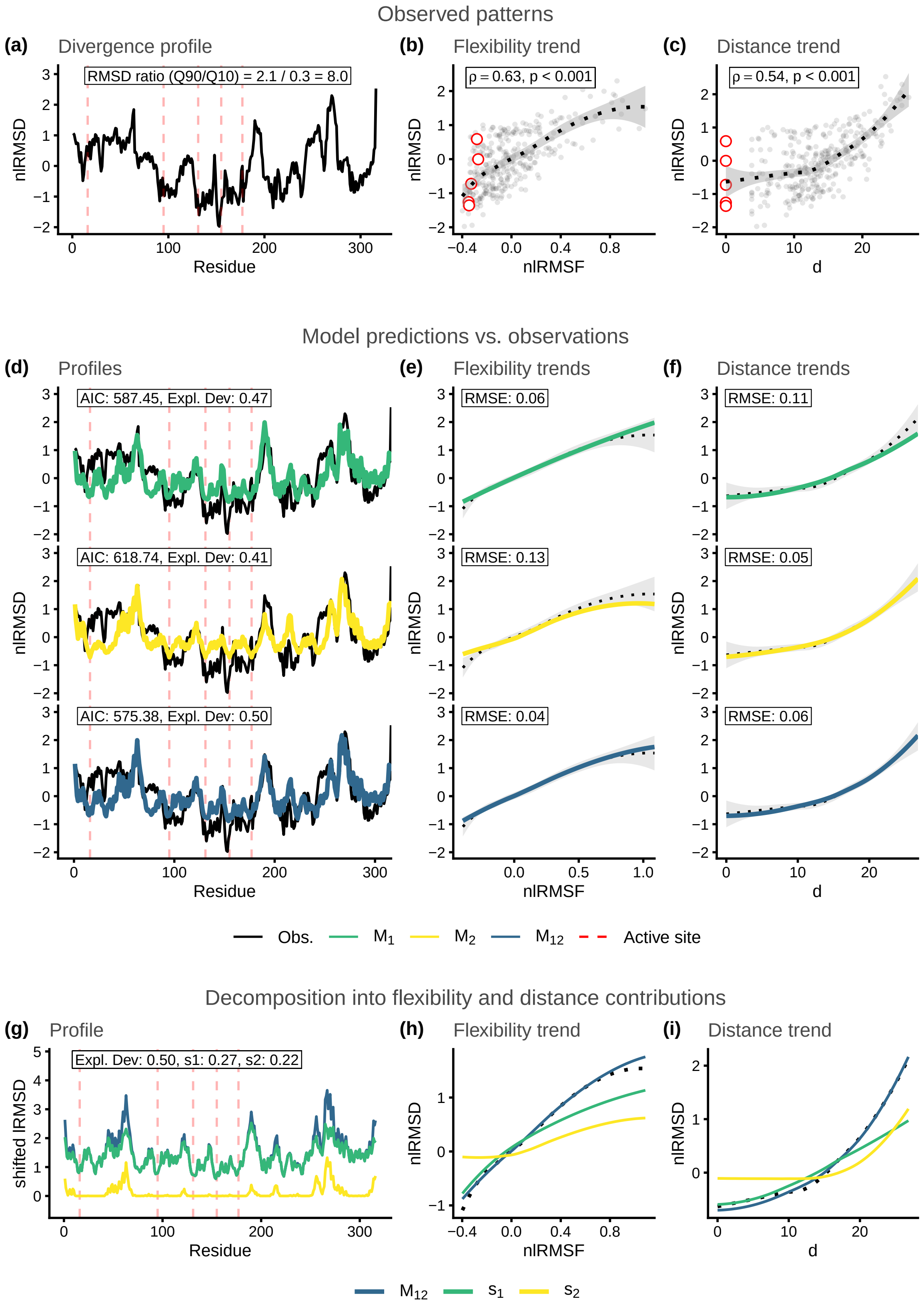

Figure S11: Structural divergence analysis for enzyme family MCSA ID: 148. Reference protein PDB ID: 1onr_A.

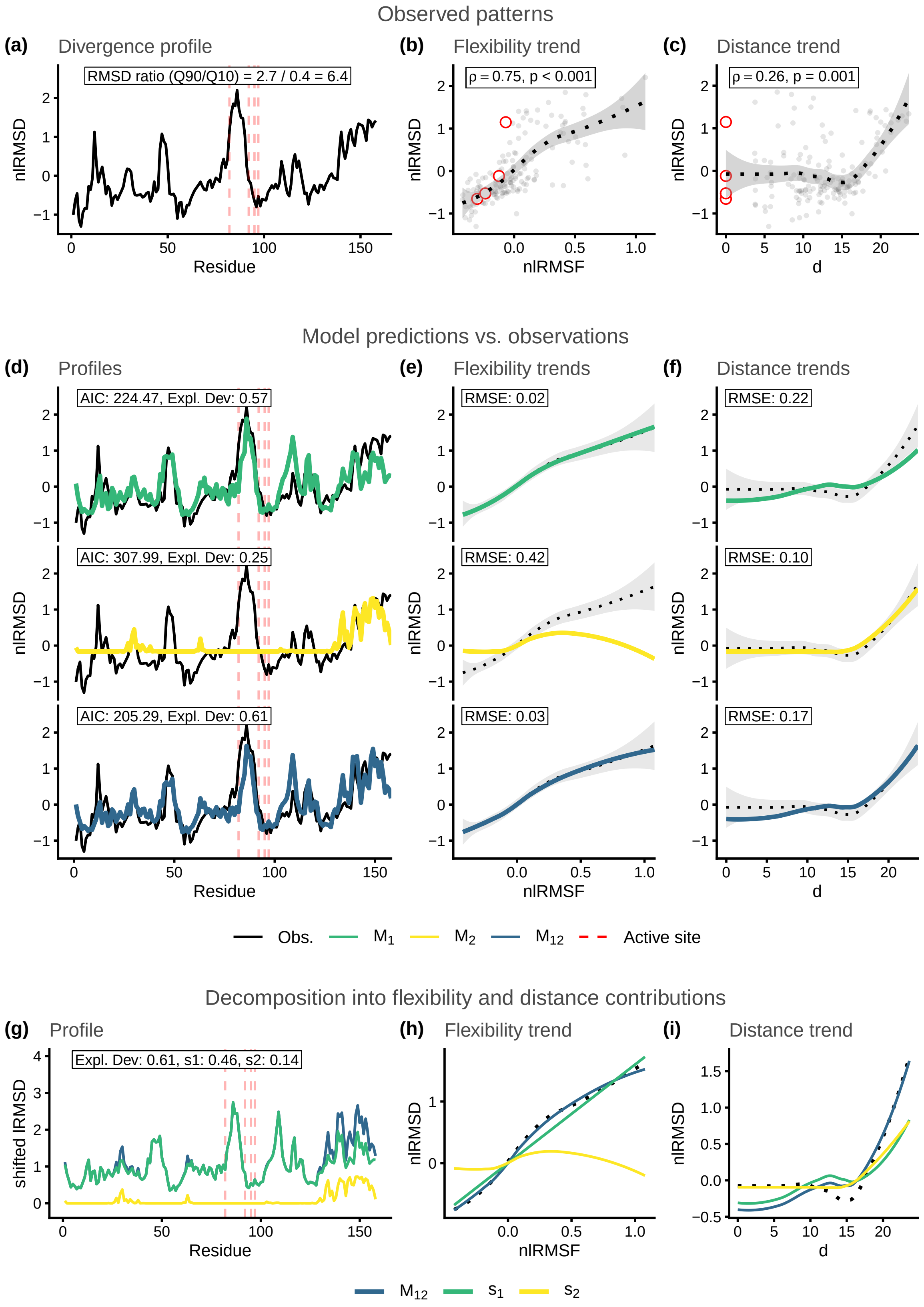

Figure S12: Structural divergence analysis for enzyme family MCSA ID: 151. Reference protein PDB ID: 1q0n_A.

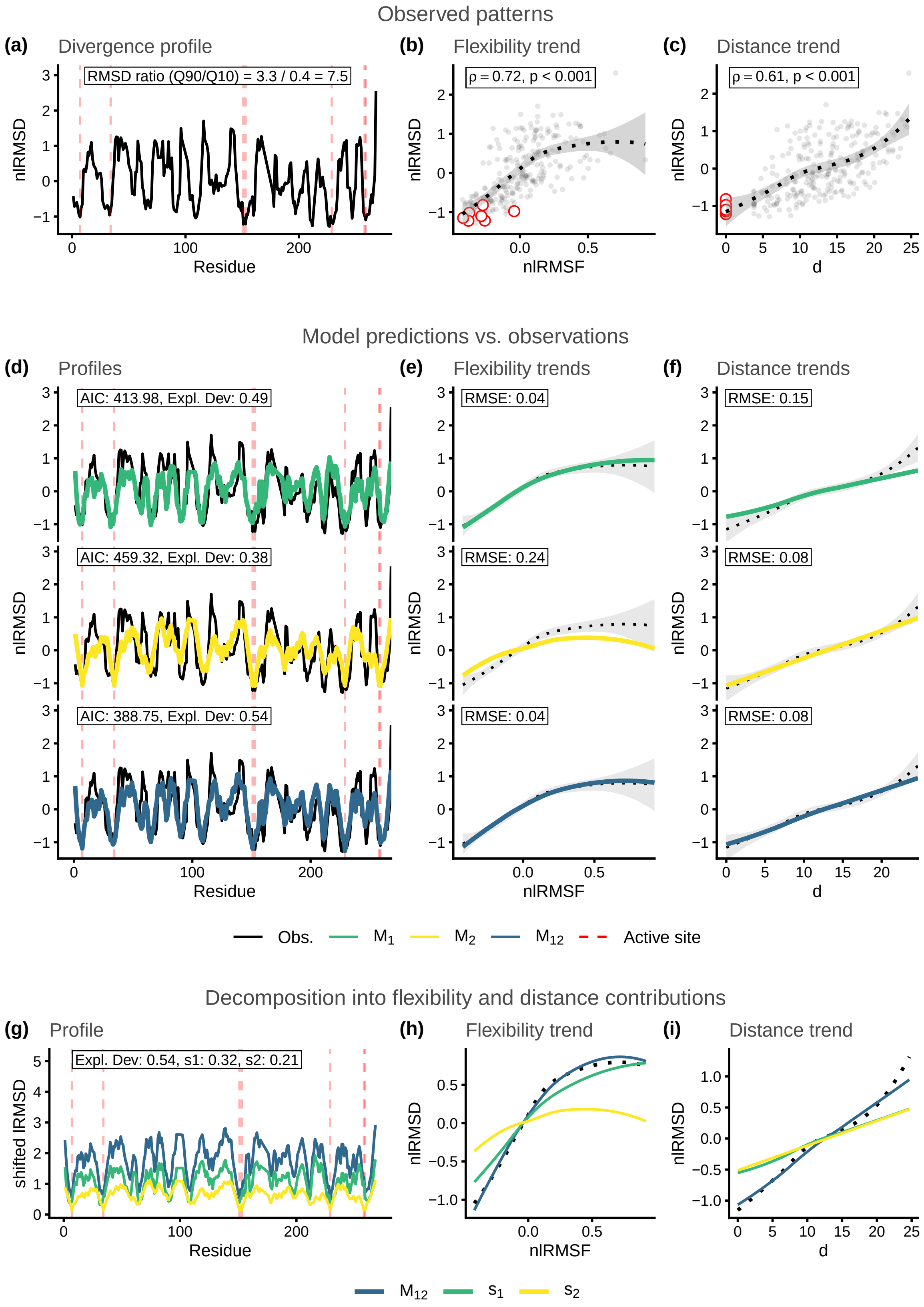

Figure S13: Structural divergence analysis for enzyme family MCSA ID: 160. Reference protein PDB ID: 1ako_A.

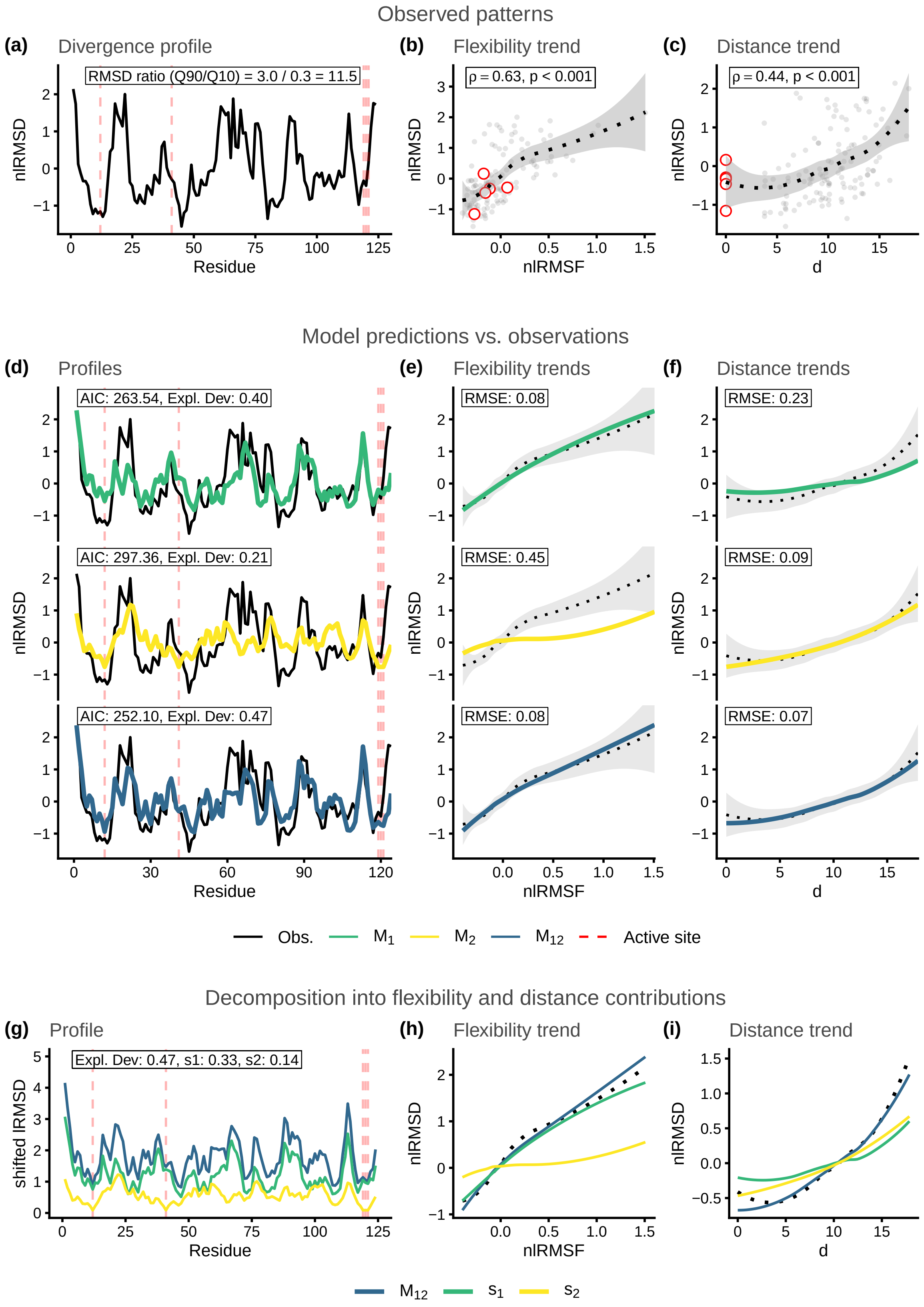

Figure S14: Structural divergence analysis for enzyme family MCSA ID: 164. Reference protein PDB ID: 1ruv_A.

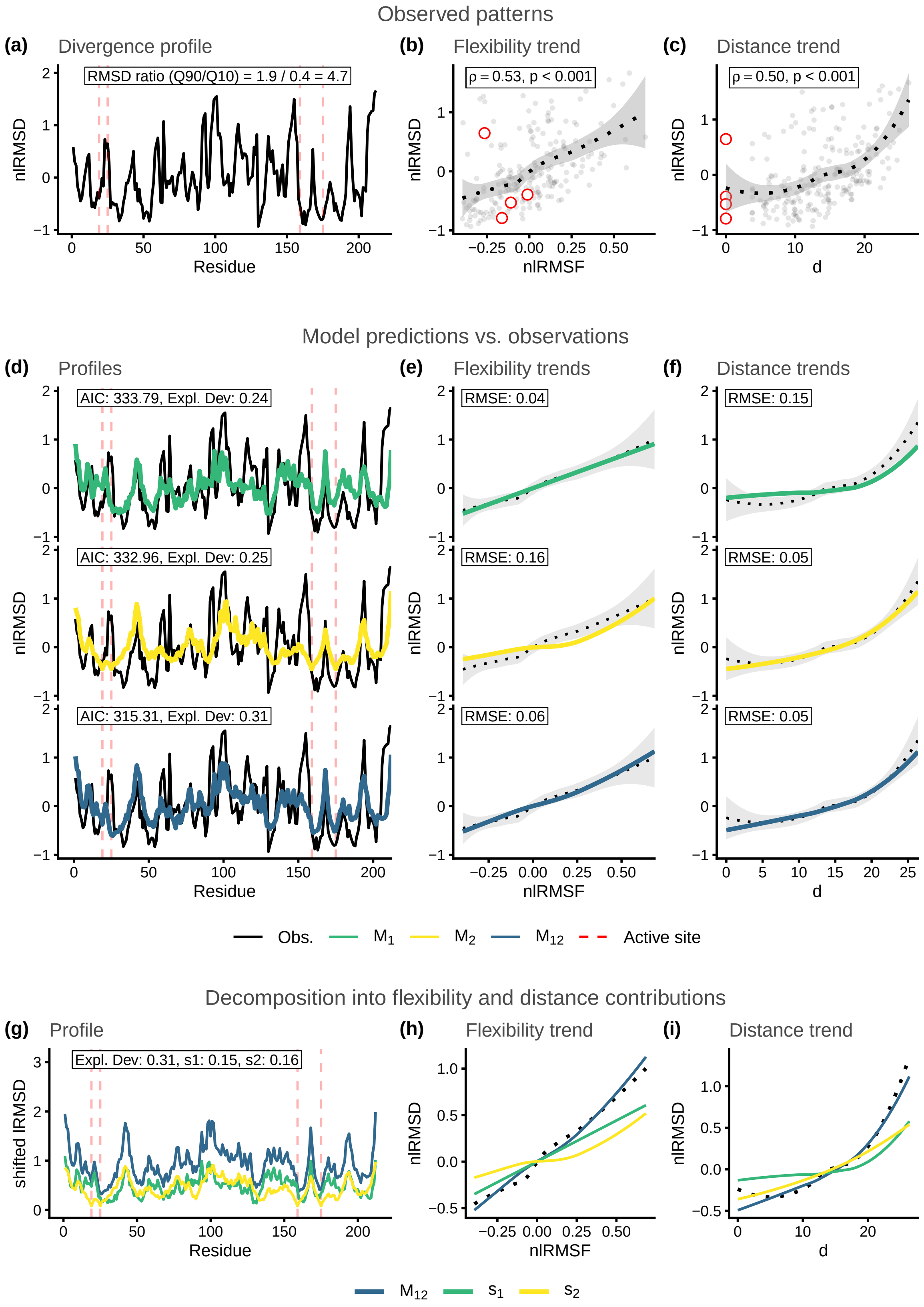

Figure S15: Structural divergence analysis for enzyme family MCSA ID: 174. Reference protein PDB ID: 9pap_A.

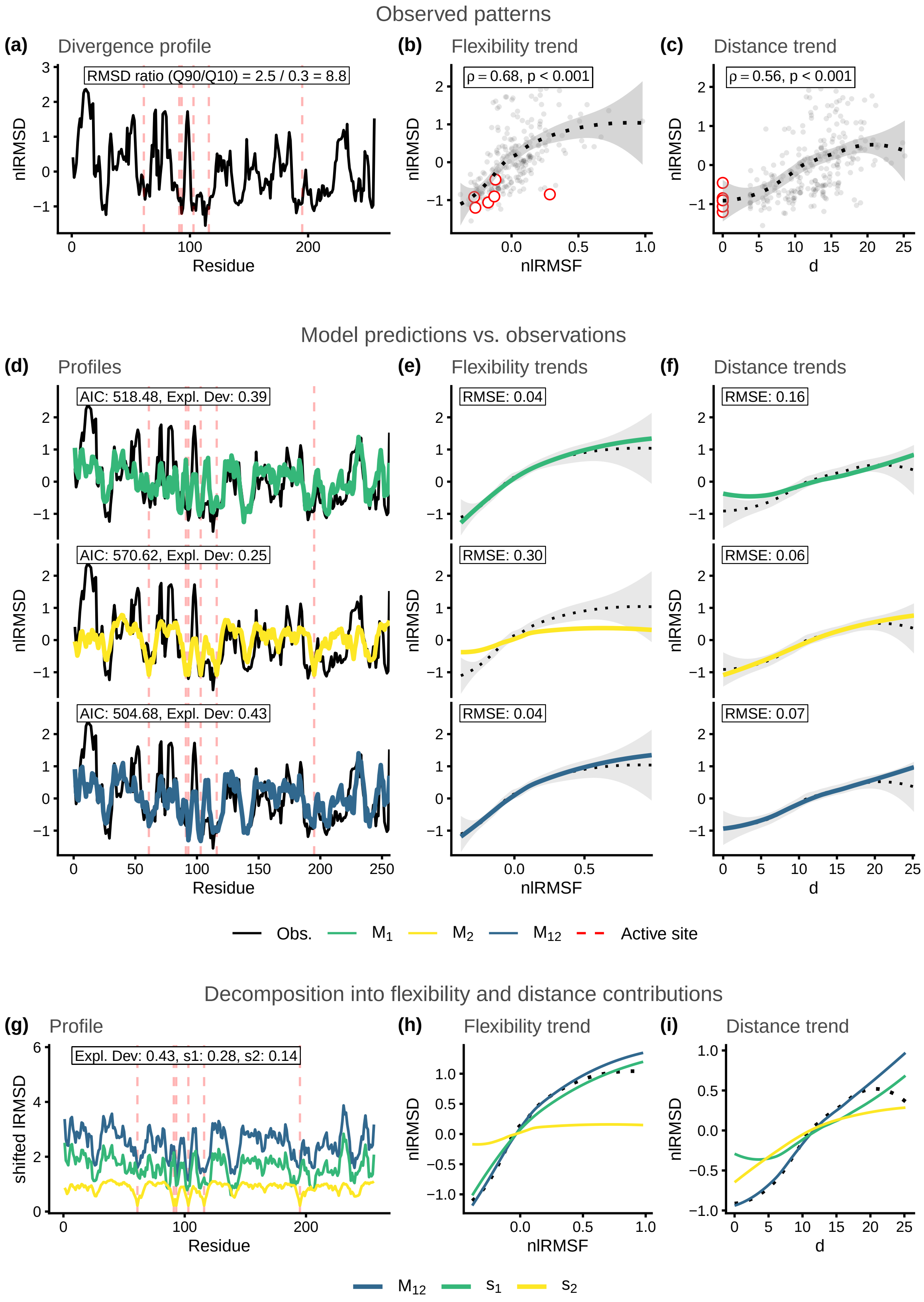

Figure S16: Structural divergence analysis for enzyme family MCSA ID: 216. Reference protein PDB ID: 1ca2_A.

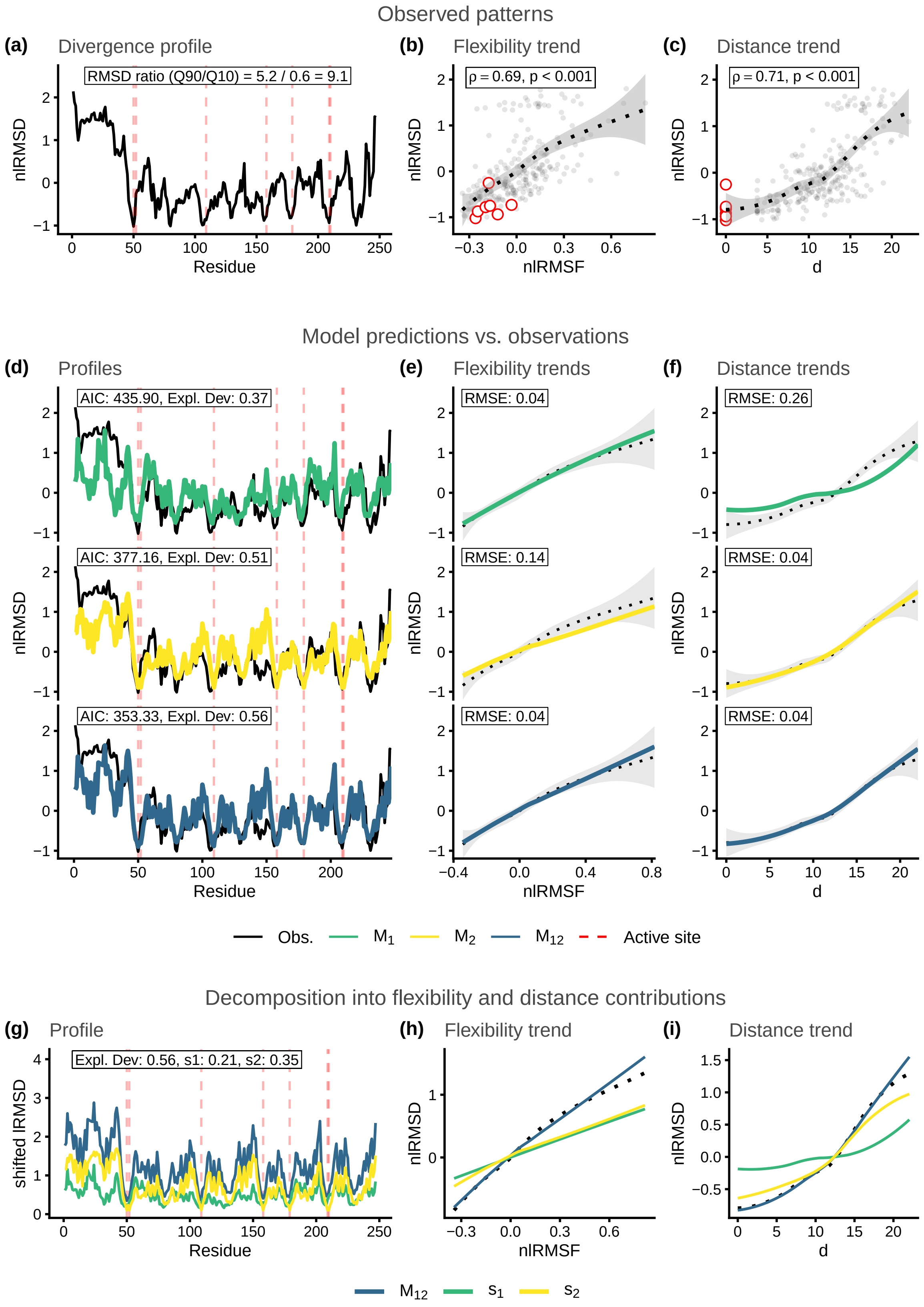

Figure S17: Structural divergence analysis for enzyme family MCSA ID: 252. Reference protein PDB ID: 1igs_A.

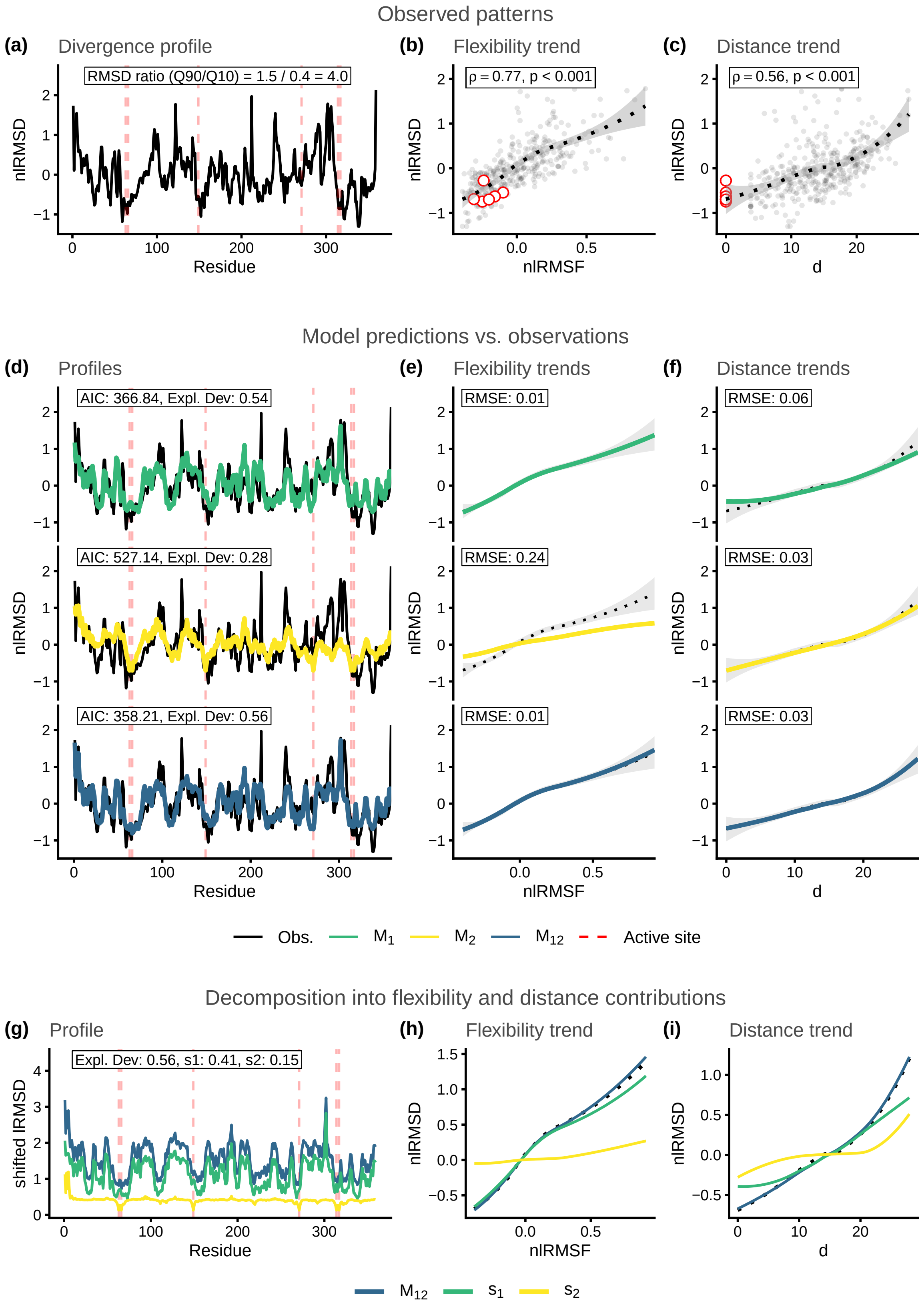

Figure S18: Structural divergence analysis for enzyme family MCSA ID: 257. Reference protein PDB ID: 1xx2_A.

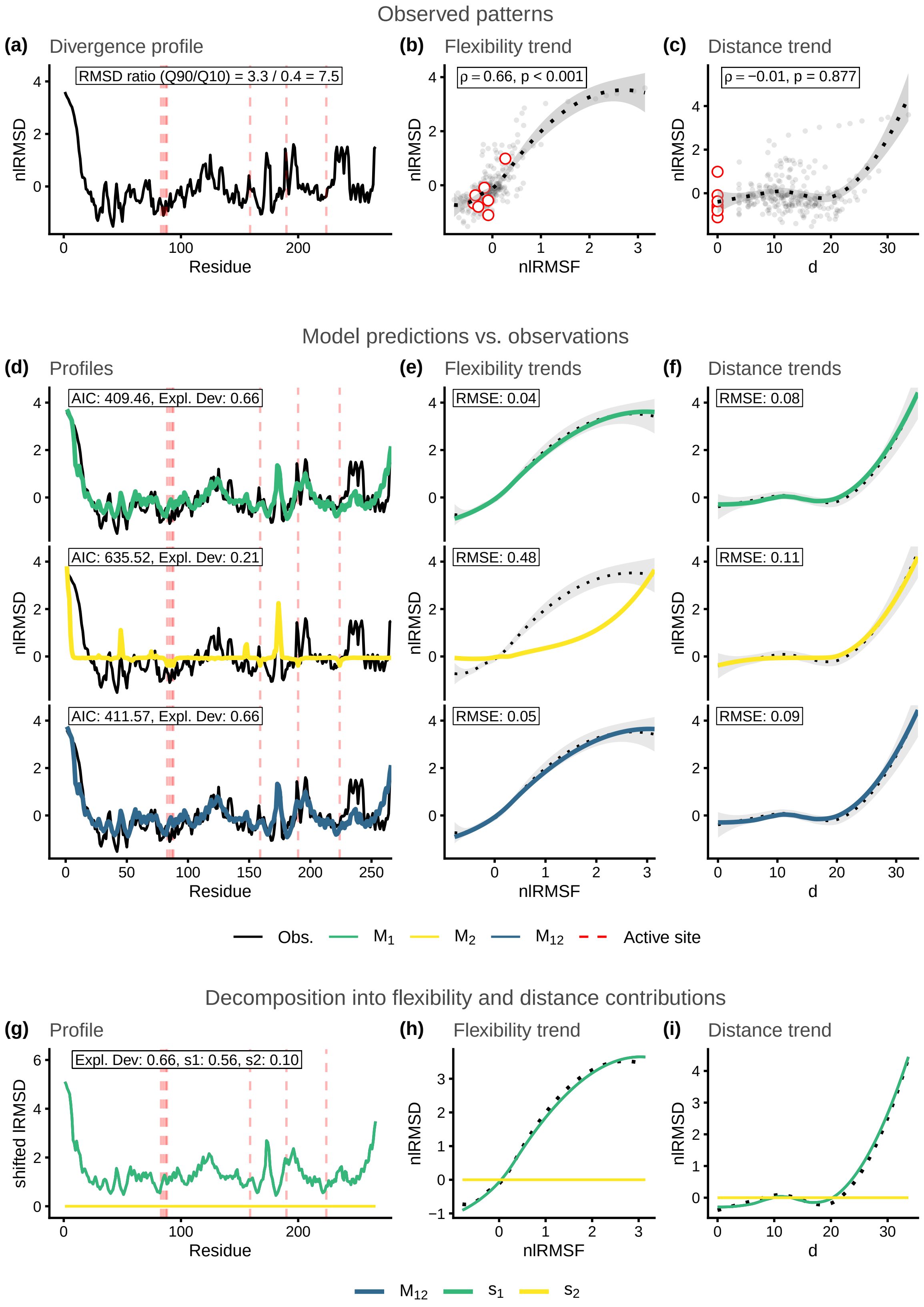

Figure S19: Structural divergence analysis for enzyme family MCSA ID: 258. Reference protein PDB ID: 1sml_A.

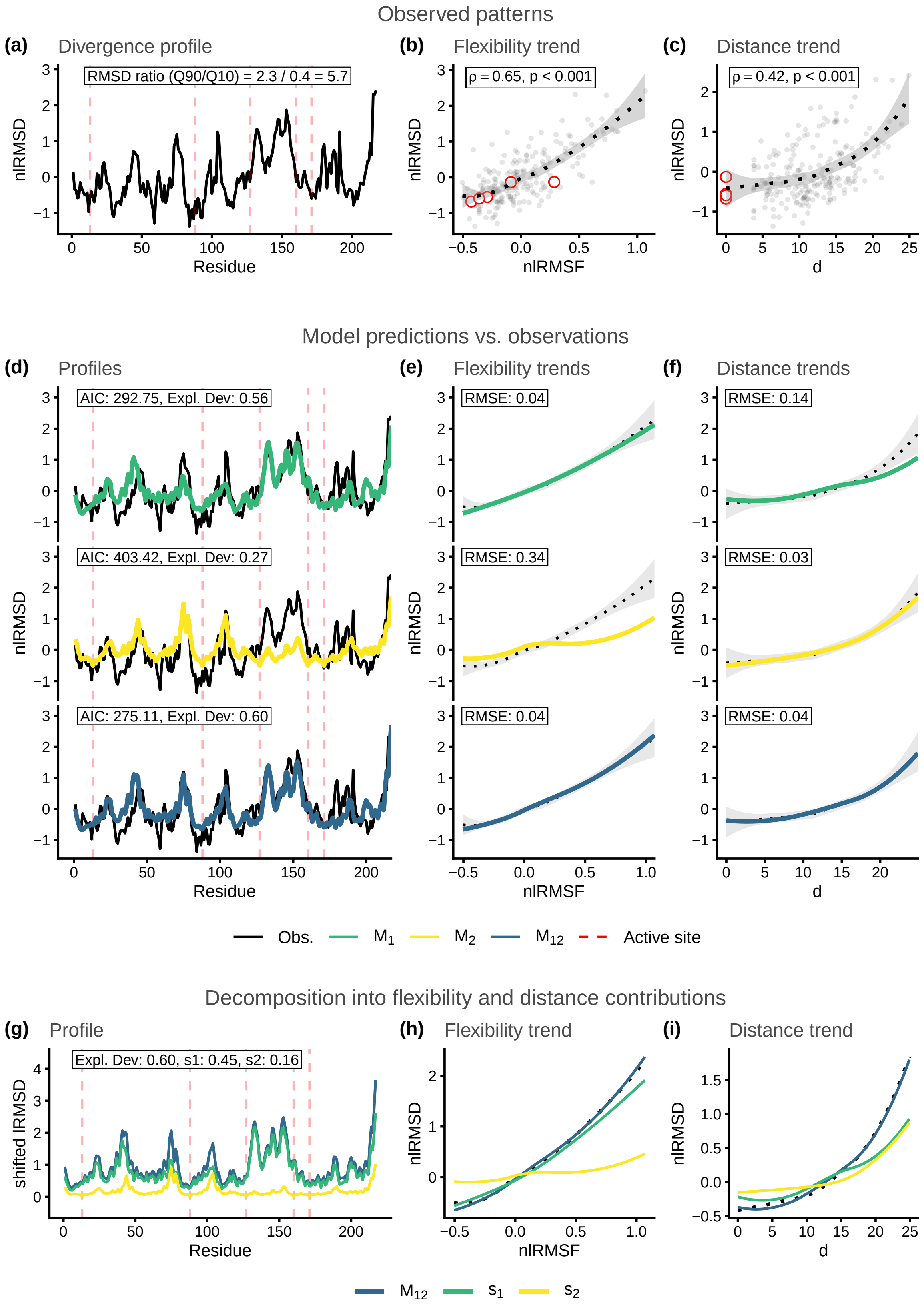

Figure S20: Structural divergence analysis for enzyme family MCSA ID: 290. Reference protein PDB ID: 1zio_A.

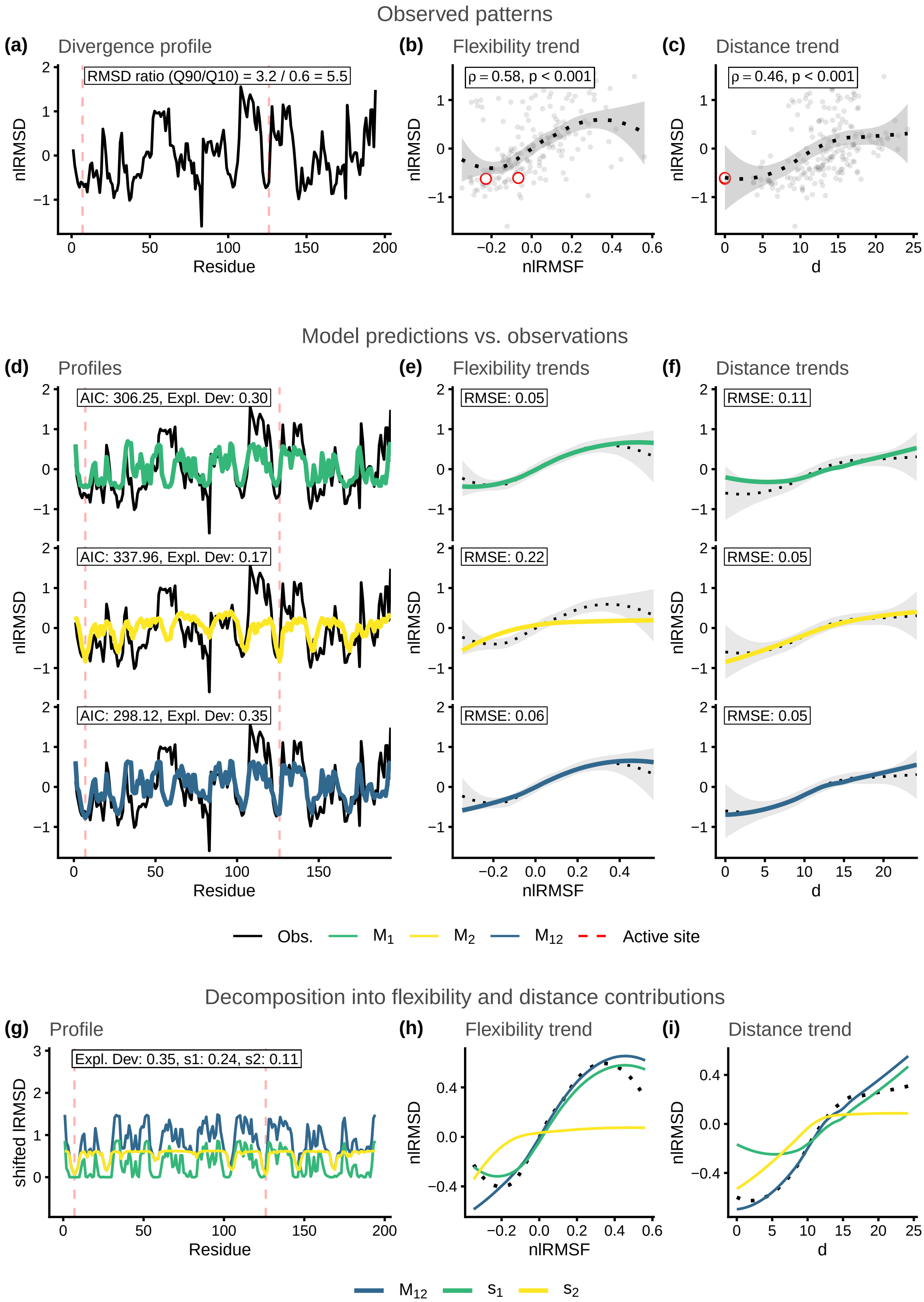

Figure S21: Structural divergence analysis for enzyme family MCSA ID: 328. Reference protein PDB ID: 1lbm_A.

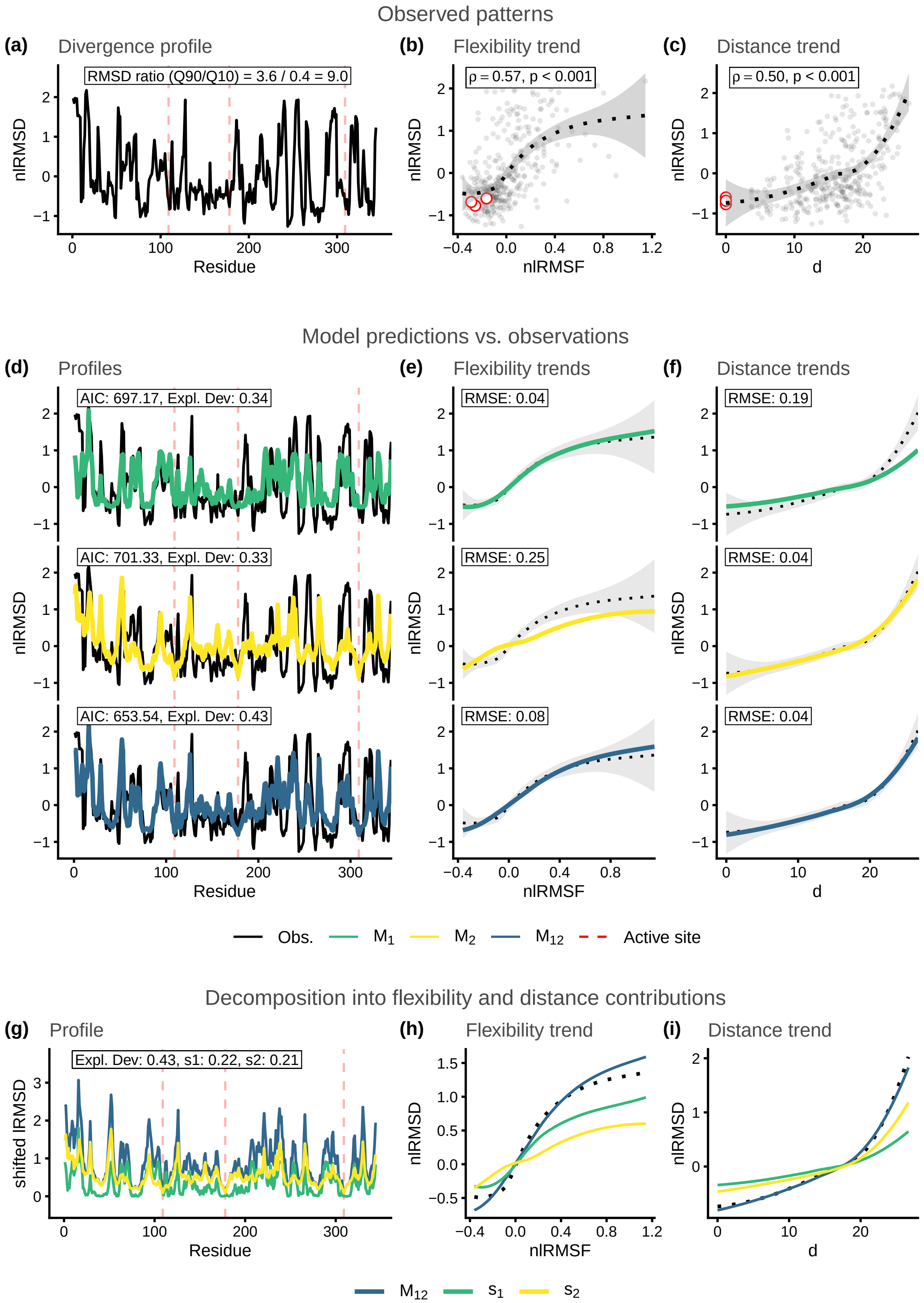

Figure S22: Structural divergence analysis for enzyme family MCSA ID: 351. Reference protein PDB ID: 1snz_A.

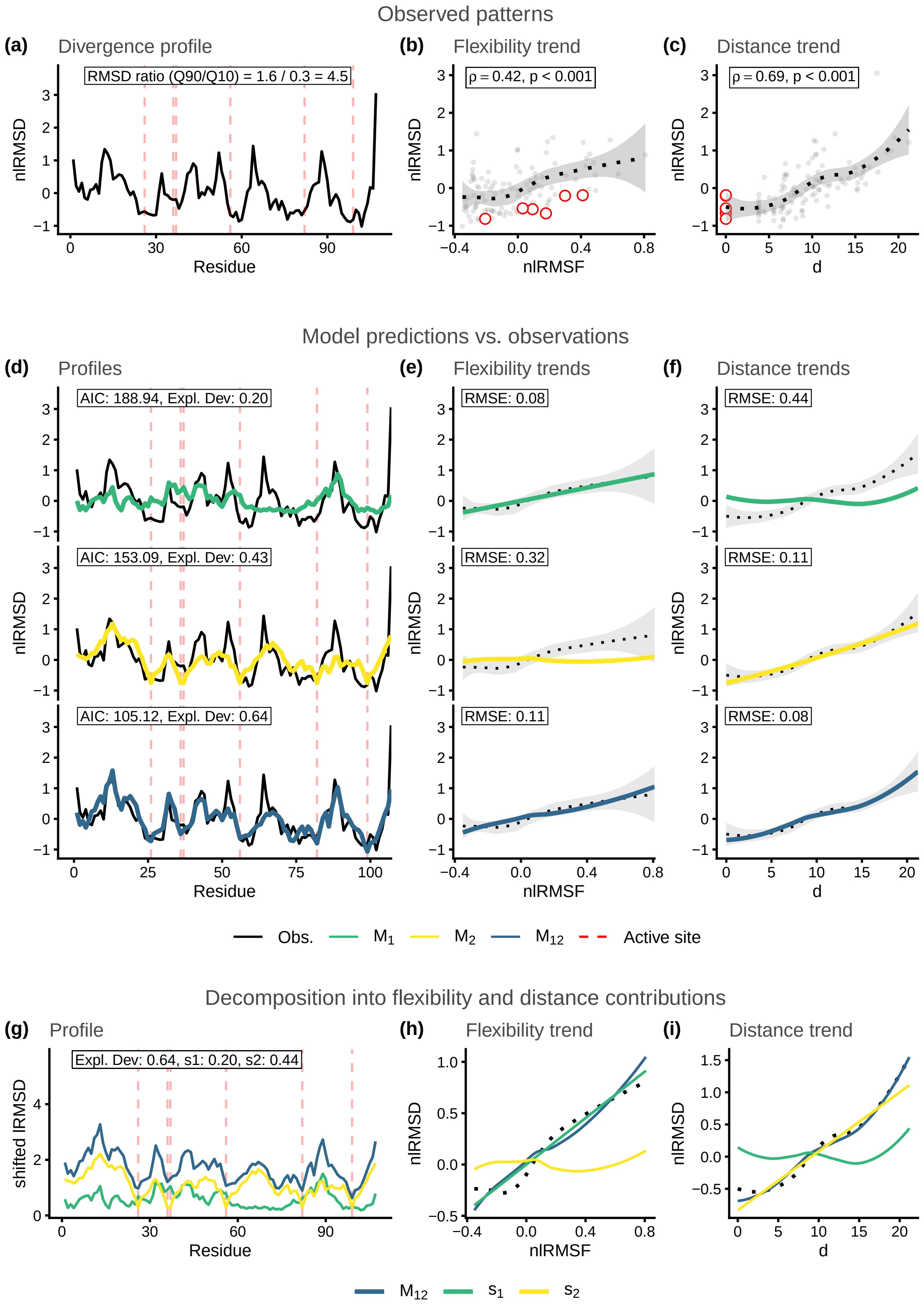

Figure S23: Structural divergence analysis for enzyme family MCSA ID: 362. Reference protein PDB ID: 1d6o_A.

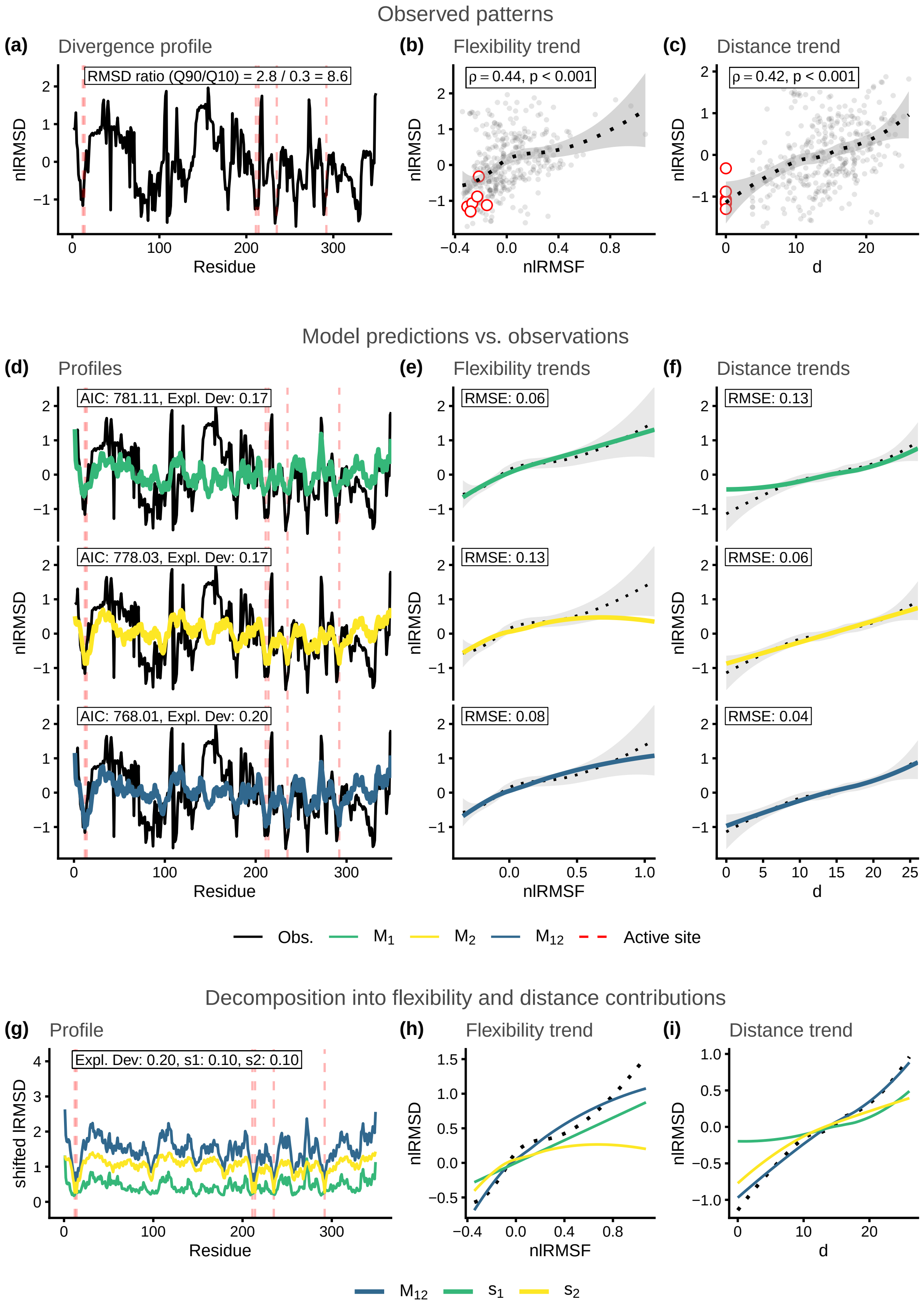

Figure S24: Structural divergence analysis for enzyme family MCSA ID: 376. Reference protein PDB ID: 1a4l_A.

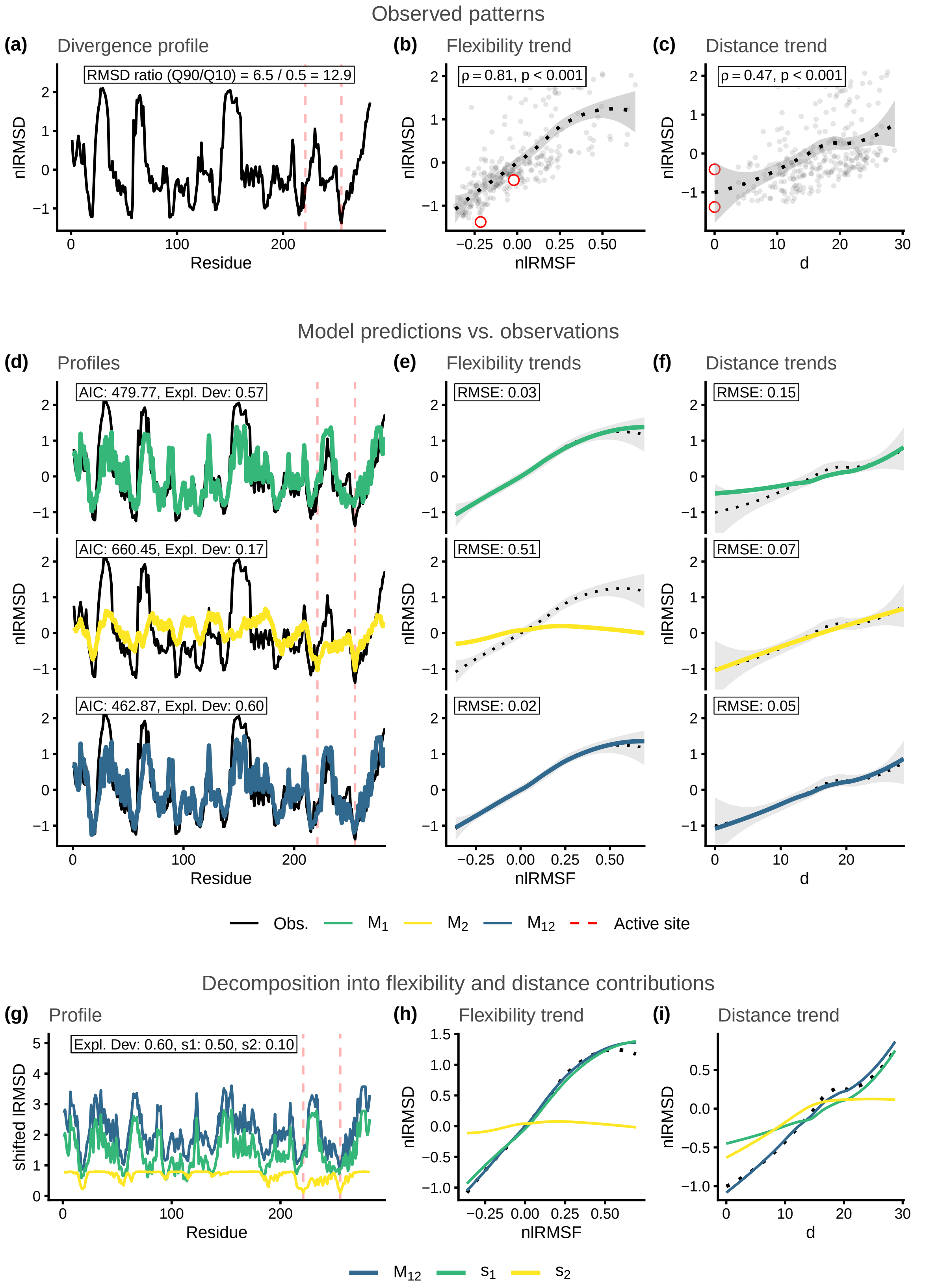

Figure S25: Structural divergence analysis for enzyme family MCSA ID: 394. Reference protein PDB ID: 1aj0_A.

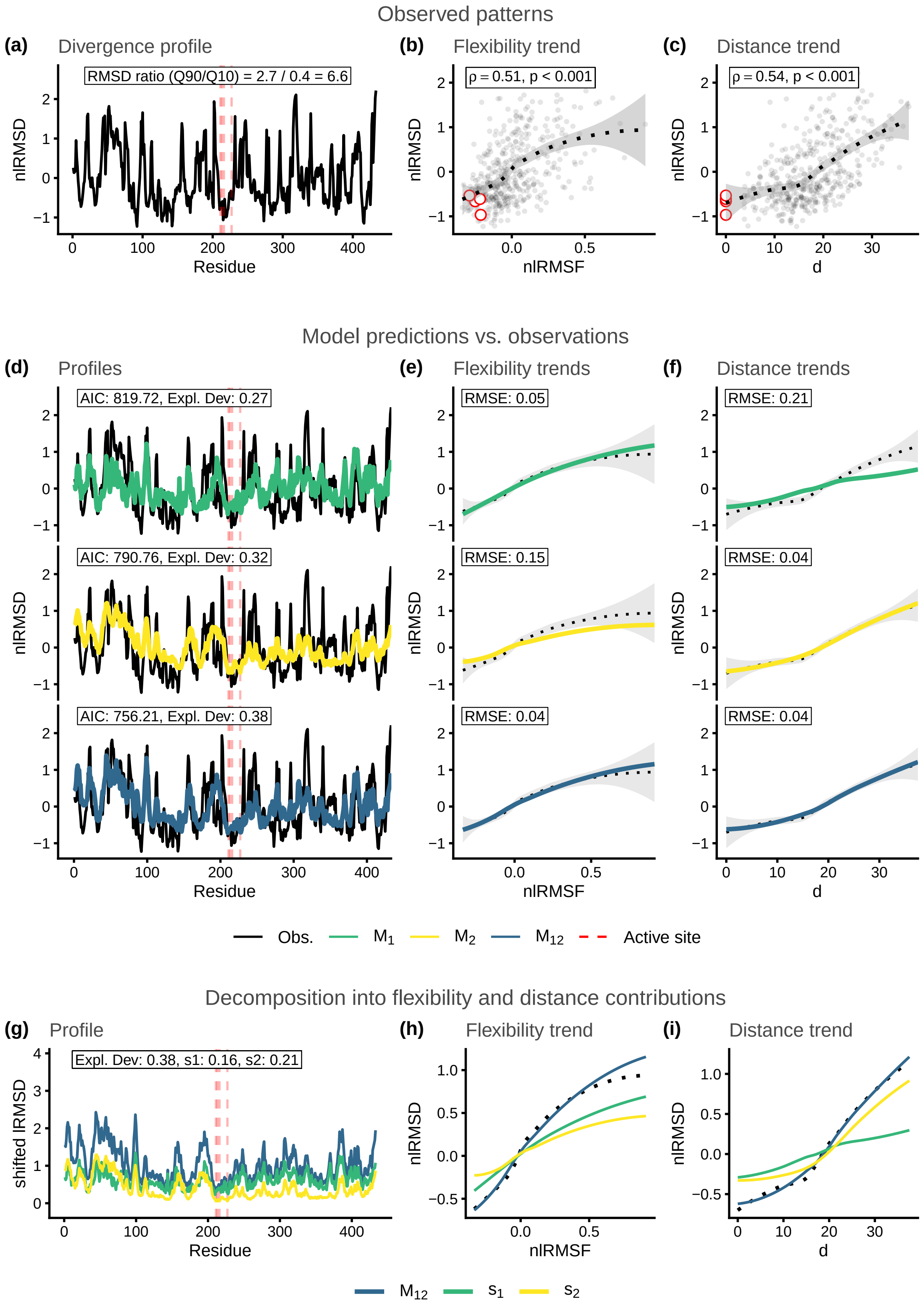

Figure S26: Structural divergence analysis for enzyme family MCSA ID: 444. Reference protein PDB ID: 1cel_A.

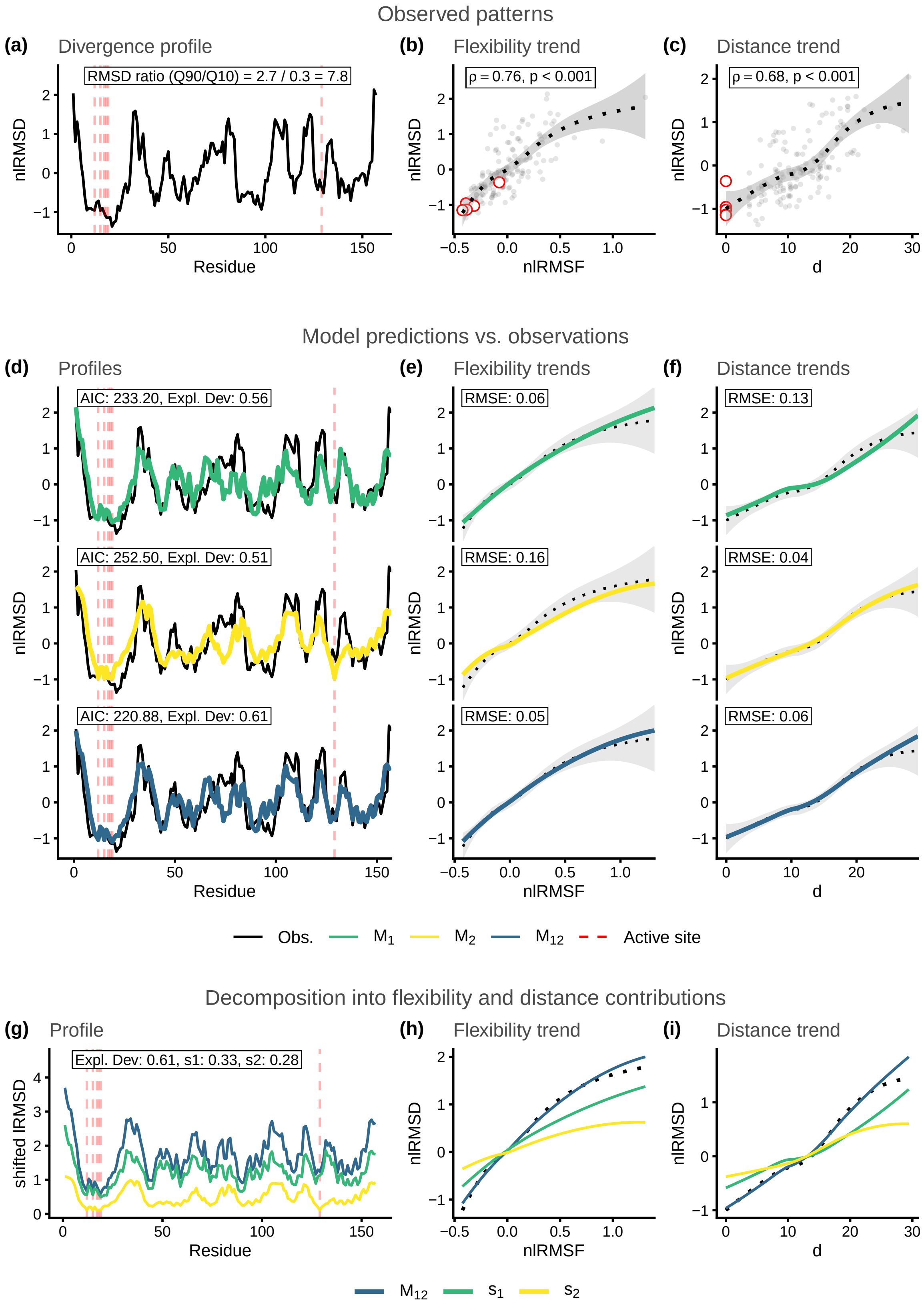

Figure S27: Structural divergence analysis for enzyme family MCSA ID: 462. Reference protein PDB ID: 1pnt_A.

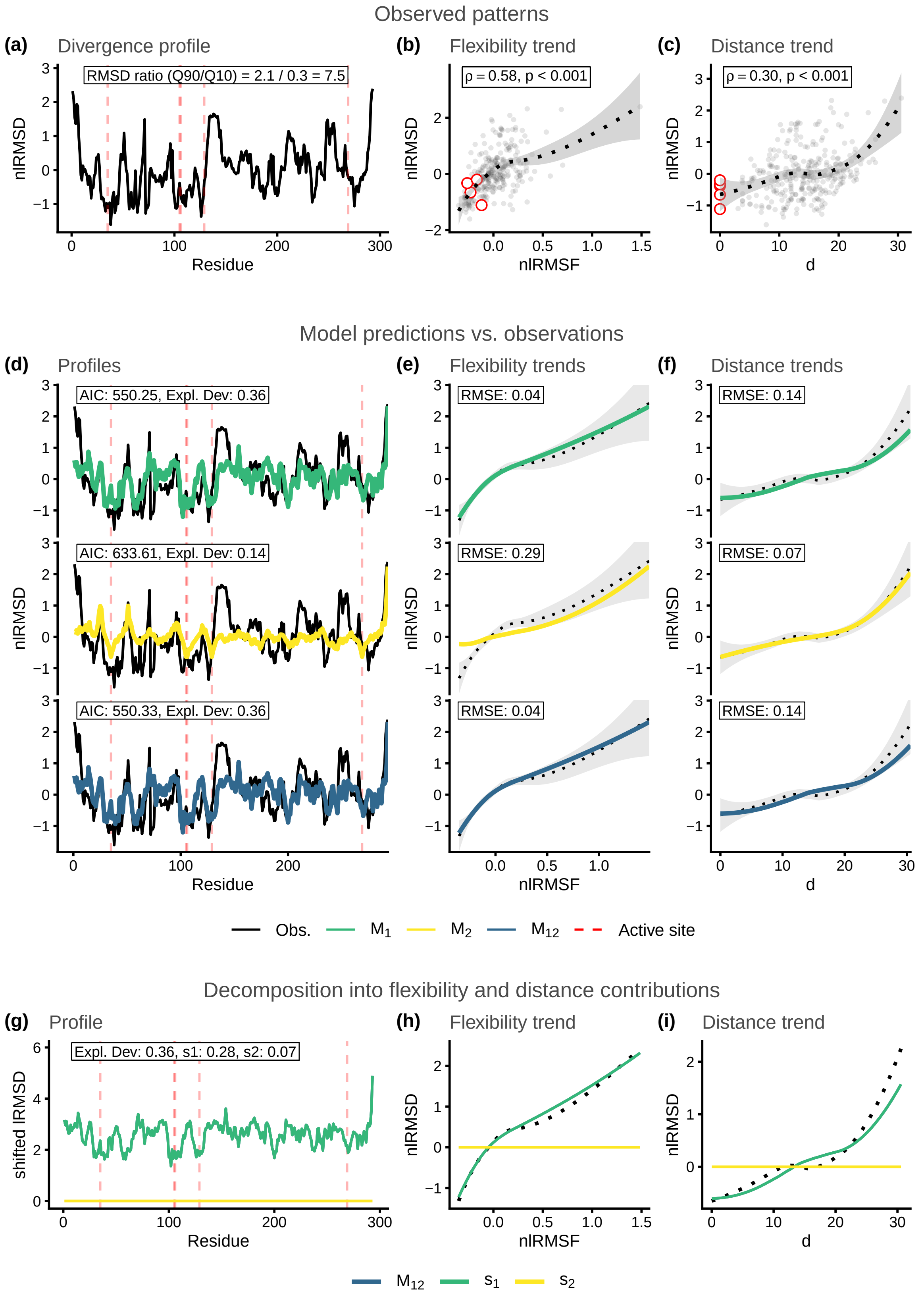

Figure S28: Structural divergence analysis for enzyme family MCSA ID: 467. Reference protein PDB ID: 1cv2_A.

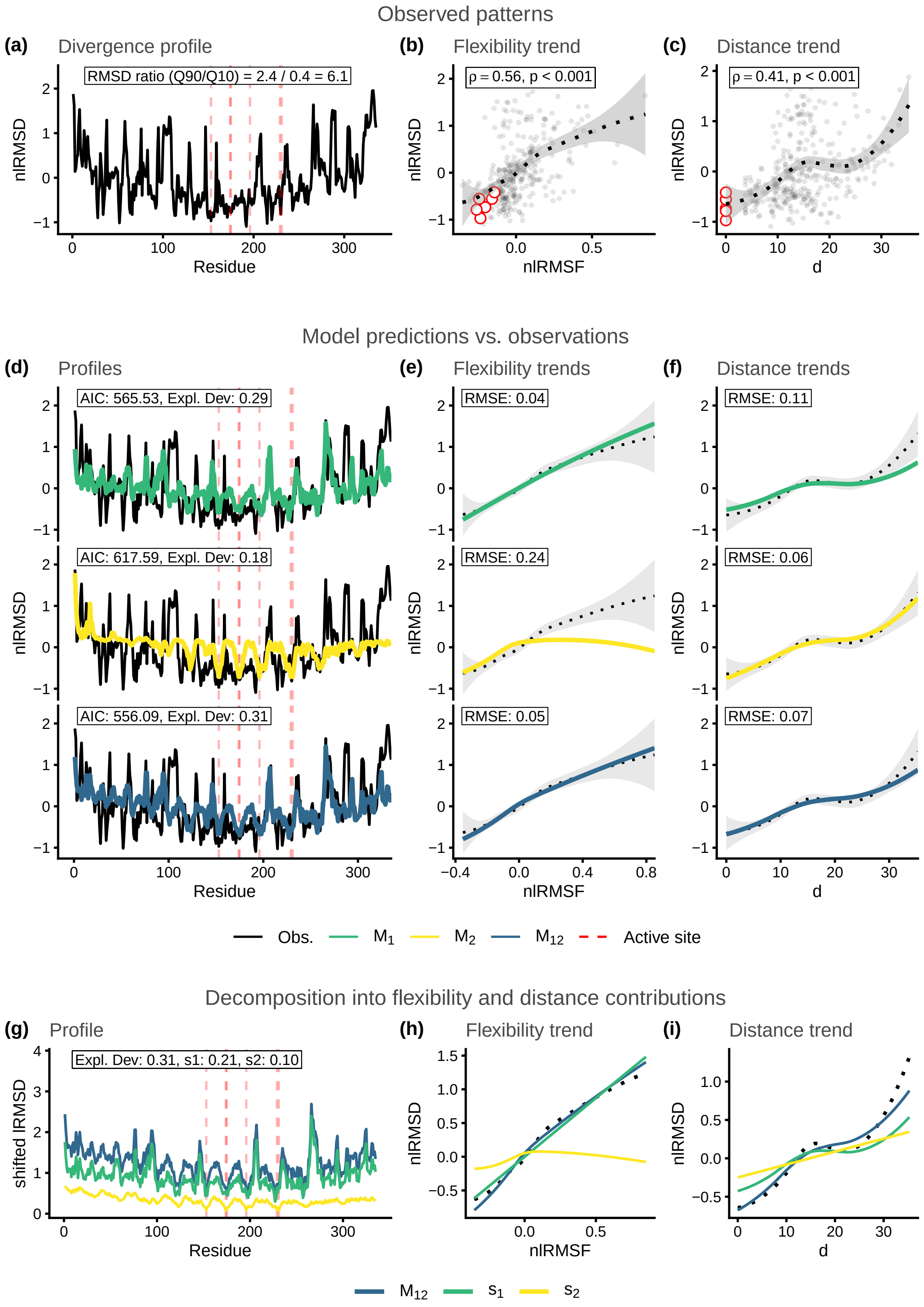

Figure S29: Structural divergence analysis for enzyme family MCSA ID: 480. Reference protein PDB ID: 1czf_A.

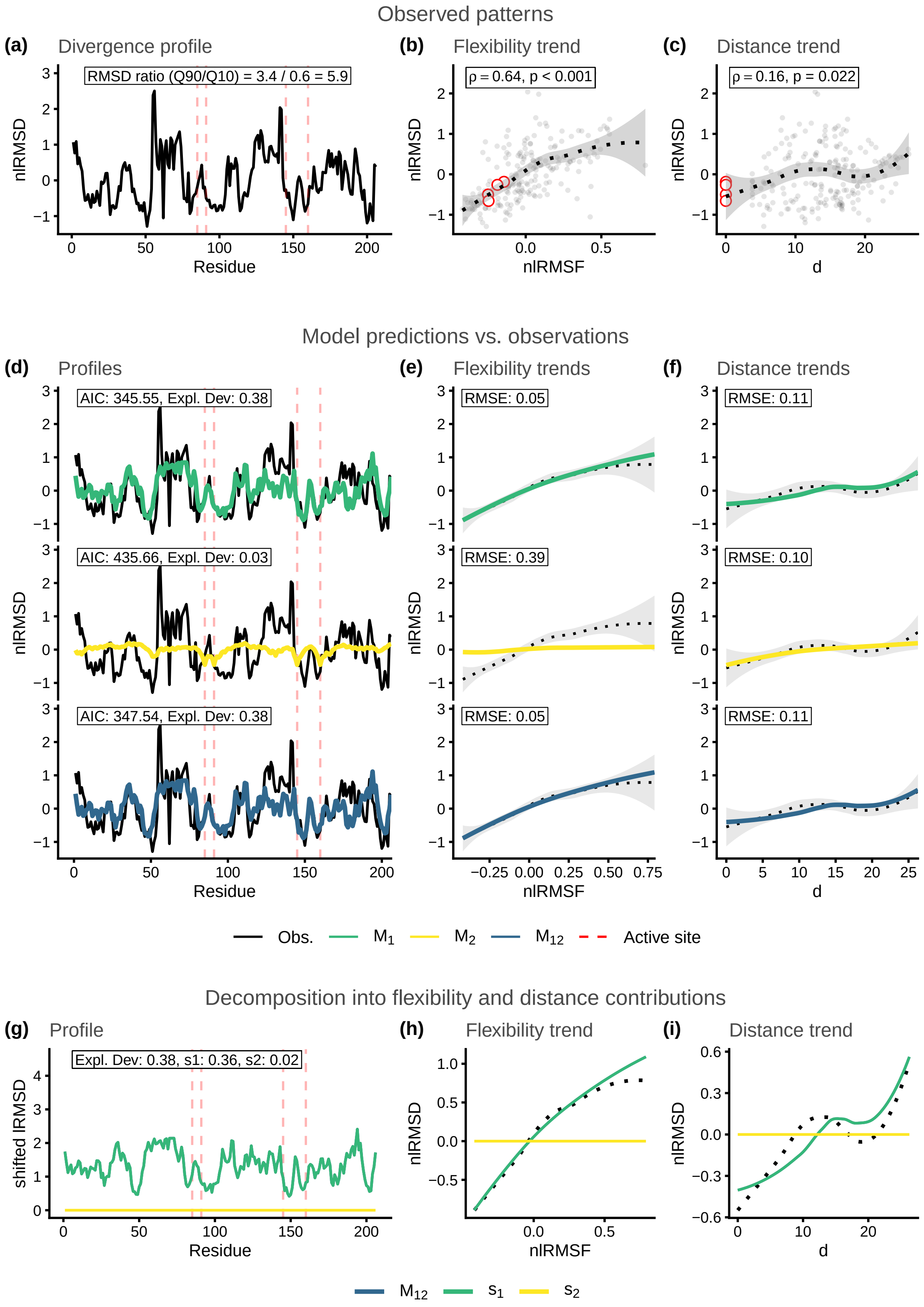

Figure S30: Structural divergence analysis for enzyme family MCSA ID: 597. Reference protein PDB ID: 1uch_A.

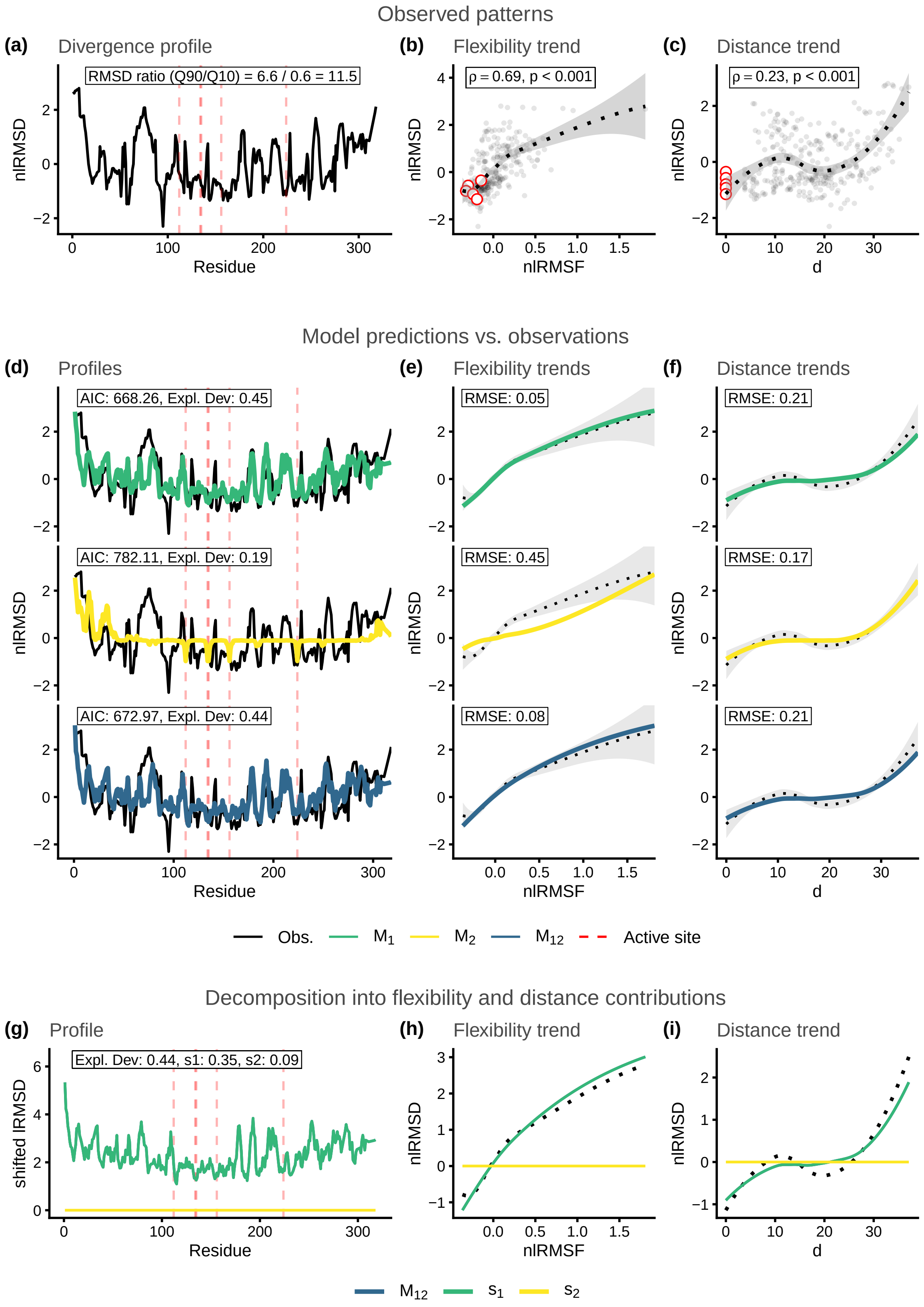

Figure S31: Structural divergence analysis for enzyme family MCSA ID: 681. Reference protein PDB ID: 1gq8_A.

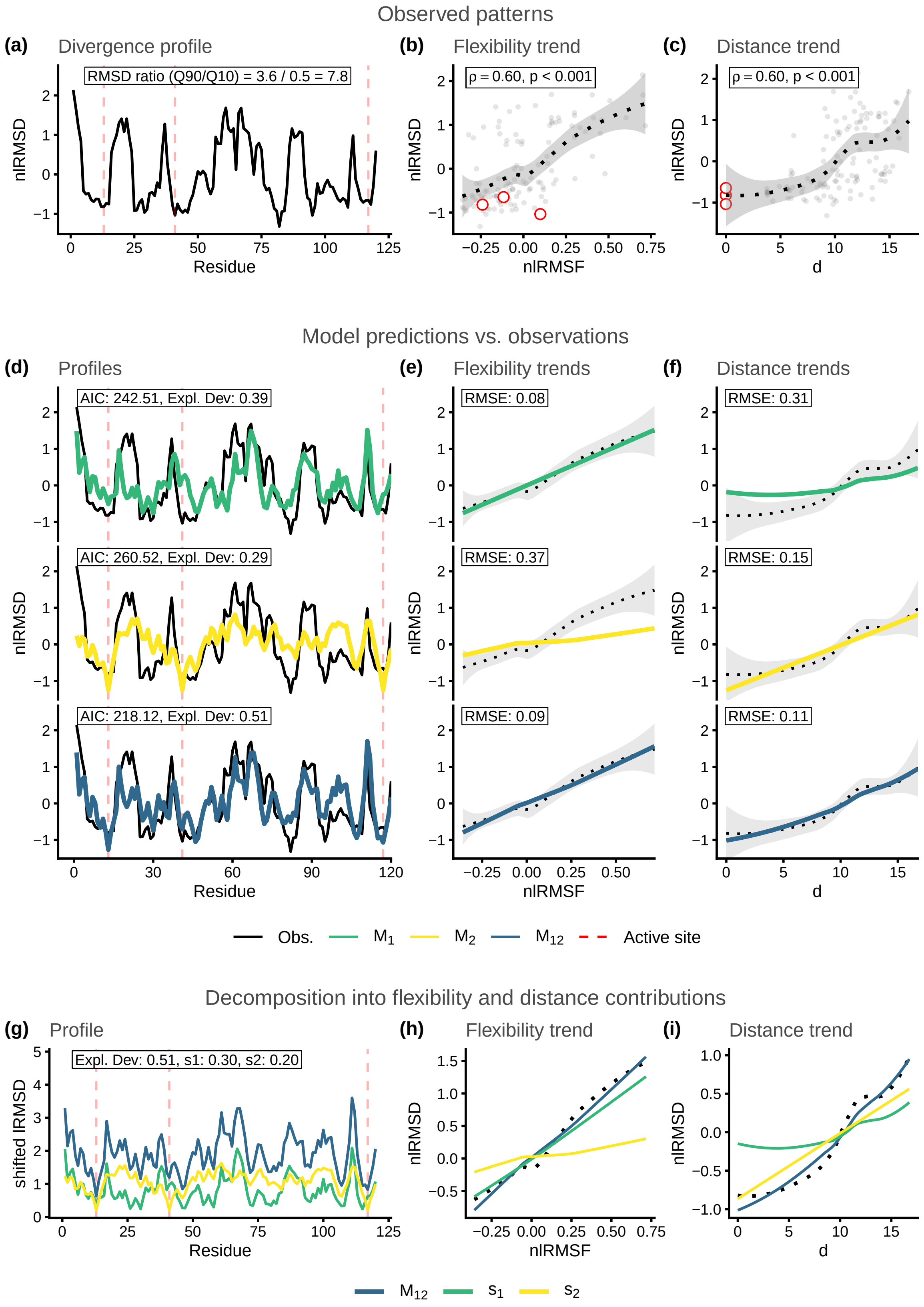

Figure S32: Structural divergence analysis for enzyme family MCSA ID: 693. Reference protein PDB ID: 2rnf_A.

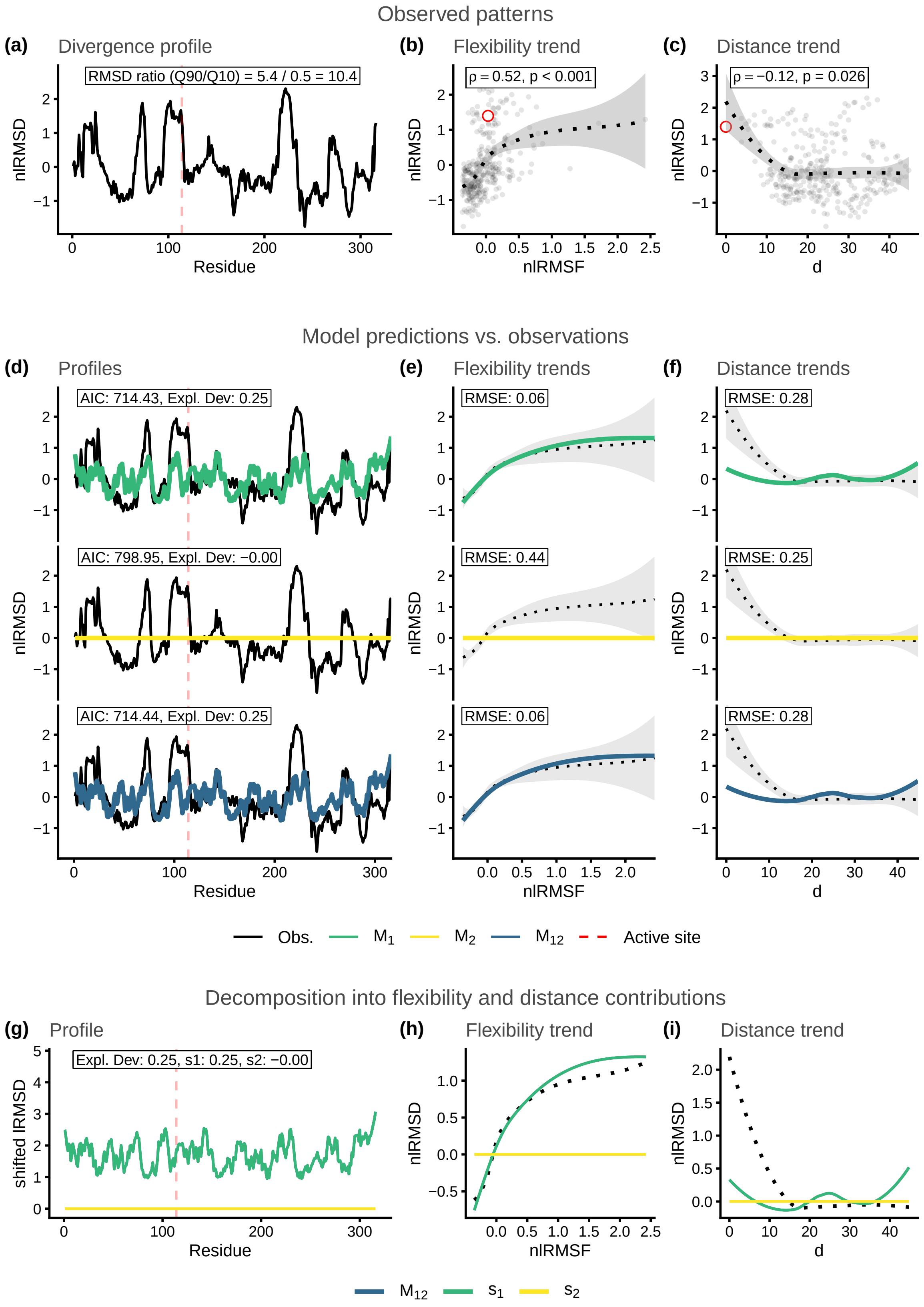

Figure S33: Structural divergence analysis for enzyme family MCSA ID: 749. Reference protein PDB ID: 1nml_A.

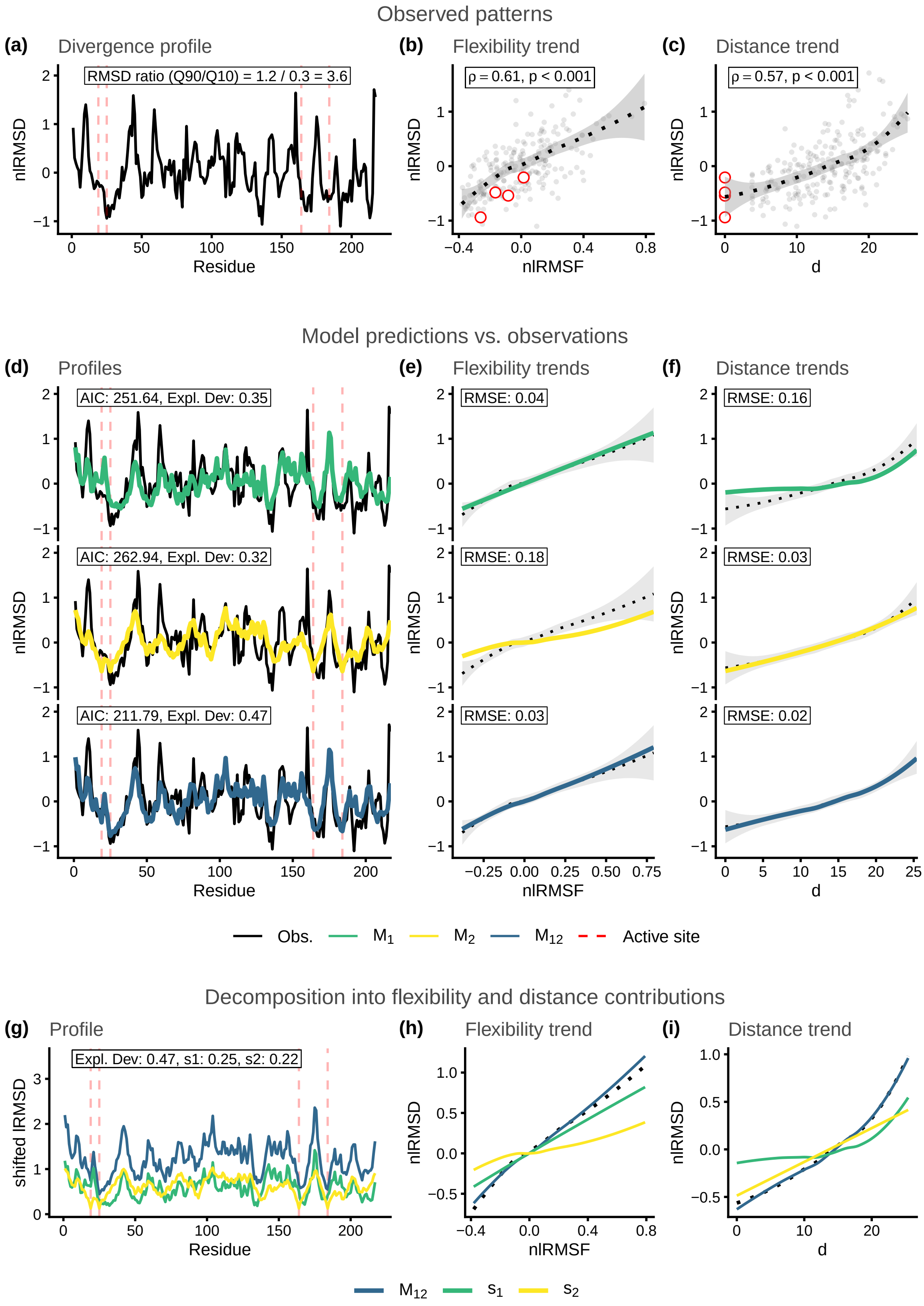

Figure S34: Structural divergence analysis for enzyme family MCSA ID: 814. Reference protein PDB ID: 1glo_A.

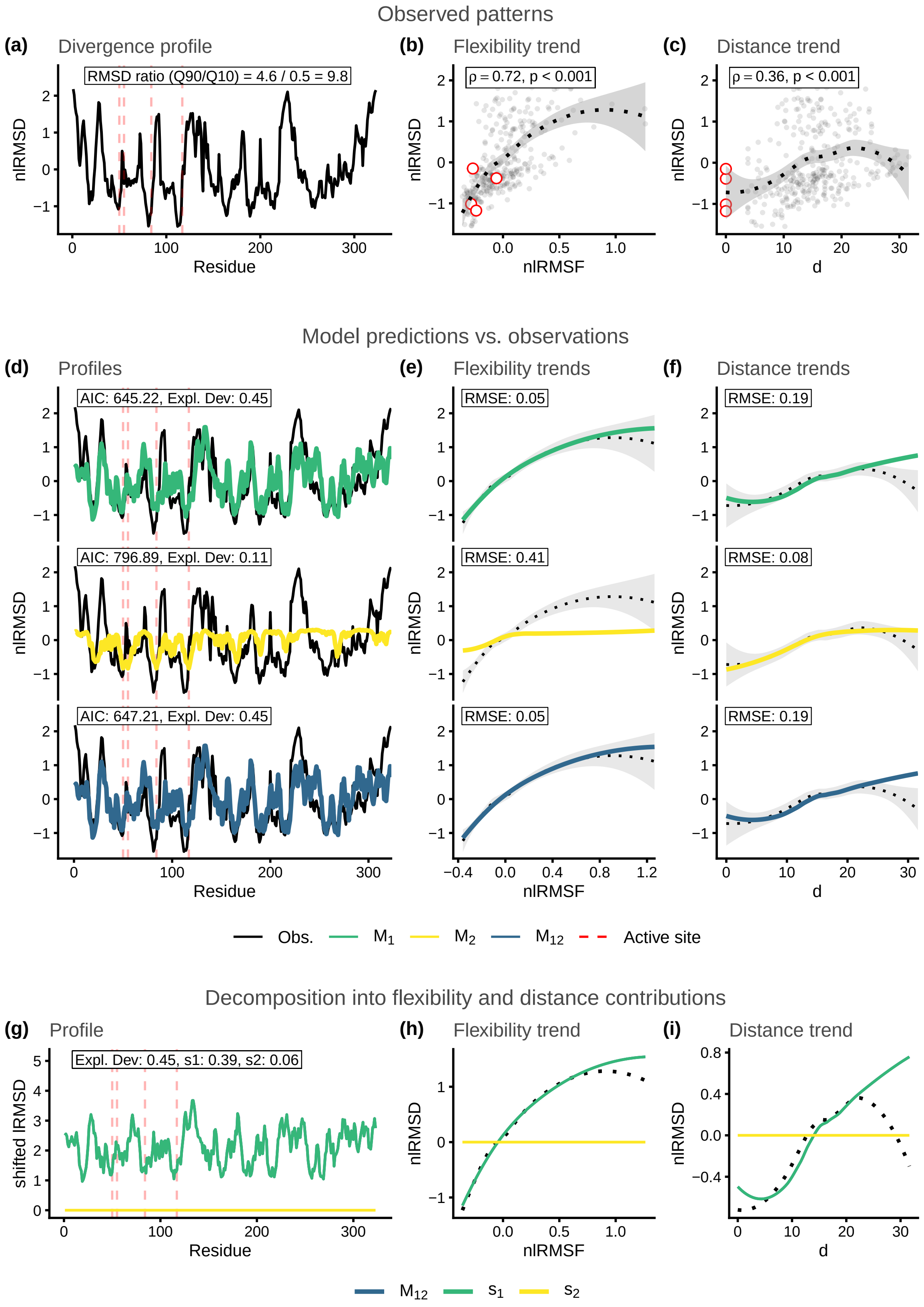

Figure S35: Structural divergence analysis for enzyme family MCSA ID: 858. Reference protein PDB ID: 1mrq_A.

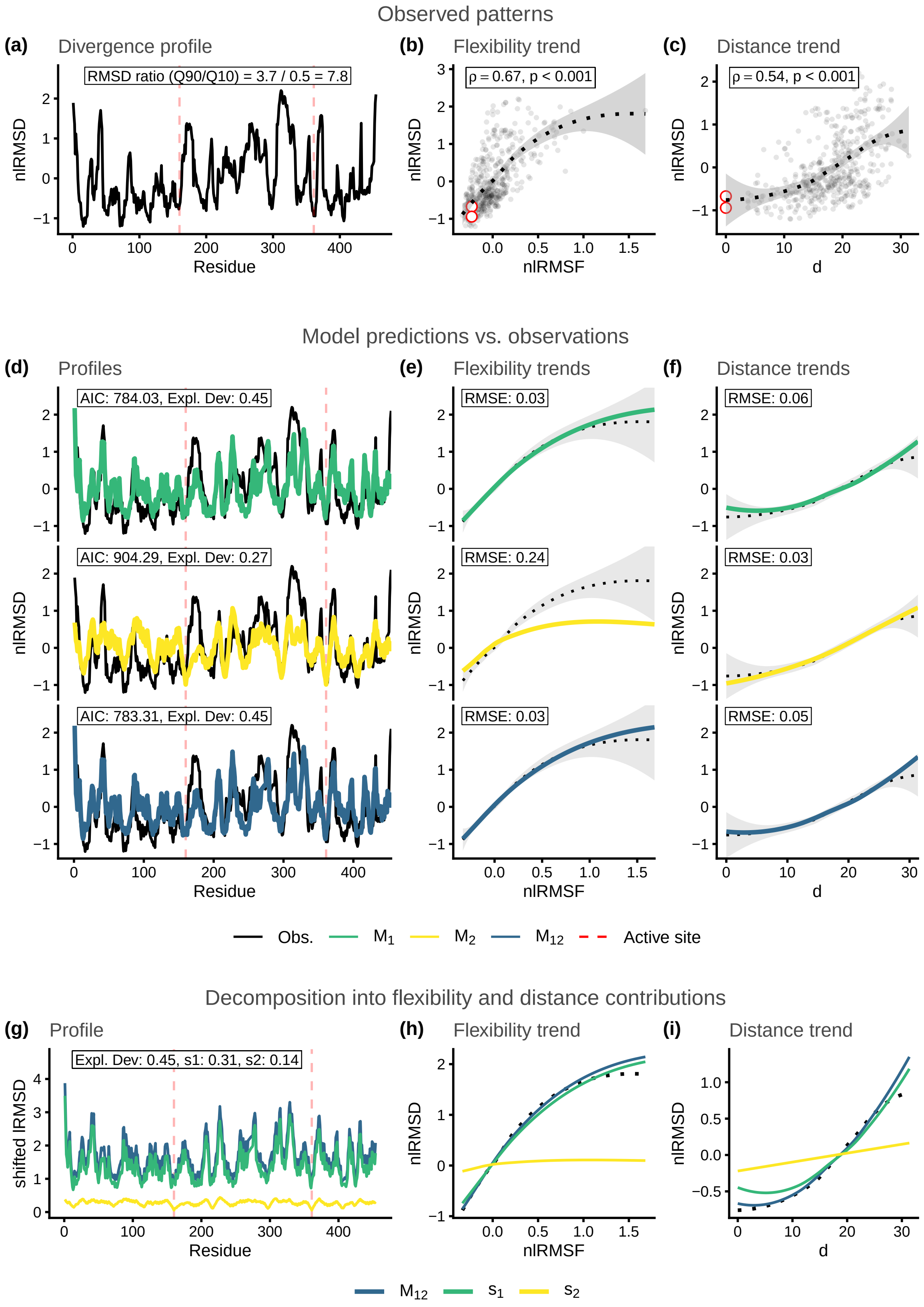

Figure S36: Structural divergence analysis for enzyme family MCSA ID: 877. Reference protein PDB ID: 1pbg_A.

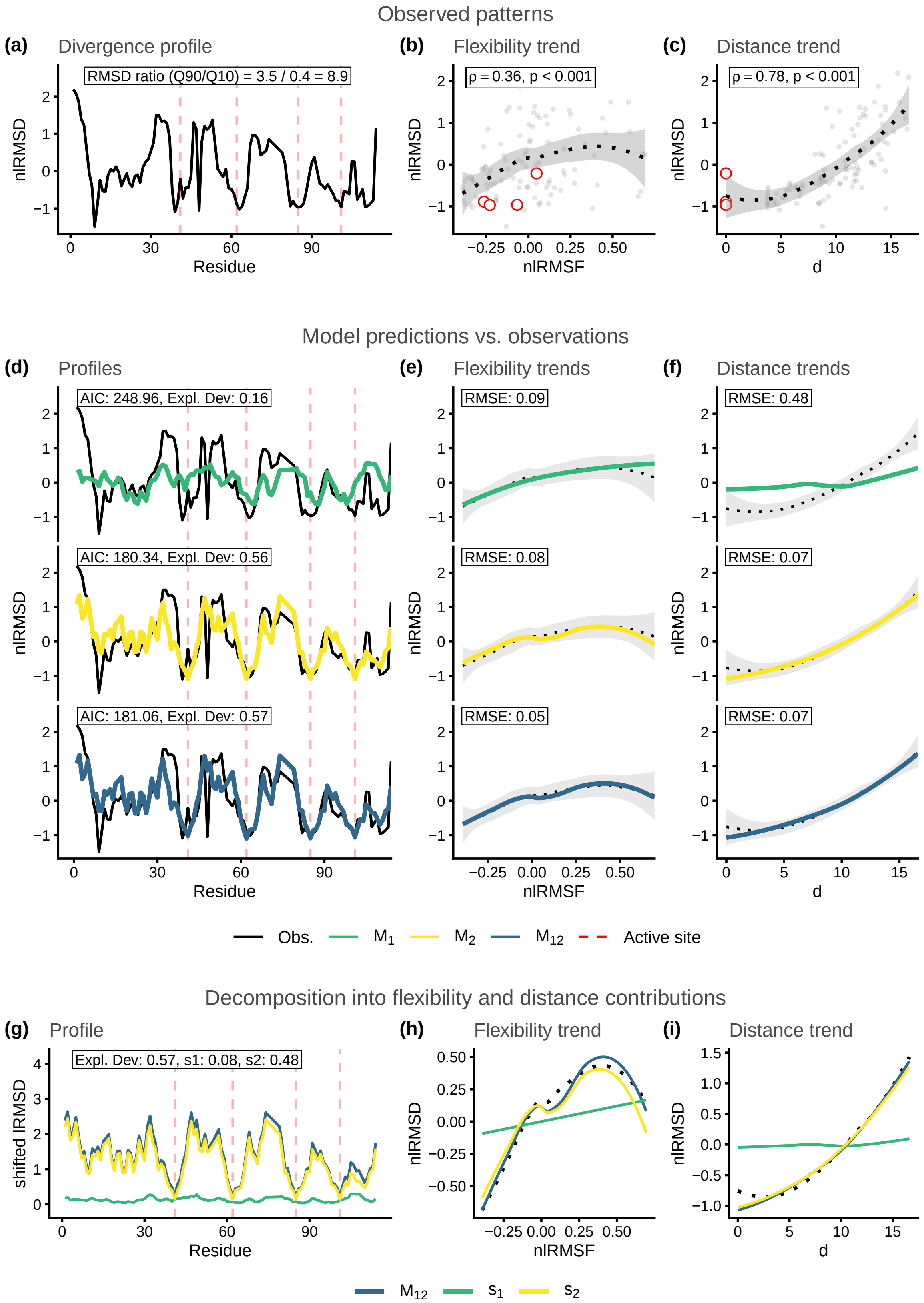

Figure S37: Structural divergence analysis for enzyme family MCSA ID: 908. Reference protein PDB ID: 1rtu_A.

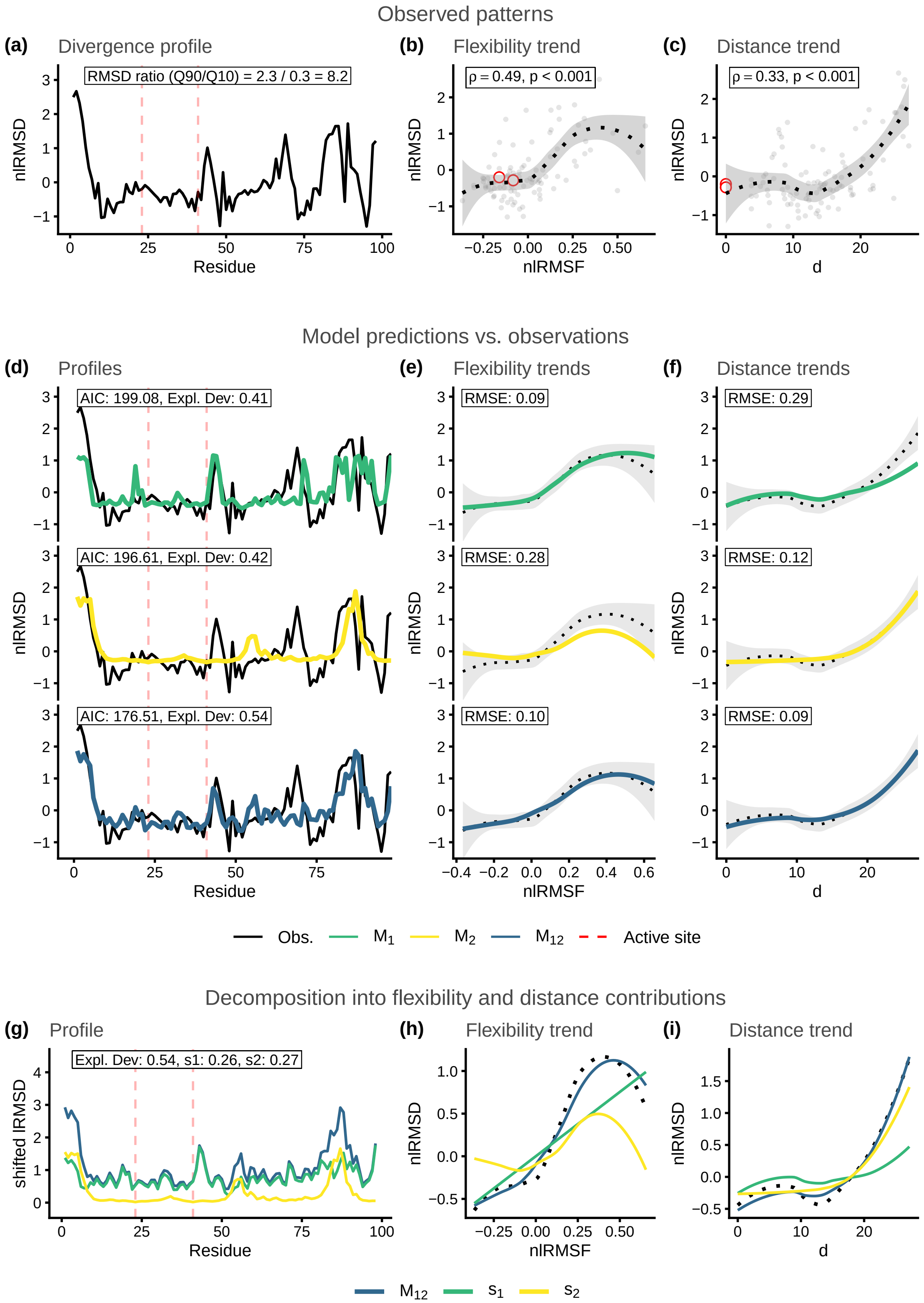

Figure S38: Structural divergence analysis for enzyme family MCSA ID: 923. Reference protein PDB ID: 2acy_A.

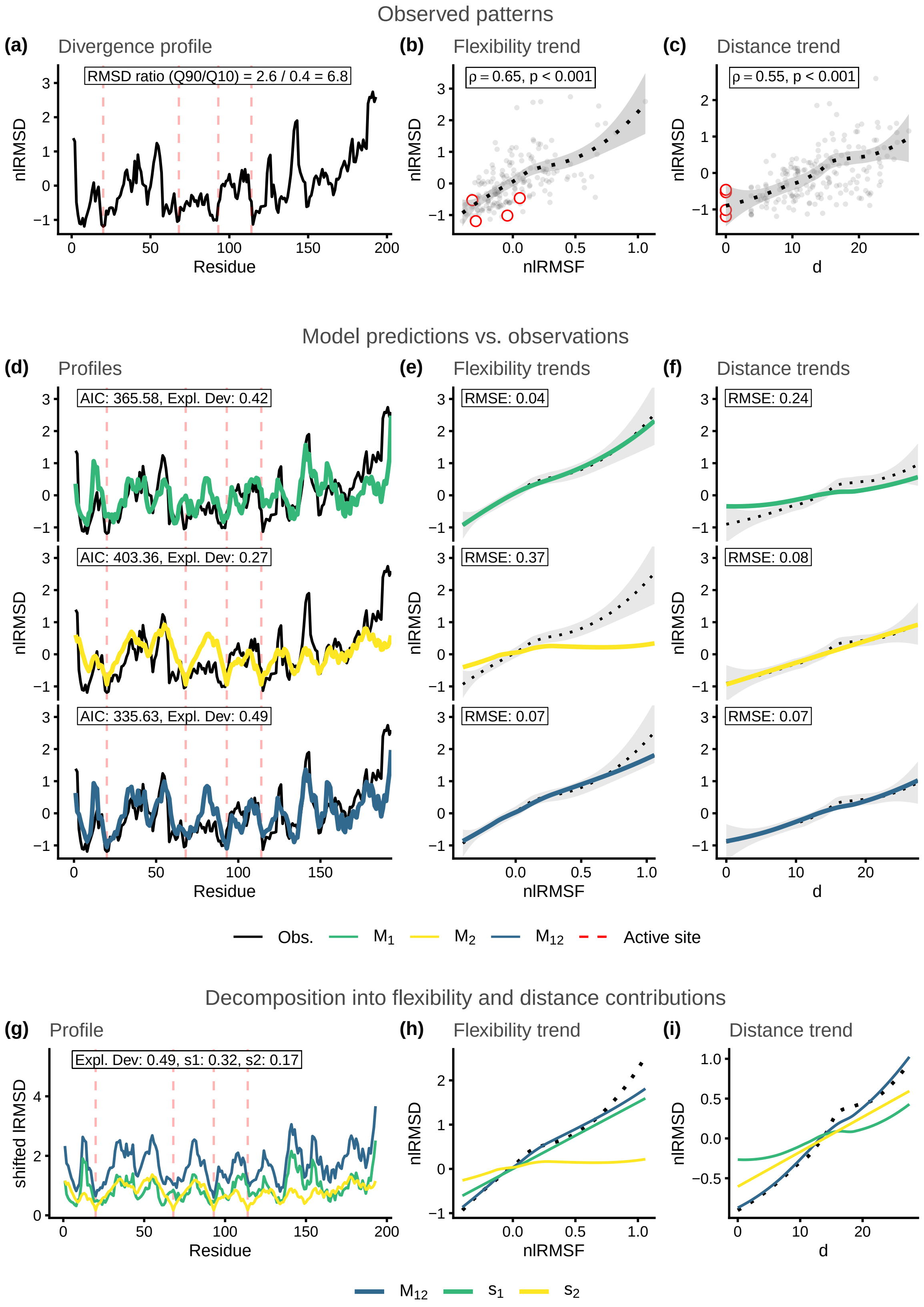

Figure S39: Structural divergence analysis for enzyme family MCSA ID: 931. Reference protein PDB ID: 2pth_A.
